# Polar growth protein Wag31 undergoes changes in homo-oligomeric network topology, and has distinct functions at both cell poles and the septum

**DOI:** 10.1101/2022.04.12.488113

**Authors:** Neda Habibi Arejan, Parthvi Bharatkumar Patel, Samantha Y. Quintanilla, Arash Emami Saleh, Cara C. Boutte

## Abstract

Mycobacterial cell elongation occurs at the cell poles; however, it is not clear how cell wall insertion is restricted to the pole and organized. Wag31 is a pole-localized cytoplasmic protein that is essential for polar growth, but its molecular function has not been described. Wag31 homo-oligomerizes in a network at the poles, but it is not known how the structure of this network affects Wag31 function. In this study we used a protein fragment complementation assay to identify Wag31 residues involved in homo-oligomeric interactions, and found that amino acids all along the length of the protein mediate these interactions. We then used both N-terminal and C-terminal splitGFP fusions to probe Wag31 network topology at different sites in the cell, and found that Wag31 C-terminal-C-terminal interactions predominate at the septa, while C-terminal-C-terminal and C-terminal-N-terminal interactions are found equally at the poles. This suggests the Wag31 network is formed through an ordered series of associations. We then dissected Wag31’s functional roles by phenotyping a series of *wag31* alanine mutants; these data show that Wag31 has separate functions in not only new and old pole elongation, but also inhibition of both septation and new pole elongation. This work establishes new functions for Wag31, and indicates that changes in Wag31 homo-oligomeric network topology may contribute to cell wall regulation in mycobacteria.

**Importance:** Many bacteria restrict cell wall elongation to their cell poles, but it is not known how polar growth is affected on the molecular level. Wag31 is a protein that is required for this polar elongation. In this work, we show that Wag31 actually has at least four distinct functions in regulating the cell wall: it promotes elongation at both poles in different ways, and it can also inhibit cell wall metabolism at the new pole and the septum. In addition, we propose a new model for how Wag31 self-associates into a protein network. This work is important because it shows that a DivIVA homolog can have distinct functions depending on cell context. And, this work clarifies that Wag31 is doing several different things in the cell, and gives us genetic tools to disentangle its functions.

## Introduction

To grow, bacteria need to expand in size by cutting the existing peptidoglycan cell wall and adding new material without disturbing its integrity. Peptidoglycan is critical for maintenance of cell shape, and made of sugar chains crosslinked by small peptides. Many rod shaped organisms perform this cell wall expansion all along the lateral walls (1, 2). Mycobacteria, as well as many other Actinomycetes and some Alphaproteobacteria, restrict cell wall expansion to the cell poles (3, 4). Lateral cell wall expansion in bacteria like *C. crescentus, E. coli*, and *B. subtilis* is controlled MreB, which directs the movement of cell wall enzymes at the site of expansion (2, 5). In Mycobacteria, there are no clear functional homologs of MreB, though the DivIVA domain protein Wag31 is essential for polar elongation (6).

New bacterial cells have a new pole, which is the product of cell division, and an old pole which is inherited intact from the mother. In mycobacteria, the old pole elongates continuously, but there is a delay in initiation of elongation at the new pole (7) which results in asymmetric cells (7, 8) (9). Polar growth is partly regulated by the Intracellular Membrane Domain (IMD), a chemically distinct region of the plasma membrane. The IMD is localized largely to the sub polar regions in growing mycobacterial cells (10). Cell wall precursors are thought to be synthesized and flipped in the IMD region (11). GlfT2 and MurG are IMD-associated enzymes which synthesize precursors for the arabinogalactan and the peptidoglycan layers of the wall, respectively (10).

In this paper we dissect the function of Wag31 (AKA DivIVA, Rv2145c, MSMEG_4217), a mycobacterial homolog of DivIVA. DivIVA proteins are found in Firmicutes and Actinobacteria (Fig. S1). They typically localize to the poles and/or septum, and function to recruit and regulate other factors that have functions in cell division, chromosome segregation and cell wall synthesis (12–16).

In pole-growing *Corynebacterium glutamicum*, DivIVA is essential for polar growth and localizes to the poles (17). DivIVA_Cg_ recruits the essential peptidoglycan synthesizing enzyme RodA to the pole (16) and interacts with PBP1 (13); these interactions are thought to help restrict peptidoglycan synthesis to the pole. In *Streptomyces coelicolor*, DivIVA is involved in polar hyphal growth. It is localized at the hyphal tips and is important for hyphal elongation and splitting(18–21), though remains unclear on the molecular level how DivIVA works.

Wag31 is essential for polar growth in mycobacteria (6, 22), is localized at the poles (23, 24) and at the septum just before division(25). Wag31 helps with chromosome segregation by recruiting the chromosome segregation factor ParA to the poles (26). Wag31 is not required for cell wall metabolism, but it is needed to direct this metabolism to the pole (27). Both RodA and PBP1 in *Msmeg* are delocalized around the entire cell instead of restricted to the poles (27). Thus, *Msmeg* does not regulate polar cell wall synthesis by restricting its essential peptidoglycan transglycosylases to the poles. However, *Msmeg* does restrict synthesis of many cell wall precursor enzymes near the poles, through association with the IMD(10, 27). Depletion of Wag31 destabilizes the IMD (11).

DivIVA proteins are predicted to have coiled coils (23), and are known to form large oligomers (13, 17, 28), which associate the membrane at the cell poles (29, 30). However, the topology of these oligomers is not understood, and is likely to differ between species (Fig. S1). The N-terminal 60-65 amino acid of DivIVA_*Bsub*_ / Wag31 has been crystallized (29, 31). These structures both have two helices involved in a parallel coiled-coil dimer (Fig. 2C). The C-terminus of DivIVA_Bsub_ is thought to form anti-parallel coiled-coils with the C-termini overlapping in a 4-helix bundle (29). The C-terminal region of Wag31 in *Mtb* has low similarity to that of DivIVA_Bsub_, so its structure is unknown. Wag31 is predicted to have coiled coils at its C-terminal helix, and an unstructured loop in the middle (Fig. S1B), which the DivIVA_B.sub_ protein lacks.

**Figure 1.**
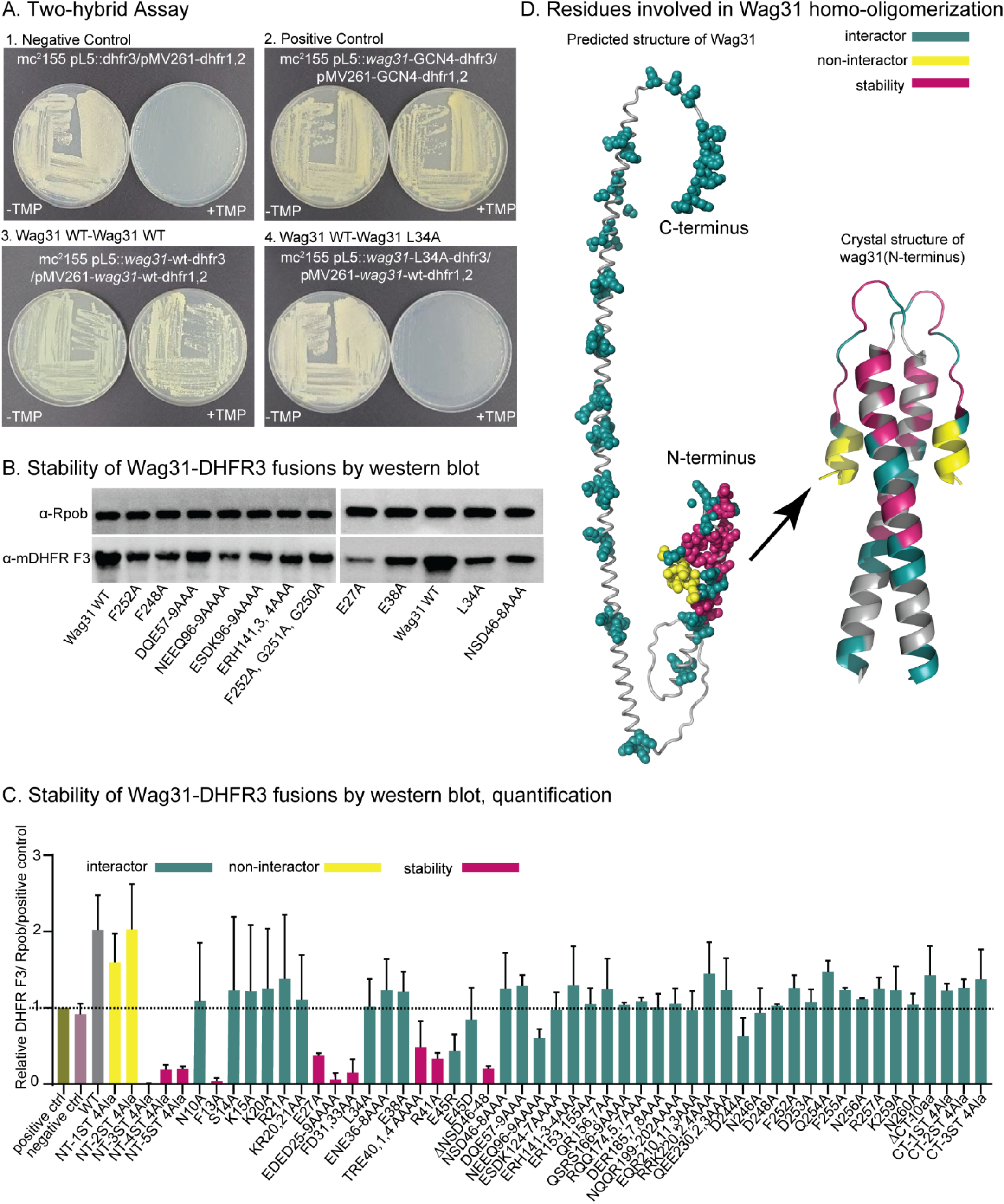
Wag31 has multiple sites of interaction of homo-oligomerization. (A) Representative plates from the Mycobacterial Protein Fragment Complementation two-hybrid assay. Indicated strains were struck out on plates with and without trimethoprim (+TMP/ -TMP). (B) Representative western blots of WT and mutants of Wag31-DHFR3. RpoB serves as a loading control. (C) Normalized quantification of western blot (as in B) band intensities: ratio of ((Wag31-allele/ matched RpoB)/(Wag31 WT/ matched RpoB))/(DHFR-GCN4/ matched RpoB). Values are the mean of western blots from two independent cultures. Error bars represent SD. The dotted line shows the protein level of the DHFR positive control. (D) Alphafold2 predicted structure (Jumper *et al*., 2021) of a Wag31 monomer (left) and crystal structure of a dimer of the N-terminal 60 amino acids of Wag31 (right) (Choukate and Chaudhuri, 2020). The residues are colored according to the results of the two-hybrid assay and the western blots. Teal: mutants of this residue were stable, but failed to interact with WT Wag31 in the two-hybrid. Pink: mutants of this residue were not stable. Yellow: mutants of this residue were stable, and still interacted with Wag31 in the two-hybrid.

**Figure 2.**
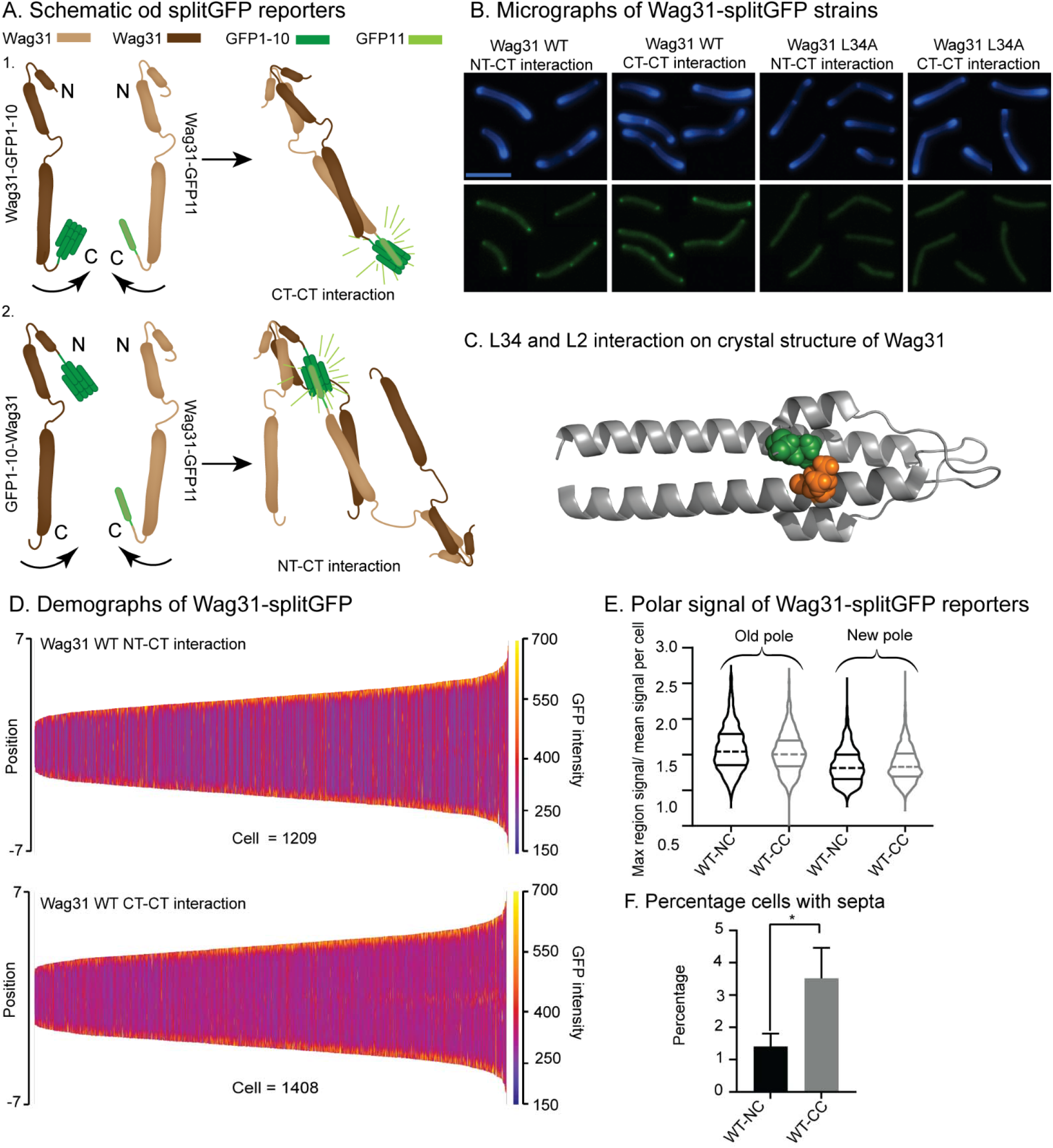
Wag31 homo-oligomeric interaction topology is different at the poles and septum, and L34 is important for initial interactions. (A) Schematic of splitGFP reporter strains. Schematic of control strains can be found in supplemental figure (Fig. S8A). (B) Images of *Msmeg* cells expressing Wag31 WT NT-CT interaction reporter, Wag31 WT CT-CT interaction reporter, Wag31 L34A NT-CT interaction reporter, and Wag31 L34A CT-CT interaction reporter. Top: HADA fluorescence; bottom: splitGFP fluorescence. The scale bar is 5 microns, and applies to all images. (C) Hydrophobic interaction between the L2 of one monomer (green) and L34A (orange) another monomer in the dimer crystal structure of the N-terminus of Wag31. (D) Demographs of splitGFP signal intensity across the length of the cell of the WT splitGFP strains of three biological replicates. At least 300 cells were analyzed from each of three independent biological replicates for each reporter. The cells were pole-sorted based on the HADA intensity, with pole 1, the pole that stains more brightly with HADA, at the top. (E) Quantification of splitGFP polar intensity of cells expressing the Wag31 WT NT-CT (black) and CT-CT (gray) splitGFP reporters. (H) Percentage of cells in (E) that have septal splitGFP signal. *, P = <0.05. *P*-values were calculated by the unpaired two-tailed student’s *t*-test.

In this work we show that much of the Wag31 protein is involved in homo-oligomerization, and that the topology of the Wag31 network seems to develop over the course of the cell cycle. We also identify a series of point mutants of *wag31* that exhibit diverse phenotypes, and indicate that Wag31 has roles in inhibition of new pole elongation and septal inhibition, in addition to its established roles in promoting elongation at the new and old poles.

## Results

### Wag31 has multiple sites of homo-oligomerization

DivIVA homo-oligomerization has been partially characterized in *B. subtilis* (28, 29), and there are separate homo-oligomeric interaction sites on the N and C-terminus of the protein. Functional and sequence differences of mycobacterial Wag31 and DivIVA_Bsub_ led us to ask which parts of Wag31_Mtb_ are involved in homo-oligomerization (Fig. S1).

We used the Mycobacterial Protein Fragment Complementation two-hybrid assay (32) to test a series of Wag31_Mtb_ alanine mutants fused to the mDHFR fragment 3, (mDHFR F3), for interaction with wild-type Wag31_Mtb_ fused to mDHFR F1,2 (Fig. 1A). Interaction of the two Wag31 proteins will reconstitue the split mouse DHFR, yielding trimethoprim resistance in *M. smegmatis*. Almost all of the tested alanine mutants of Wag31_Mtb_ failed to promote trimethoprim resistance, with the exception of eight residues at the very N-terminus (Fig. 1, Table S1A). To determine if these mutant Wag31_Mtb_ proteins are stable, we performed western blots on the two-hybrid strains using α-F3-DHFR antibodies (Fig. 1BC) and found that some of the mutants disrupted stability, but most interrupted homo-oligomerization (Fig. 1D).

These data indicate that many residues involved in homo-oligomerization, thus any single mutation must not completely prevent interaction as other interaction residues are still intact. Since our western blotting shows that these constructs are stable, we conclude that this assay does detect differences in protein associations, but that it has limited dynamic range with Wag31 fusions. Our results indicate that there are likely sites of Wag31_Mtb_ homo-oligomerization beyond just the N and C-terminal interactions described in *B. subtilis* (29). These multiple interaction sites should allow for a complex protein network.

### Wag31 homo-oligomers have differing topology at different sites in the cell

To test whether oligomers of Wag31 have different conformations at the poles and the septa, we used splitGFP probes (Fig. 2A) (33). In this method, GFP1-10 (the first ten beta-strands of superfolderGFP) was fused to either the N-terminus or C-terminus of *wag31*_Msmeg_, and GFP11(final beta strand) was fused to the C-terminus of *wag31*_Msmeg_(Fig. 2A). One strain, called here the NT-CT reporter, (GFP1-10-Wag31 and Wag31-GFP11) will fluoresce when the N-terminus of one Wag31 is near the C-terminus of another. The other strain, called the CT-CT reporter (Wag31-GFP1-10 and Wag31-GFP11) will fluoresce when the C-terminus of two Wag31 are near each other (Fig. 2A). We were unable to test an NT-NT reporter, as the GFP11-Wag31 construct seemed to block polar localization. We performed controls to ensure that the observed signal is not due to splitGFP self-assembly (Fig. S7).

When the fusion constructs were induced, both reporter strains exhibited similar polar fluorescence, which indicates that Wag31 homo-oligomers have both CT-CT and NT-CT interactions (Fig. 2BE). At the septum, the number of cells with septal Wag31 is significantly more in the CT-CT reporter than the NT-CT strain. Because the septum is the site where Wag31 first homo-oligomerizes (Santi *et al*., 2013), our data suggest that newly formed Wag31 structures are more likely to have CT-CT homo-oligomeric interactions. Once the septa become poles, CT-CT and NT-CT interactions are equivalent.

We chose a subset of *wag31* alanine mutants to test in this assay. We used the WT Wag31 NT and CT fusions to GFP1-10, and fused the *wag31* D7A, K20A, L34A, NQQR199-202, and F255A mutants to GFP11 to evaluate whether these mutations alter the topology of Wag31 homo-oligomers. We found that L34A reporter strain resulted in diffuse signal in both NT-CT and CT-CT constructs (Fig. 2B). This leucine forms an apparent hydrophobic interaction with L2 of the other monomer in the crystal structure of the N-terminus (34). The Wag31 L34A protein is largely functional (Fig. 3). These splitGFP data imply that L34A may be required for an early step in homo-oligomerization, and that this mutant protein does not compete well with the WT Wag31 proteins present in this merodiploid strain. SplitGFP data with the other mutants did not show significant, reproducible differences from the wild-type reporters.

**Figure 3.**
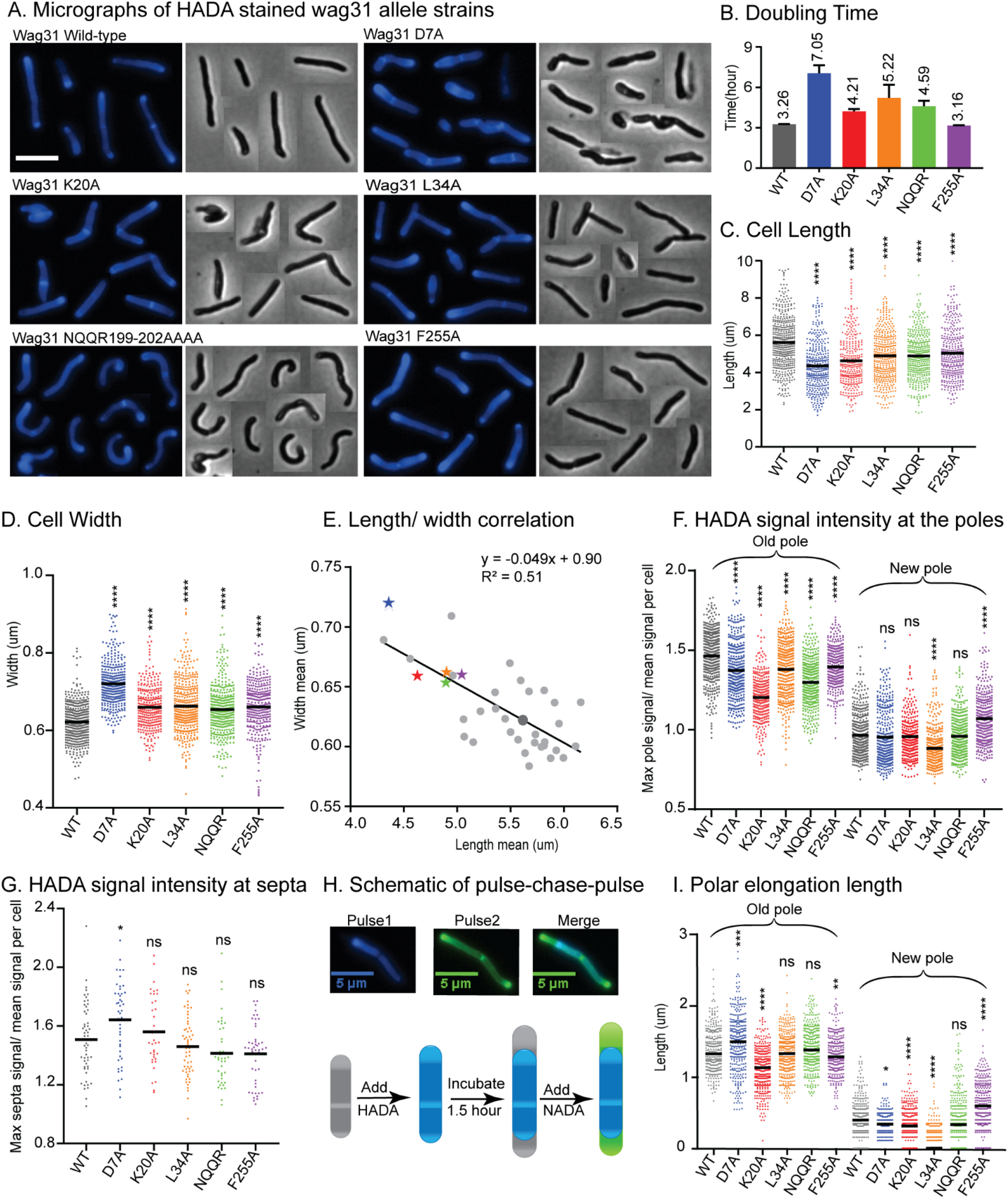
Wag31 has multiple distinct roles in both elongation and division. (A) Phase (right) and fluorescence (left) images of *wag31* allele strains stained with HADA. The scale bar is 5 microns, and it applies to all images. (B) Doubling times of *Msmeg* cells expressing WT or *wag31* alanine mutants. The means (on top of bars) are an average of three biological replicates. Error bars represent SD. (C) Cell lengths of the *wag31* allele strains. Black bars are at the mean. (D) Cell widths of the *wag31* allele strains. Black bars are at the mean. (E) Correlation between cell length and cell width. Black bar is linear regression fit. The five mutants in the other panels are shown as colored stars, the rest of mutants are shown as light gray dots, *wag31* WT is a dark gray dot. (F), (G) Relative polar and septal HADA intensity of *wag31* allele strains. (H) Schematic of pulse-chase-pulse staining method. (I) Length of polar elongation in the *wag31* allele strains, as measured by the pulse-chase-pulse method. Black bar is at the median. ns, p >0.05, *, P = < 0.05, **, P=< 0.005, ***, P =<0.0005, ****, P = <0.0001. All *P*-values are calculated by one-way ANOVA, Dunnett’s multiple comparisons test.

### Wag31 has separate roles in several steps of the cell cycle

Depletion of Wag31 leads to arrest of polar elongation, loss of pole structure, (6, 35) and delocalization of peptidoglycan and mycomembrane metabolism (22). To dissect Wag31’s functional roles in these processes, we profiled the phenotypes of *Msmeg* strains with *wag31* WT replaced by the stable alanine mutants of *wag31*, using L5 allele swapping (36) (Fig. 1BC). We were unable to swap some of the mutants, thus P2, T4, K15, and EQR 210-3 are essential (Fig S2C).

We phenotypically profiled the *wag31* allele swap strains (Fig 3A-I). Most of the mutants at the N-terminus of Wag31 impair growth rate, while most mutants in the middle and C-terminus of the protein do not (Fig. 3B, S2A). Microscopy experiments many of the mutants are short and/ or wide (Fig. 3ABCDE, S2). Most of the *wag31*_Mtb_ mutant strains have higher HADA signal, which is a proxy for peptidoglycan metabolism, than *wag31*_Mtb_ WT strain (Fig. S3A, S5, S6), possibly because of increased permeability. To compare localized peptidoglycan between strains, we used relative intensity at the poles and septa. HADA stains the old pole more brightly (37); throughout we assume that the brighter pole is the old pole.

Most mutants had lower relative HADA signal than WT at the old pole, and none have higher signal (Fig. 3F, S3B). At the new pole most mutants have the similar signal to WT, but a few near the N-terminus are dimmer than WT, and a few near the C-terminus are brighter than WT (Fig. 3F, S3C). This data indicates that mutants in Wag31 can have different impacts on the peptidoglycan metabolism at the two cell poles. Septal HADA intensity is increased in a few mutants (D7A, K20A, DER 185,7,8), while it is decreased in others (Fig. S4B), suggesting that Wag31 regulates peptidoglycan synthesis at the septa as well. We also observe that the septal placement is shifted toward either pole in some of the mutants (Fig. S4C).

A variety of different phenotypes indicative of assorted functional roles for Wag31 were apparent across all the mutants. We chose five mutants that represented the range of observed phenotypes for further study: D7A, K20A, L34A, NQQR199-202AAAA, and F255A.

### Wag31 is involved in septal inhibition via glutamate 7

As the first eight residues of Wag31 are not required for homo-oligomerization (Fig. 1). We chose one mutant in this region for further study, D7A. The *wag31* D7A strain grows slowly, and the cells are short and wide (Fig. 3BCD). There is a slight decrease in HADA intensity, compared to WT, at both poles (Fig. 3F), but HADA intensity is significantly increased at the septa (Fig. 3G).

To determine whether HADA staining at the poles is indicative of polar elongation or peptidoglycan remodeling(37), we performed pulse-chase-pulse staining with two colors of fluorescent D-amino acids. We stained first with HADA, then outgrew the cells without stain for 1.5 hours, then stained with the green fluorescent D-alanine NADA, and measured the length of the poles that are green and not blue (Fig. 3H). Results from this experiment show that the *wag31* D7A strain elongates more than the WT at the old pole, and elongation at the new pole is similar to WT (Fig. 3I).

Our data show that the *wag31* D7A mutant is not defective for elongation (Fig. 3I); however, the cell length is substantially decreased (Fig. 3C), suggesting that division activated at shorter cell lengths. The higher septal HADA staining (Fig. 3G, S4B) suggests that there is more peptidoglycan metabolism at the septum than in WT cells. This cell division may sometimes be too early, which could lead to the unusual number of ghosts in the *wag31* D7A phase images (Fig. 3A). These data suggest that Wag31 is involved in inhibition of septation through D7. As D7A is not involved in homo-oligomerization (Fig. 1), the effects of this mutation could be due to an inability to interact with other proteins or the cell membrane. We localized Wag31 D7A-GFPmut3 in a merodiploid strain and find that it localizes like the Wag31 WT-GFPmut3 at the poles and septum (Fig. 4).

**Figure 4.**
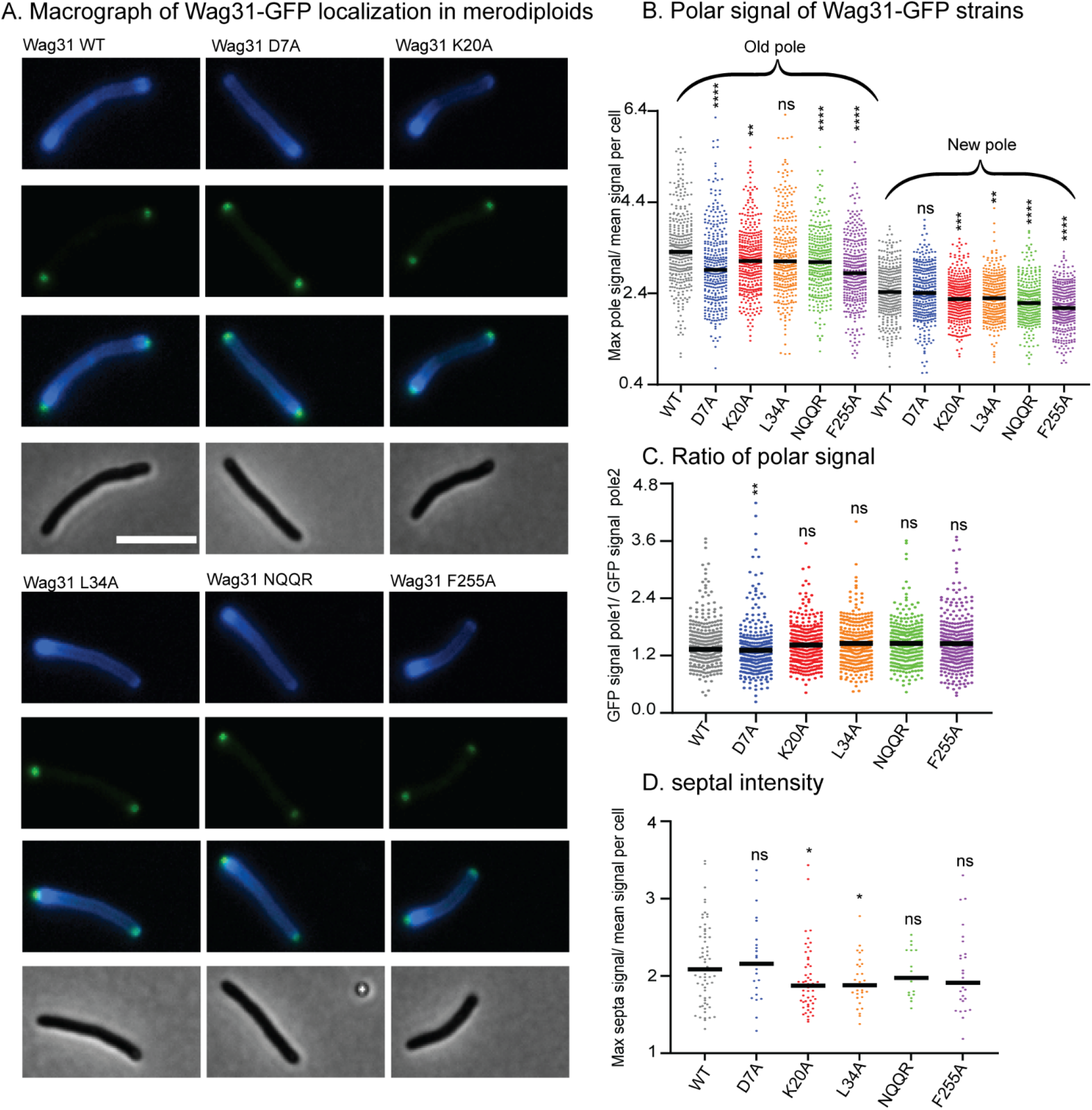
Wag31 mutants localize normally in merodiploid strains. (A) Micrographs of WT *Msmeg* expressing Wag31-GFPmut3 constructs with the indicated mutations. Top: HADA; second: GFP; third: merged; bottom: phase. The scale bar is 5 microns, and it applies to all images. (B) Relative polar intensity of GFPmut3 signal from cells in A. (C) Ratio of normalized signal at the old pole over normalized signal at the new pole, from B. (D) Relative septal intensity of GFPmut3 signal from cells in A. ns, P >0.05, *, P =< 0.05, **, P=< 0.005, ***, P=< 0.0005, ****, P= <0.0001. All *P*-values (B), (C) are calculated by one-way ANOVA, Dunnett’s multiple comparisons test and the *P*-value (D) is calculated by the Welch’s t-test.

### Wag31 regulates polar elongation through N-terminal residue K20

The N-terminus of Wag31 is highly conserved (Fig. S1). This part of the protein dimerizes and has been structurally characterized (29, 34). In *Bacillus subtilis*, F17 is involved in interaction with the membrane at the cell pole (29). In *Msmeg*, K20 is equivalent to F17 in DivIVA_*Bsub*_.

Our microscopy data shows that the *wag31* K20A mutant has decreased HADA staining compared to WT at the old pole, but equivalent staining at the new pole (Fig. 3F). To determine whether the HADA staining difference represents a difference in elongation, we performed the pulse-chase-pulse experiment (Fig. 3H). The results show that the *Wag31* K20A mutant elongates ∼.26 mm less than the WT at the old pole, and ∼.14 mm less at the new pole (Fig. 3FHI). The defect in *wag31* K20A elongation (22% less) is similar to the defect in cell length (18%), indicating that this mutant is defective in elongation, especially at the old pole. The *wag31* K20A strain has similar septal intensity as the WT (Fig 3G), implying that peptidoglycan metabolism there is comparable; however, the septal location is shifted slightly toward the old pole (Fig. S4C), which could be explained because the old pole is elongating more slowly.

To determine if K20 is involved in membrane association, we localized K20A-GFPmut3 in a merodiploid strain, because the allele swaps were not viable. The Wag31 first associates with the membrane at the septum. Localization of Wag31-GFPmut3 to the septum reflects its interaction with the membrane, whereas localization at the poles reflects interaction with other Wag31 proteins. The Wag31 K20A-GFPmut3 protein does not show a defect in polar localization indicating that K20 residue alone is not necessary for polar localization. But it is defective in septal localization (Fig. 4BCD), indicating that the Wag31 K20A mutant may be slightly impaired in its ability to adhere to the polar membranes. Residue K20 promotes Wag31 homo-oligomerization (Fig. 1ACD). The polar growth phenotypes in the *wag31* K20A mutant could be due to defects in an unknown hetero-oligomeric interaction, or defects in the structure of the homo-oligomeric network which could affect membrane structure or dynamics at the pole, or affect the activity or localization of other proteins.

### Wag31 regulates new pole elongation through L34

We found that the *wag31* L34A mutant also has a defect in polar elongation. The *wag31* L34A mutant has decreased HADA staining at both poles, the old pole staining is 1.5% less, and the new pole is ∼11% less than the WT (Fig. 3F). The old pole of the *wag31* L34A strain elongates the same amount as the WT strain, but the new pole has ∼24% less elongation (Fig. 3I). In the population of *wag31* L34A mutants that we characterized, ∼65% had no observable elongation at the new pole, while only 11% of cells in the *wag31* WT population had no new pole elongation in the 1.5 hour assay. Thus, it appears that the Wag31 L34 residue contributes to both initiation and extension of the new pole. The short cell length and slow growth rate (Fig. 3ABC) are likely due to this defect in new pole elongation, as we did not observe defects in the intensity or location of septa (Fig. 3G, S4C). We localized the Wag31 L34A-GFPmut3 in a merodiploid, and it does not have defects in polar localization. It is slightly dimmer than the Wag31 WT-GFPmut3 at the septum (Fig. 4D), which is the site of new pole construction.

### Wag31 inhibits new pole initiation through F255

We made several alanine mutants in the C-terminus of Wag31, which is not part of the conserved DivIVA domain (Fig. S1). We found that in the *wag31* F255A mutant, there is increased intensity of HADA at the new pole, and less at the old pole (Fig.3F). The pulse-chase-pulse experiment shows that the *wag31* F255A strain elongates ∼47% more than the WT at the new pole, while the old pole elongates 5% less compared to the WT strain (Fig. 3I). There is no difference in polar and septal localization between the Wag31 F255A-GFPmut3 and the Wag31 WT-GFPmut3 (Fig. 4). The HADA septal intensity is similar in the *wag31* F255A mutant compared to the WT (Fig. 3G), while septal location is shifted toward to the old pole, which could result from the increased elongation activity at the new pole (Fig. S4C). The *wag31* F255A grows as fast as the WT strain (Fig. 3B), and the cells are slightly shorter and thicker than the WT (Fig. 3CDE). The apparently increased elongation may be balanced by increased septation to result in shorter cells. These data indicate that Wag31 has inhibitory role on elongation of the new pole through its C-terminal residues (Fig. S3). So, Wag31 can both promote and inhibit polar growth.

### Maintenance of polar integrity is controlled by NQQR199-202 residues

Wag31 is necessary for rod shape (6, 22, 23). Forming a straight rod requires that new cell wall material be added evenly around the circumference of the cell. Previous work showed cell bending when Wag31-eGFP replaces native Wag31 (24), suggesting that deforming the Wag31 network may lead to uneven cell wall insertion. We observed a similar curved cell morphology in the *wag31* NQQR199-202AAAA mutant (Fig. 3A). While the *wag31* NQQR199-202AAAA mutant has slightly decreased HADA intensity at the old pole (Fig. 3F), pulse-chase-pulse staining shows that that elongation is similar to the Wag31 WT at both poles (Fig. 3I). There are no differences in localization of Wag31 NQQR199-202AAAA-GFPmut3 and Wag31 WT-GFPmut3 (Fig. 4). The NQQR199-202 residues are toward the middle and involved in Wag31 homo-oligomerization (Fig. 1). We propose that the Wag31 NQQR199-202AAAA mutation alters the structure of the oligomeric network to cause cell wall insertion to be uneven around the circumference of the cell, leading to slight polar bulging and curved poles (Fig. 3A).

### Wag31’s effects on MurG and GlfT2 localization do not correlate with effects on polar growth

Recent work suggests that localization of the enzymes that synthesize cell wall precursors to the subpolar Intracellular Membrane Domain (IMD) may be critical in restricting elongation to the pole (10, 27). Depletion of Wag31 has been shown to delocalize the IMD(11), though it is not known if Wag31 regulates the IMD, or if the IMD localization is merely dependent on an intact cell pole.

To probe whether Wag31 regulates IMD structure, we localized both GlfT2-mcherry2B and MurG-Venus in selected Wag31 alanine mutants. GlfT2 is the last galactan cytoplasmic enzyme in arabinogalactan synthesis (38). GlfT2 associates with the IMD, and localizes in the typical pattern of IMD-associated proteins (10). MurG is the final cytoplasmic enzyme in peptidoglycan precursor synthesis, has a similar localization pattern as GlfT2, (24) and is found in both cytoplasmic and IMD fractions (11).

Our microscopy data show that localization of GlfT2-mcherry2B is significantly decreased at the old pole in the *wag31* K20A, D7A, L34A and F255A mutants compared to the WT (Fig. 5ABC), while it was unchanged at the new pole (Fig 5ABD). Localization of MurG-Venus is slightly decreased at the old pole in the *wag31* D7A, L34A and NQQR199-202 mutants compared to the WT (Fig. 5E), and largely unaffected at the new pole (Fig. 5F). Across the mutants tested, polar elongation (Fig. 3HI) does not correlate either with GlfT2 or with MurG localization signals at the poles (Fig. 5). These data suggest that Wag31 may regulate the localization of IMD proteins, probably indirectly and largely at the old pole, but that this regulation does not directly control polar elongation.

**Figure 5.**
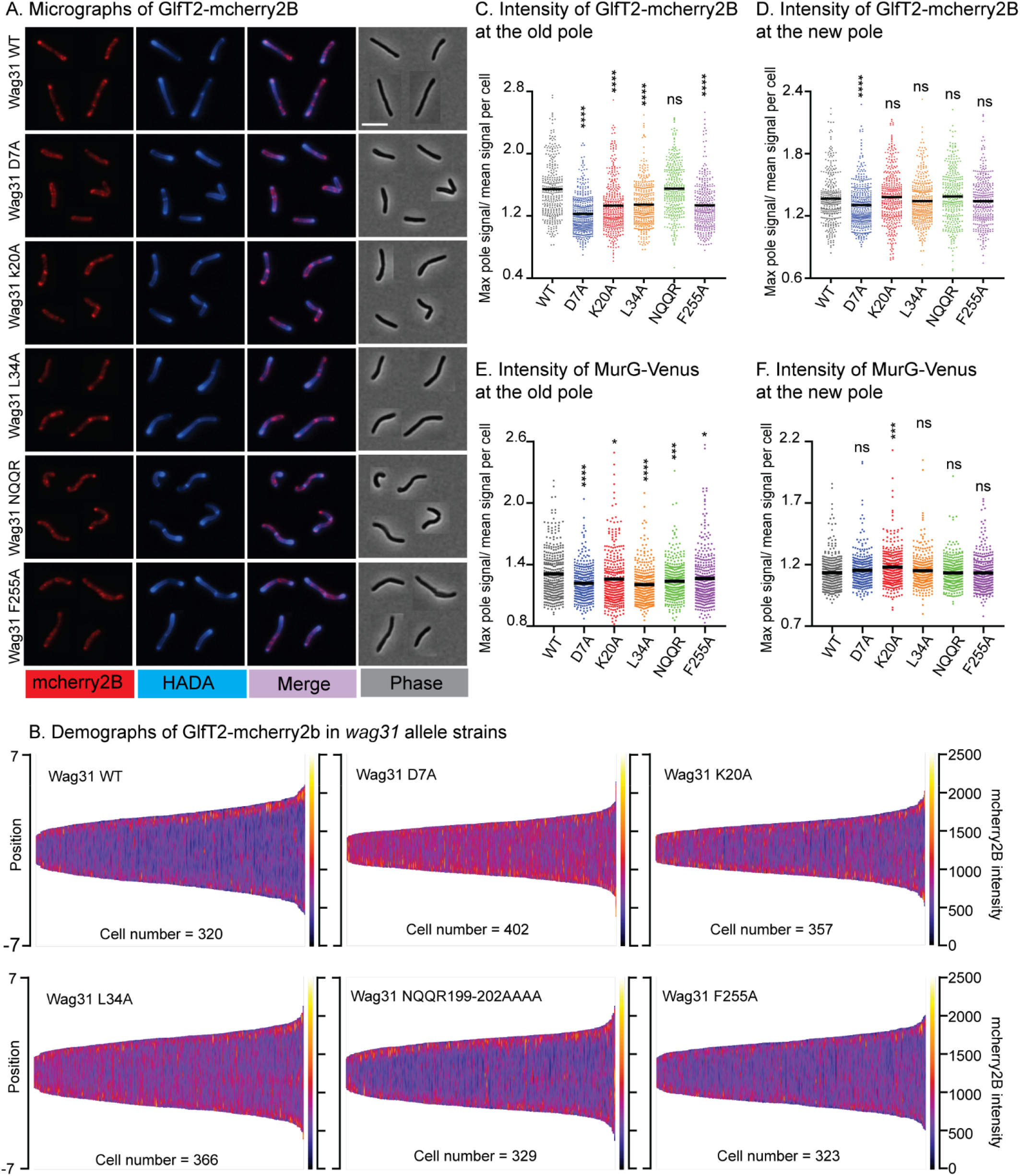
Localization of IMD proteins does not determine changes in polar growth. (A) Micrographs of *wag31* allele strains expressing GlfT2-mcherry2b and stained with HADA. The scale bar is 5 microns, and it applies to all images. (B) Demographs of GlfT2-mcherry2b intensity across the length of each cell (Y axis of each plot, with intensity indicated by color – scale to the right of each plot), arranged by cell length (along the X axis). At least 100 cells were analyzed for each of three biological replicate cultures. (C) and (D) Polar intensity of GlfT2-mcherry2b at the old pole and the new pole, respectively, from cells in A. (E) and (F) Polar intensity of MurG-Venus at the old pole and the new pole, respectively. ns, P >0.05, ***, P=< 0.0005, ****, P= <0.0001. *P*-value is calculated by one-way ANOVA, Dunnett’s multiple comparisons test.

## Discussion

Wag31’s molecular function in regulating the cell wall has remained obscure. Homology with other systems predicts that Wag31’s would recruit and regulate cell wall enzymes at the pole (12, 16). However, the heterooligomeric interactions that have been described for Wag31 – AccA3 (24, 39) and FtsI (40) - do not connect to Wag31’s role in restricting cell wall metabolism to the poles. Additionally, these interactions may be indirect (AccA3,) or only exist under certain stress conditions (FtsI). The other interaction that Wag31 makes is with itself. Here, we sought to probe the assembly of the homo-oligomeric network, and to dissect Wag31’s function in regulating the cell wall (Fig. 6).

**Figure 6.**
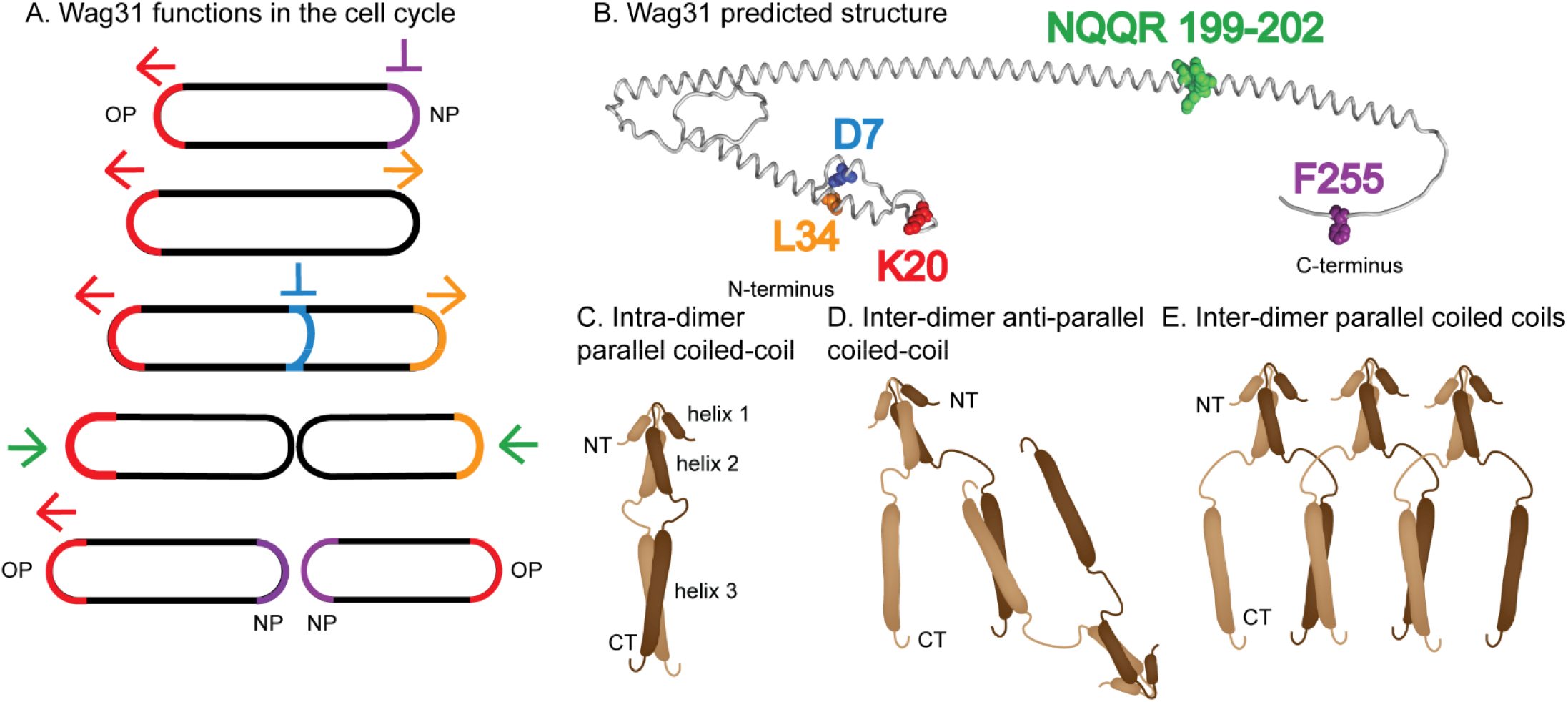
Models. (A) Multiple functions affected by Wag31 are highlighted in different steps of the cell cycle. Red = old pole elongation; purple = inhibition of new pole elongation; orange = new pole elongation; blue = inhibition of septation; green = maintenance of pole structure. (B) The residues shows to affect the functions in (A) are highlighted on an Alphafold2 predicted structure (Jumper *et al*., 2021) of a Wag31 monomer. (C) Model of a Wag31 dimer showing the N-terminal coiled coil of the helices 1 and 2 from the crystal structure (Choukate and Chaudhuri, 2020), and a intra-dimer parallel coiled coil of helix 3. (D) Model of a portion of Wag31 network showing an inter-dimer anti-parallel coiled coil of helix 3, with intra-dimer N-terminal interactions as in (C). (E) Model of a portion of Wag31 network showing an inter-dimer parallel coiled coil of helix 3, with intra-dimer N-terminal interactions as in (C).

### Predictions about Wag31 network structure

The predicted structure of DivIVA in *B. subtilis* has only two alpha-helices which are thought to be arranged end-to-end, forming a chain (29). In Wag31 in mycobacteria, the predicted structure has three helices, and a 33 amino acid loop between the helices 2 and 3 (Fig 6C). We predict that the long loop will allow more flexibility in the topology of the network and would likely allow branching of the network as helices 1 and 2 of a monomer could interact with different monomers than helix 3 (*e*.*g*., Fig 6C vs Fig. 6DE). Wag31 is predicted to form coiled coils at its helices through hydrophobic interfaces. The hydrophobicity map of the predicted Wag31 Alphafold2 structure (41) shows a hydrophobic interface that forms the coiled coil of the N-terminal region (helices 1 and 2) in the crystal structure. Helix 3 also has a stripe of hydrophobic residues, which is likely the site of a double or triple coiled coil (42)(Fig. S1B). Helix 3 in Wag31 might form either a parallel or anti-parallel coiled coil with the helix 3 in another monomer (Fig. 6CDE). A parallel coiled coil would bring the C-terminal of two monomers near each other, while the anti-parallel coiled coil might bring the N-terminus of one monomer near the C-terminus of another.

We made one leucine to alanine mutation in helix 2, and this disrupted homo-oligomerization in both the two-hybrid and split-GFP assays (Fig. 1,2). The inability of this Wag31 L34A mutant to interact with Wag31 WT in the split-GFP assay, where it is competing for interaction with WT protein, indicates that this L34 residue may be important for an initial dimerization event. We therefore propose that these hydrophobic coiled-coil interactions may be the first to form as Wag31 monomers associate. We also tested how numerous charged and polar residues along the length of the Wag31 protein affect homo-oligomerization (Fig. 1). We found that, with the exception a few residues at the very N-terminus, all the residues we tested contributed to Wag31 homo-oligomerization in the sensitive two-hybrid assay. We propose that the charged residues on the surface of the coiled-coil structure may mediate further salt-bridge interactions that help to expand the network via stacking of coiled-coils.

We used splitGFP reporters to measure the relative proportion of CT-CT interactions, and NT-CT interactions. The CT-CT signal might result from parallel coiled-coils forming on helix 3, while the NT-CT signal could result from either anti-parallel coiled-coils at helix 3, or from stacking of chains of coiled-coils through the proposed salt bridges (Fig. 6CDE). We found that at the septum, where new Wag31 first interacts with the membrane, CT-CT interactions predominate. This supports the idea that parallel coiled coils form from helix 3 of neighboring dimers, and that these interactions form early in the construction of the network (Fig. 6E). We expect that Wag31, like DivIVA in close relative *C. glutamicum* (30), will diffuse very little once incorporated into the network. Thus, new coiled-coil Wag31 chains likely accumulate at the poles in layers, where CT-CT and NT-CT interactions are evenly balanced (Fig. 2). It is likely that other proteins interact with the Wag31 network and alter its topology and structure (30).

### Dissection of Wag31 functions

Wag31 had previously been known to be essential for polar growth (6, 27). Here, we phenotypically profiled alanine mutants of residues all along the length Wag31 and found a variety of different phenotypes which indicate that Wag31 has several distinct functions in cell growth (Fig. 6AB). We found that the *wag31* K20A mutant is specifically more defective in old pole elongation while the *wag31* L34A mutant is specifically more defective in new pole elongation (Fig. 3). These results indicate that Wag31 is doing something different at the new and old poles. The structure of the Wag31 homo-oligomeric network could be different at the two poles, and these mutations could interfere with the network structure. Or, these mutations might interfere with interactions of Wag31 with other proteins that regulate cell wall synthesis.

Our characterization of the *wag31* D7A mutant indicates that Wag31 is involved in downregulating septation. Wag31 had previously been shown to localize to the cell division site before septation (25), and overexpression of Wag31-GFP was shown to inhibit and mis-localize septa (23). D7 is not required for homo-oligomerization, so it is likely involved a hetero-oligomeric interaction. Wag31 interacts with the septal PBP FtsI and regulates FtsI stability under oxidative stress through residues 46-48 (NSD) (40). We tested a *wag31* Δ46-48 mutant, as per (40) and found it was destabilized (Fig. 1C). We also made a *wag31* NSD46-48AAA mutant, and found that it had normal cell length (Fig. S2), so those residues do not affect FtsI activity in log. phase. The Wag31 D7 residue could possibly form a separate interaction with FtsI, or other septal factors (43).

We found that the *wag31* F255A mutant had increased polar elongation at the new pole (Fig. 3). This suggests that Wag31 has a role in inhibiting new pole elongation. The slower elongation at the new pole is partly controlled by membrane protein LamA (8). Our work suggests that either Wag31 works separately to inhibit new pole elongation, or that LamA may interact with Wag31 through F255 and interfere with Wag31’s promotion of new pole elongation.

Most of the mutants we characterized show a defect in GlfT2-mcherry2B and MurG-Venus localizations at the old pole, but not at the new pole. The localization of GlfT2 and MurG at the poles did not correlate with polar elongation defects (Fig. 3,5), indicating that polar localization of the IMD proteins is not the only determinant of polar growth. These data suggest that Wag31 is at best an indirect regulator of the IMD, and that regulation of the IMD is likely not Wag31’s essential function.

In this work, we describe a new model for the assembly of the Wag31 homo-oligomeric network (Fig 6). Many questions remain about the structure of this network, but our work indicates that there are likely numerous homo-oligomeric interfaces and that the structure of the network likely develops as a septal site turns into a pole (Fig. 2). We also define new roles for Wag31 in regulating both septation and the inhibition of the new pole, and identify residues that have dominant roles in Wag31’s different functions (Fig 6).

## Materials and Methods

### Bacterial strains and culture conditions

All *Msmeg* strains were grown in liquid culture in 7H9 (BD, Sparks, MD) medium supplemented with 0.2% glycerol, 0.05%Tween 80, and ADC (5g/L albumin, 2 g/L dextrose, 0.85 g/L NaCl, 0.003 g/L catalase). For most experiments, *Msmeg* strains were grown on LB Lennox plates. DH5a, TOP10, and XL1-Blue *E*.*coli* cells were used for cloning. For *E. coli*, antibiotics concentrations were: kanamycin – 50 <u>μ</u>g/ml; hygromycin - 100 <u>μ</u>g/ml; nourseothricin - 40 <u>μ</u>g/ml. For *Msmeg*, antibiotic concentrations were: kanamycin - 25 <u>μ</u>g/ml; hygromycin - 50 <u>μ</u>g/ml; nourseothricin – 20 <u>μ</u>g/ml; trimethoprim - 50 <u>μ</u>g/ml.

### Growth curves

Strains were grown to log. phase in 7H9 with appropriate antibiotics, then diluted to OD600 = 0.1 without antibiotics in a 96 well plate. A Synergy Neo2 linear Multi-Mode Reader was used to shake the plates continuously for 18 hours at 37°C, and read OD every 15 min. To find the doubling time for each strain, the raw data were analyzed with exponential growth equation model using GraphPad Prism (version 9.1.2).

### Strain construction

To build the allele replacement strains, first *wag31*_Msm_ under its native promotor was cloned into a kanamycin-marked L5 integrating vector and transformed into *Msmeg* mc^2^155. Then, *wag31* at its native locus was replaced with hygromycin resistance cassette by dsDNA recombineering (44). This cassette was oriented so the hygR promoter drives expression of downstream genes in the operon. Deletion of *wag31* from the genome was confirmed with PCR checks. *wag31*_Msm_ at the L5 site was swapped with *wag31*_Mtb_ alleles in a nourseothricin-marked L5-integrating vector using L5-phage site allelic exchange as described (36). The transformants were screened by antibiotic resistance in order to confirm allele swap. Vectors containing *wag31*_Mtb_ mutants were created using PCR stitching. All vectors were made using Gibson cloning (45). All strains, plasmids and primers used are listed in supplemental material part (Table S2A, S2B, S2C).

### Mycobacterial protein fragment complementation (M-PFC) assay

*Msmeg* strains carrying Mycobacterial protein fragment complementation (M-PFC) assay vectors were grown to log. phase and struck on 7H11 agar (BD Difco™, Thermo Fisher Scientific) with or without trimethoprim at 50 <u>μ</u>g/ml as described (32). Plates were incubated at 37°C for 4 days, and then photographed.

### Immunoblot analysis

*Msmeg* strains carrying M-PFC vectors were grown to log. phase (OD600 = 0.5-0.7), pelleted, and lysed by bead beating (Disruptor Beads 0.1mm from Disruptor Genie®, Mini-Beadbeater-16, Biospect). Supernatants were separated by SDS-PAGE and transferred onto polyvinylidene difluoride (PVDF) membranes (GE Healthcare) and blocked with BSA for 1 hour. The blot was probed with a-C-terminal-DHFR rabbit antibody (1 :1000 Sigma-Aldrich, D0942) and goat anti-rabbit IgG HRP-conjugated secondary antibody (1:10,000, Thermo Fisher Scientific 31460) in PBST. a-Rpob was used as loading control (10 :10000, Thermo Fisher Scientific, MA1-25425). All bands were quantified using Fiji and normalized to the controls.

### Microscopy

All microscopy was performed on three biological replicate cultures of each strain using a Nikon Ti-2 widefield epifluorescence microscope with a Photometrics Prime 95B camera and a Plan Apo 100x, 1.45-numerical-aperture (NA) lens objective. Cells were taken from log. phase culture in 7H9 and immobilized on 1.5% agarose pads made with Hdb media. To detect GFPmut3 and splitGFP signal, a filter cube with a 470/40nm excitation filter, a 525/50nm emission filter and a 495nm dichroic mirror was used. To detect mCherry2B signal, a filter cube with a 560/40nm excitation filter, a 630/70nm emission filter and a 585nm dichroic mirror was used. To detect HADA, a filter cube with a 350/50nm excitation filter, a 460/50nm emission filter, and a 400nm dichroic mirror was used. Image analysis was performed using MicrobeJ (46) to make cell ROIs and extract fluorescence data from them. Fluorescence intensity data from MicrobeJ was further analyzed using bespoke MATLAB code (see supplement).

### SplitGFP probing of Wag31 homo-oligomerization topology

The split superfolder GFP used was described in (33). GFP1-10 (beta-strands 1-10 of superfolderGFP) was fused to both the N and C terminus of wild-type *wag31*_Msmeg_ in L5 integrating vectors under a P750-tet promotor. GFP11(final beta strand) was fused to the C-terminus of *wag31*_Msmeg_ wild-type or mutant alleles or mcherry2B under a P750-tet promoter in a Giles integrating vector. All strains were grown to log. phase (OD600 = 0.15-0.2) and induced with Anhydrotetracycline hydrochloride (ATc) for 15 minutes, then stained with HADA and continued inducing for 15 min more, then washed with PBS tween80, fixed with 1.6% paraformaldehyde for 10 min, then resuspended again in PBS tween 80 before microscopy. HADA staining allows us to distinguish the poles, as it stains the old pole more brightly (37). All microscopy images were analyzed with Fiji and MicrobeJ. Unpaired T-test was used to calculate the P-values using GraphPad Prism.

### Cell staining

Cultures were stained with 1 μg/mL HADA (R&D systems) for 15 minutes with rolling and incubation at 37°C. Stained cultures were then pelleted and washed in 7H9 before imaging. For the pulse-chase-pulse elongation test, cells after 15 minutes of HADA staining were resuspended into 7H9 for 1.5 hours of rolling and incubation at 37°C, then stained with 1 μg/mL NADA (R&D systems) for 2 minutes at room temperature, washed and resuspended in 7H9 before imaging.

## Acknowledgements

This work was supported by NIH grant R01AI148917 to CCB.

