## Supplemental files for "Polar growth protein Wag31 undergoes changes in homo-oligomeric network topology, and has distinct functions at both cell poles and the septum"

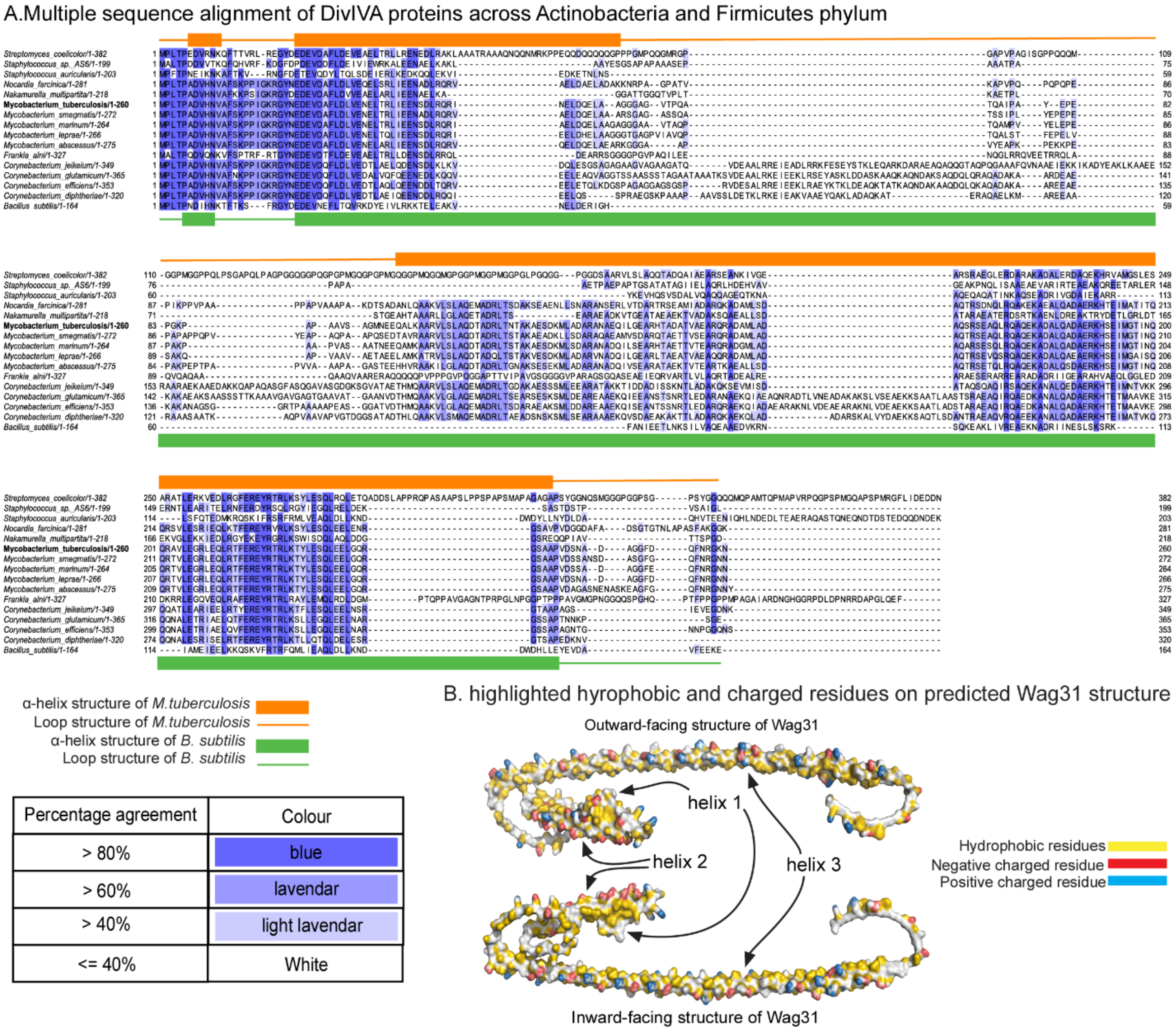

**Supplemental Figure 1. Wag31 vs. DivIVA.** (A) Multiple sequence alignment of DivIVA proteins across Actinobacteria and Firmicutes phyla using Clustal Omega program (1–3) and visualized with Jalview version 2(4, 5). Secondary structure of is shown as boxes (helices) and lines (loops) at the top (*M*) bottom (*B. subtilis*) of the alignments. (B) Predicted Alphafold2 (6) Wag31 structure with hydrophobic and charged residues on highlighted on the surface (7).

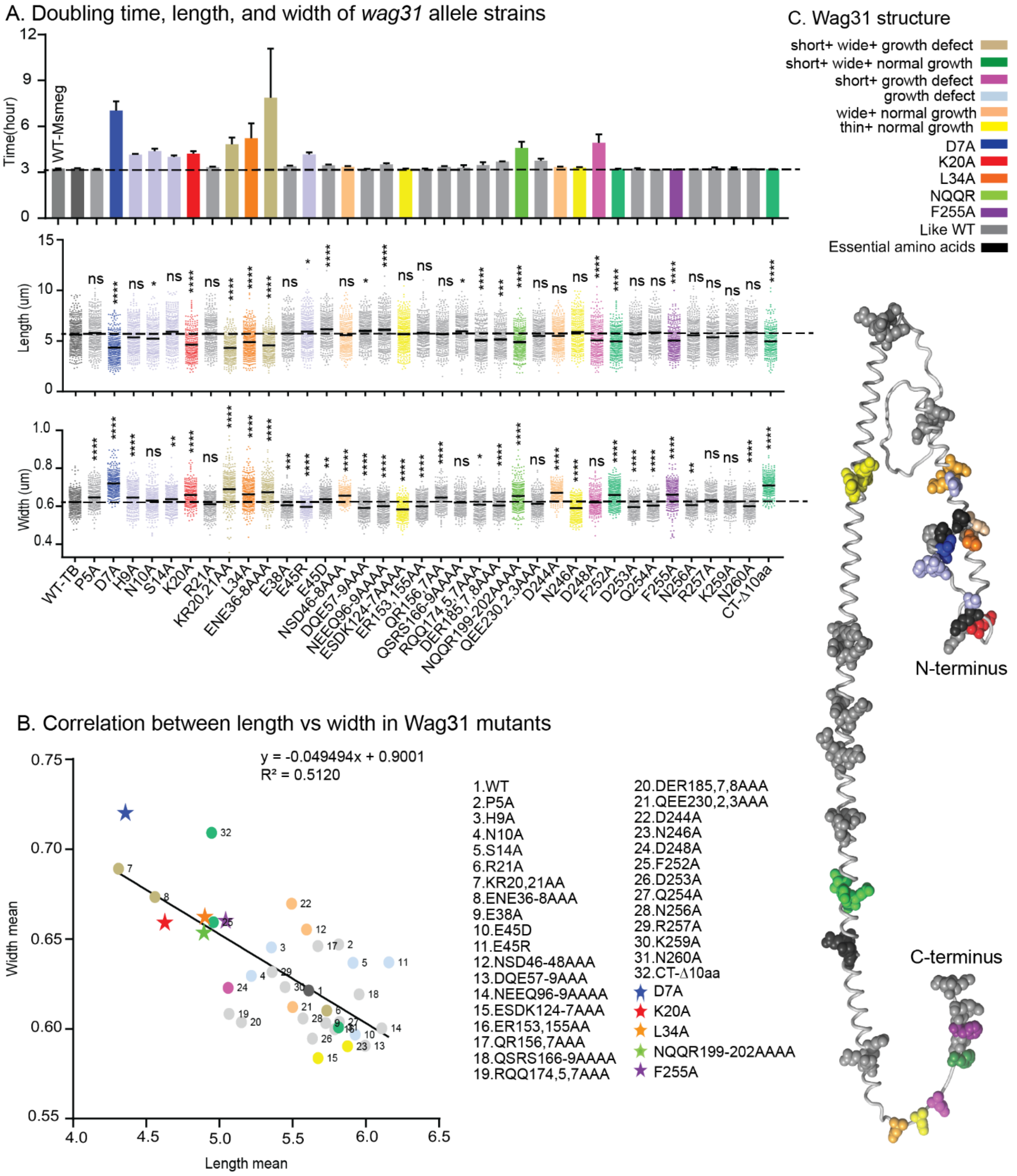

**Supplemental Figure 2. Gross cell phenotypes.** (A) Doubling time (top), cell length (middle) and cell width (bottom) in *wag31* allele strains. Cells with a mean at least 10% less than the WT mean, and with a width at least 5% more than the mean of the WT strain, are considered short and wide, respectively. ns, P >0.05, *, P =< 0.05, **, P=< 0.005, ***, P=< 0.0005, ****, P= <0.0001. *P*-value is calculated by one-way ANOVA, Dunnett's multiple comparisons test. (B) Mean length of *wag31* allele strains plotted as a function of their mean width. Black line is a linear fit. (C) Residues mutated in the alanine scanning mutagenesis are shown as spheres on the Alphafold2 predicted structure of Wag31. Residues are colored according to phenotype categories in (A). Essential residues (black) are the mutants that were unable to replace the *wag31* WT allele.

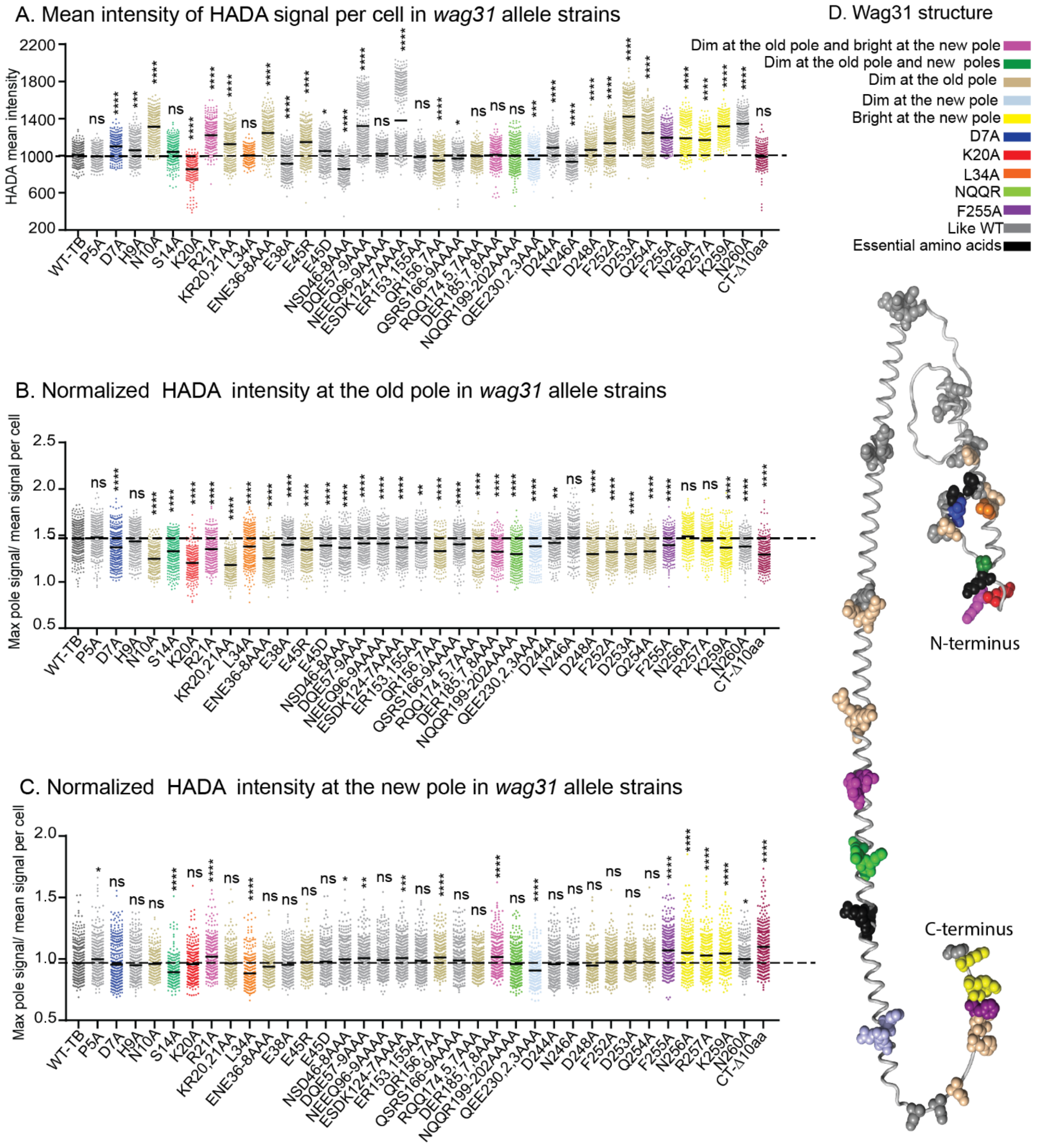

**Supplemental Figure 3: Polar peptidoglycan metabolism.** (A) Mean intensity of HADA signal per cell in *wag31* allele strains. (B), (C) Relative HADA signal at the old and new poles, normalized to the cell mean. Black bars are at the mean. Cells with polar intensity at least 7% less than the mean of the *wag31* WT strain are considered dim at the old pole. Cells with polar intensity at least 5% more than the mean of the *wag31* WT strain are considered bright at the new pole. Cells with polar intensity at least 5% less than the mean of the *wag31* WT strain are considered dim at the new pole. (D) Residues mutated in the alanine scanning mutagenesis are shown as spheres on the Alphafold2 predicted structure of Wag31. Residues are colored according to phenotype categories in A,B,C.  ns, P >0.05, *, P =< 0.05, **, P=< 0.005, ***, P=< 0.0005, ****, P= <0.0001. *P*-value is calculated by one-way ANOVA, Dunnett's multiple comparisons test.

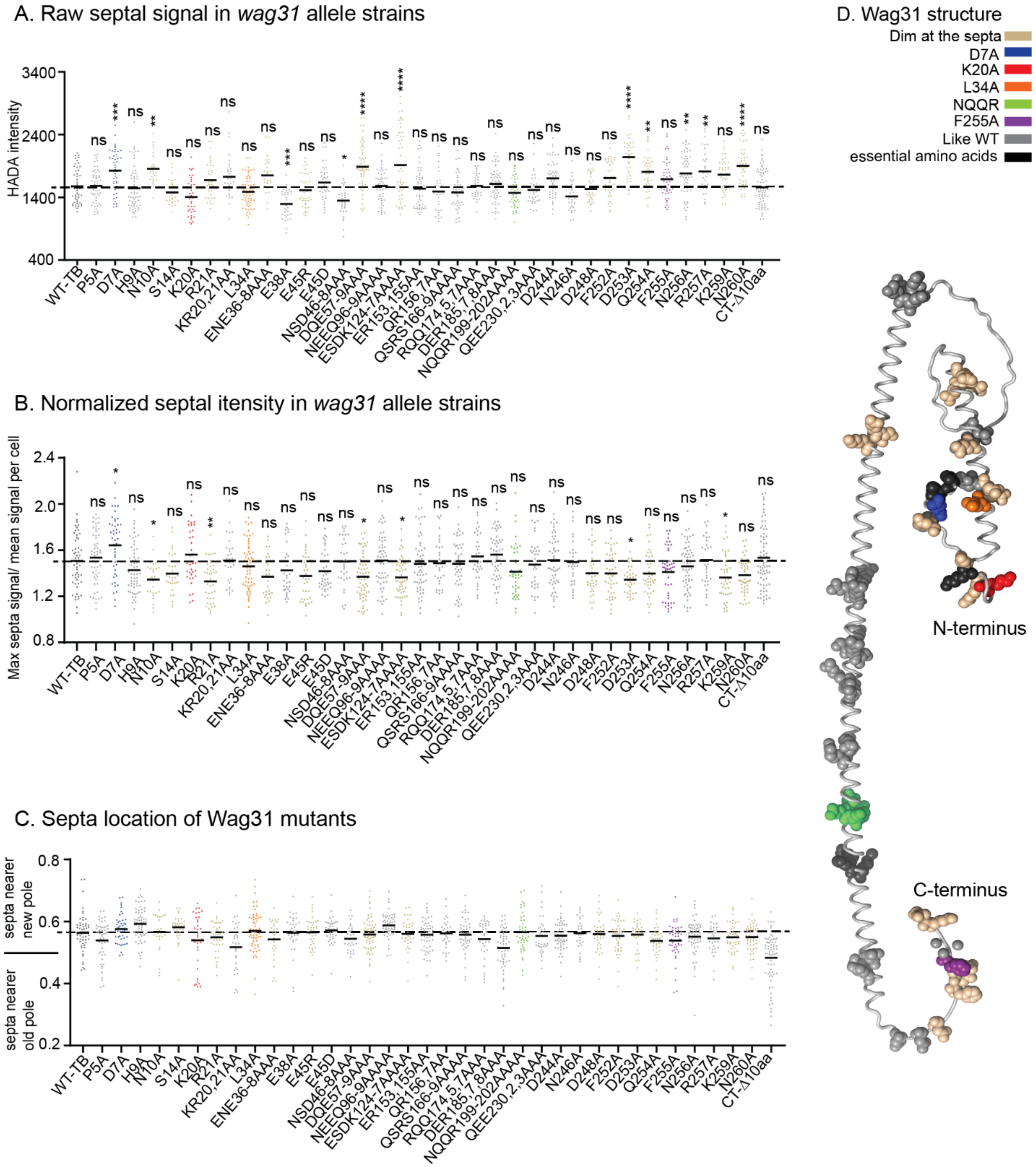

**Supplemental Figure 4: Septal peptidoglycan metabolism.** (A) Raw and, (B) relative septal HADA intensity of *wag31* allele strains. Black bars are at the mean. (C) Septal location in *wag31* allele strains. (D) Residues mutated in the alanine scanning mutagenesis are shown as spheres on the Alphafold2 predicted structure of Wag31.

Residues that have at least 7% less normalized septal intensity than the WT are defined as dim. ns, P >0.05, *, P =< 0.05, **, P=< 0.005, ***, P=< 0.0005, ****, P= <0.0001. *P*-value is calculated by one-way ANOVA, Dunnett's multiple comparisons test.

Demographs of HADA intensity of *wag31* allele strains

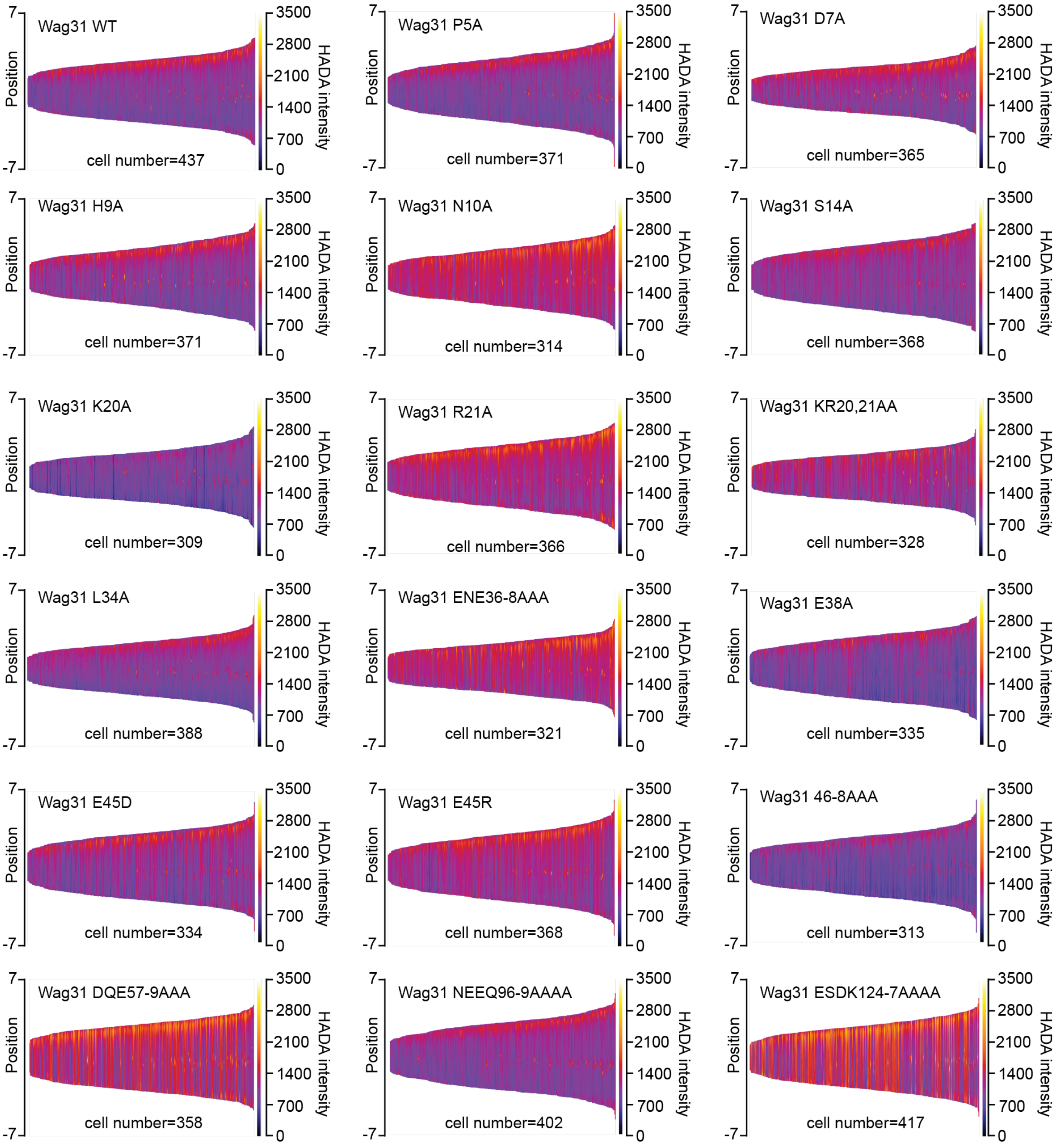

**Supplemental Figure 5:** Demographs of HADA intensity (color scale) across the length of the cell (Y axis) of the *wag31* allele strains. The cells were sorted by length, with shortest cells on the left and longest on the right of each demograph. Cells were also pole-sorted according to HADA intensity, such that the brighter pole (presumed to be the old pole) was oriented to the top along the Y axis. At least 100 cells were analyzed from each of three independent biological replicates of each strain.

Demographs of HADA intensity of *wag31* allele strains

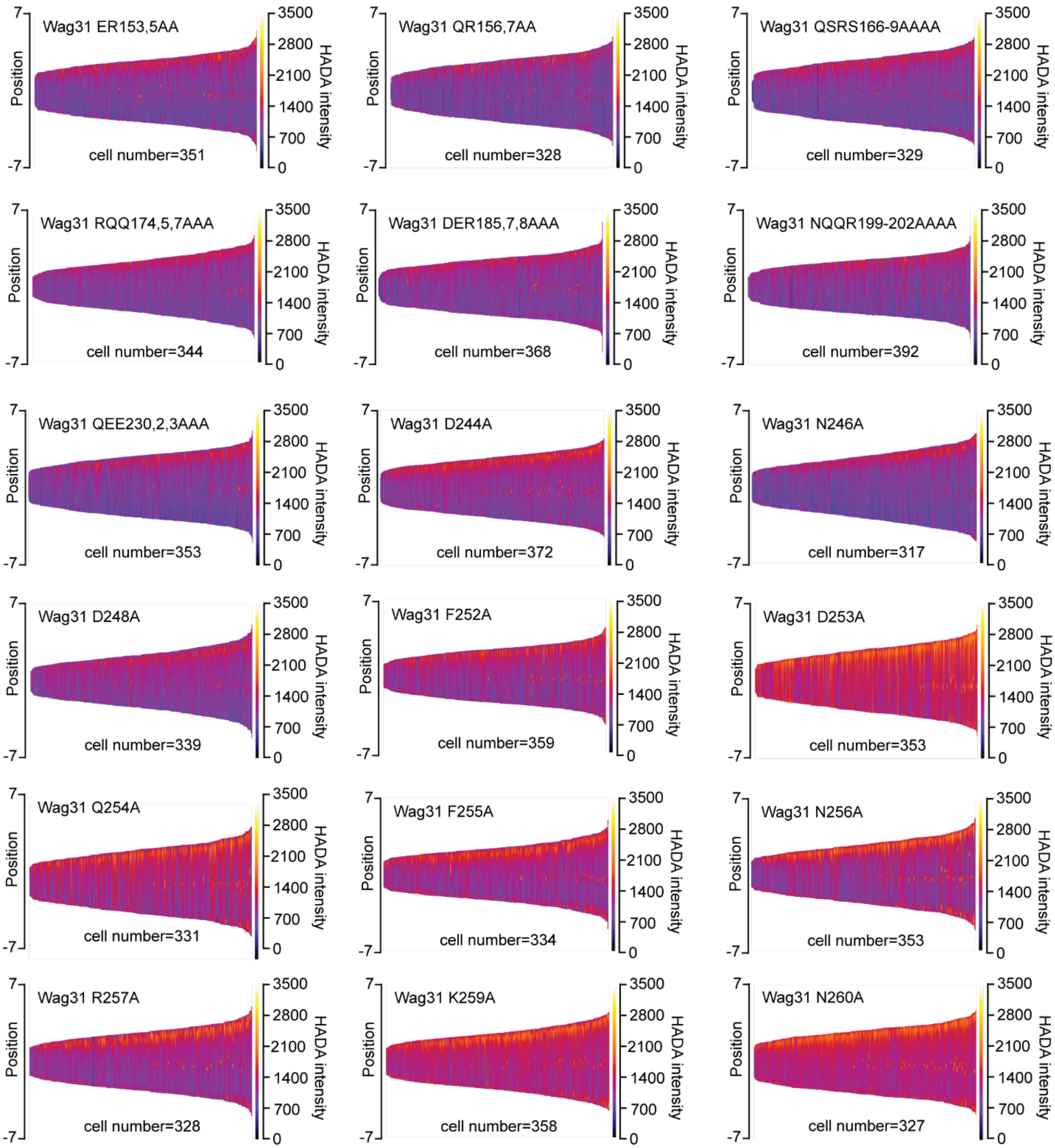

**Supplemental Figure 6:** Demographs of HADA intensity (color scale) across the length of the cell (Y axis) of the *wag31* allele strains. The cells were sorted by length, with shortest cells on the left and longest on the right of each demograph. Cells were also pole-sorted according to HADA intensity, such that the brighter pole (presumed to be the old pole) was oriented to the top along the Y axis. At least 100 cells were analyzed from each of three independent biological replicates of each strain.

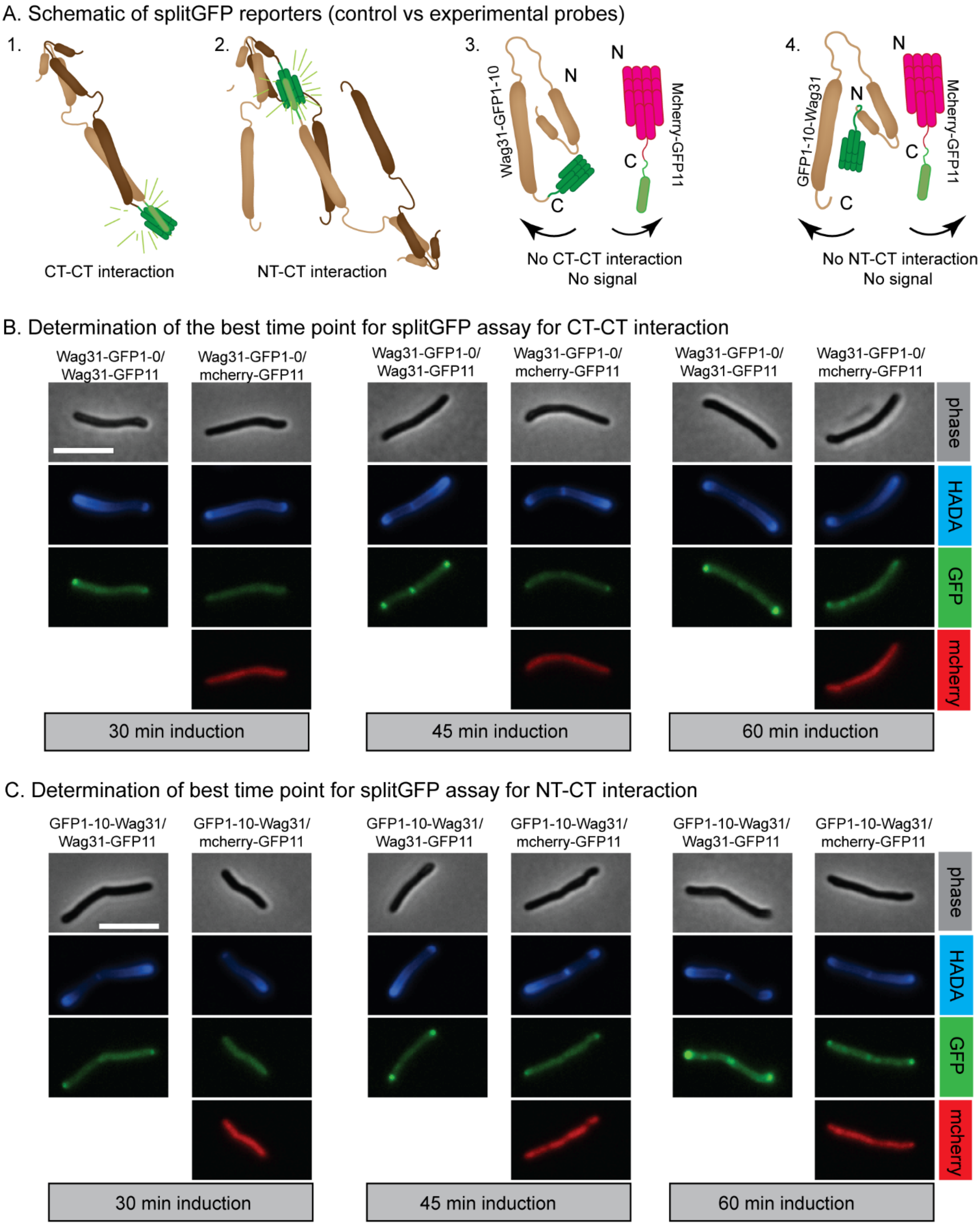
A. Schematic of splitGFP reporters (control vs experimental probes)
1.
2.
3.
4.
CT-CT interaction
NT-CT interaction
Wag31-GFP1-10
N
C
N
C
Mcherry-GFP11
No CT-CT interaction
No signal
GFP1-10-Wag31
N
C
N
C
Mcherry-GFP11
No NT-CT interaction
No signal
B. Determination of the best time point for splitGFP assay for CT-CT interaction
Wag31-GFP1-0/
Wag31-GFP11
Wag31-GFP1-0/
mcherry-GFP11
phase
HADA
GFP
mcherry
30 min induction
45 min induction
60 min induction
C. Determination of best time point for splitGFP assay for NT-CT interaction
GFP1-10-Wag31/
Wag31-GFP11
GFP1-10-Wag31/
mcherry-GFP11
phase
HADA
GFP
mcherry
30 min induction
45 min induction
60 min induction

**Supplemental Figure 7:** (A) Schematic of splitGFP reporter and control strains. A1 and A2 shows experimental reporter strains. A3 and A4 are schematic of control strains. (B,C) Representative images of cells of the Wag31 splitGFP reporters and control strains taken at 30, 45 and 60 minutes after induction of the splitGFP fusion proteins. (B) CT-CT reporter and controls. (C) NT-CT reporter and controls. Polar splitGFP green signal in the control strains is indicative of artefactual signal from self-association of the splitGFP fragments. The scale bar is 5 microns, and applies to all images.

**Table S1A. Results of Wag31 WT and Wag31 mutants interaction using two-hybrid assay**

| Wag31 mutants | Growth on 7H11 plate | Growth on 7H11 +TMP plate |
|---|---|---|
| WT | + | + |
| PLTP2-5AAAA | + | + |

| | | |
|---|---|---|
| DVH7-9AAA | + | + |
| NVF10, 11, 13AAA | + | - |
| SKPP14-17AAAA | + | - |
| IGKR18-21AAAA | + | - |
| N10A | + | - |
| F13A | + | - |
| S14A | + | - |
| K15A | + | - |
| K20A | + | - |
| R21A | + | - |
| KR20, 21-AA | + | - |
| E27A | + | - |
| EDED25-29AAAA | + | - |
| FD31, 33AA | + | - |
| L34A | + | - |
| ENE36-38AAA | + | - |
| E38A | + | - |
| TRE40, 41, 44AAA | + | - |
| R41A | + | - |
| E45R | + | - |
| E45D | + | - |
| ΔNSD46-48 | + | - |
| NSD46-48AAA | + | - |
| DQE57-9AAA | + | - |
| NEEQ96-99AAAA | + | - |
| ESDK124-127AAA | + | - |
| ERH141-143AAA | + | - |
| ER153,155AA | + | - |
| QR156,7AA | + | - |
| QSRS166-9AAAA | + | - |
| RQQ174,5,7AAA | + | - |
| DER-185,7,8AAA | + | - |
| NQQR-199-202AAAA | + | - |
| EQR210,11,13AAA | + | - |
| RRK220,2,4AAA | + | - |
| QEE230,2,3AAA | + | - |
| GGF250-2AAA | + | - |
| RGKN257-260AAAA | + | - |
| DQFN253-6AAAA | + | - |
| D244A | + | - |
| N246A | + | - |
| D248A | + | - |
| F252A | + | - |
| D253A | + | - |
| Q254A | + | - |

| | | |
|---|---|---|
| F255A | + | - |
| N256A | + | - |
| R257A | + | - |
| K259A | + | - |
| N260A | + | - |
| ∆Ct-10aa | + | - |

**Table S2A. Strain list.**

| Strains | Genotype | Figure panel |
|---|---|---|
| CB1612 | mc$^2$155 L5::pUAB400 -hsp60-dhfr3/ pUAB300- hsp60 -dhfr1,2 | 1 |
| CB1613 | mc$^2$155 L5::pUAB200 -hsp60-gcn4-dhfr3/ pUAB100- hsp60-gcn4 -dhfr1,2 | 1 |
| CB1704 | mc$^2$155 L5::pUAB200- wag31(Mtb)-wt-dhfr3/ pUAB100- wag31(Mtb)-wt-dhfr1,2 | 1 |
| CB1757 | mc$^2$155 L5::pUAB200-wag31(Mtb)-∆Ct -10aa-dhfr3/ pUAB100-wag31(Mtb)-wt-dhfr1,2 | 1 |
| CB1769 | mc$^2$155 L5::pUAB200- wag31(Mtb)-PLTP2-5AAAA-dhfr3/ pUAB100-wag31(Mtb)-wt-dhfr1,2 | 1 |
| CB1770 | mc$^2$155 L5::pUAB200- wag31(Mtb)-DVH7-9AAA-dhfr3/ pUAB100-wag31(Mtb)-wt-dhfr1,2 | 1 |
| CB1771 | mc$^2$155 L5::pUAB200- wag31(Mtb)-NVF10, 11, 13AAA-dhfr3/ pUAB100- wag31(Mtb)-wt-dhfr1,2 | 1 |
| CB1772 | mc$^2$155 L5::pUAB200- wag31(Mtb)-SKPP14-17AAAA-dhfr3/ pUAB100- wag31(Mtb)-wt-dhfr1,2 | 1 |
| CB1773 | mc$^2$155 L5::pUAB200- wag31(Mtb)-IGKR18-21AAAA-dhfr3/ pUAB100-wag31(Mtb)-wt-dhfr1,2 | 1 |
| CB1800 | mc$^2$155 L5::pUAB200- wag31(Mtb)- RGKN257-260AAAA-dhfr3/ pUAB100- wag31(Mtb)-wt-dhfr1,2 | 1 |
| CB1801 | mc$^2$155 L5::pUAB200- wag31(Mtb)- DQFN253-6AAAA-dhfr3/ pUAB100- wag31(Mtb)-wt-dhfr1,2 | 1 |
| CB1802 | mc$^2$155 L5::pUAB200- wag31(Mtb)- GGF250-2AAA-dhfr3/ pUAB100-wag31(Mtb)-wt-dhfr1,2 | 1 |
| CB1843 | mc$^2$155 L5::pUAB200- wag31(Mtb)-N10A-dhfr3/ pUAB100-wag31(Mtb)-wt-dhfr1,2 | 1 |
| CB1844 | mc$^2$155 L5::pUAB200- wag31(Mtb)-F13A-dhfr3/ pUAB100-wag31(Mtb)-wt-dhfr1,2 | 1 |
| CB1845 | mc$^2$155 L5::pUAB200- wag31(Mtb)-S14A-dhfr3/ pUAB100-wag31(Mtb)-wt-dhfr1,2 | 1 |
| CB1846 | mc$^2$155 L5::pUAB200- wag31(Mtb)-K15A-dhfr3/ pUAB100-wag31(Mtb)-wt-dhfr1,2 | 1 |
| CB1847 | mc$^2$155 L5::pUAB200- wag31(Mtb)-K20A-dhfr3/ pUAB100-wag31(Mtb)-wt-dhfr1,2 | 1 |
| CB1848 | mc$^2$155 L5::pUAB200- wag31(Mtb)-R21A-dhfr3/ pUAB100-wag31(Mtb)-wt-dhfr1,2 | 1 |

| | | |
|---|---|---|
| CB1849 | $mc^2155$ L5::pUAB200- wag31(Mtb)-KR20, 21AA-dhfr3/ pUAB100- wag31(Mtb)-wt-dhfr1,2 | 1 |
| CB1850 | $mc^2155$ L5::pUAB200- wag31(Mtb)-EDED25-29AAAA-dhfr3/ pUAB100- wag31(Mtb)1-wt-dhfr1,2 | 1 |
| CB1851 | $mc^2155$ L5::pUAB200- wag31(Mtb)-FD31,33AA-dhfr3/ pUAB100- wag31(Mtb)-wt-dhfr1,2 | 1 |
| CB1852 | $mc^2155$ L5::pUAB200- wag31(Mtb)-ENE36-38AAA-dhfr3/ pUAB100- wag31(Mtb)-wt-dhfr1,2 | 1 |
| CB1853 | $mc^2155$ L5::pUAB200- wag31(Mtb)-TRE40-41-44AAA-dhfr3/ pUAB100- wag31(Mtb)-wt-dhfr1,2 | 1 |
| CB1891 | $mc^2155$ L5::pUAB200- wag31(Mtb)-E45R-dhfr3/ pUAB100- wag31(Mtb)-wt-dhfr1,2 | 1 |
| CB1892 | $mc^2155$ L5::pUAB200- wag31(Mtb)-E45D-dhfr3/ pUAB100- wag31(Mtb)-wt-dhfr1,2 | 1 |
| CB1893 | $mc^2155$ L5::pUAB200- wag31(Mtb)-N260A-dhfr3/ pUAB100- wag31(Mtb)-wt-dhfr1,2 | 1 |
| CB1894 | $mc^2155$ L5::pUAB200- wag31(Mtb)-K259A-dhfr3/ pUAB100- wag31(Mtb)-wt-dhfr1,2 | 1 |
| CB1895 | $mc^2155$ L5::pUAB200- wag31(Mtb)-R257A-dhfr3/ pUAB100- wag31(Mtb)-wt-dhfr1,2 | 1 |
| CB1896 | $mc^2155$ L5::pUAB200- wag31(Mtb)-N256A-dhfr3/ pUAB100- wag31(Mtb)-wt-dhfr1,2 | 1 |
| CB1897 | $mc^2155$ L5::pUAB200- wag31(Mtb)-F255A-dhfr3/ pUAB100- wag31(Mtb)-wt-dhfr1,2 | 1 |
| CB1898 | $mc^2155$ L5::pUAB200- wag31(Mtb)-Q254A-dhfr3/ pUAB100- wag31(Mtb)-wt-dhfr1,2 | 1 |
| CB1899 | $mc^2155$ L5::pUAB200- wag31(Mtb)-D253A-dhfr3/ pUAB100- wag31(Mtb) -wt-dhfr1,2 | 1 |
| CB1900 | $mc^2155$ L5::pUAB200- wag31(Mtb)-F252A-dhfr3/ pUAB100- wag31(Mtb)-wt-dhfr1,2 | 1 |
| CB1901 | $mc^2155$ L5::pUAB200- wag31(Mtb)-D248A-dhfr3/ pUAB100- wag31(Mtb)-wt-dhfr1,2 | 1 |
| CB1902 | $mc^2155$ L5::pUAB200- wag31(Mtb)-DQE57-9AAA-dhfr3/ pUAB100- wag31(Mtb)-wt-dhfr1,2 | 1 |
| CB1903 | $mc^2155$ L5::pUAB200- wag31(Mtb)-NEEQ96-9AAAA-dhfr3/ pUAB100- wag31(Mtb)-wt-dhfr1,2 | 1 |
| CB1904 | $mc^2155$ L5::pUAB200- wag31(Mtb)-ESDK124-127AAAA-dhfr3/ pUAB100- wag31(Mtb)-wt-dhfr1,2 | 1 |
| CB1905 | $mc^2155$ L5::pUAB200- wag31(Mtb)-ERH141,143,144AAA-dhfr3/ pUAB100- wag31(Mtb)-wt-dhfr1,2 | 1 |
| CB2101 | $mc^2155$ L5::pUAB200- wag31(Mtb)-E27A-dhfr3/ pUAB100- wag31(Mtb)-wt-dhfr1,2 | 1 |
| CB2102 | $mc^2155$ L5::pUAB200- wag31(Mtb)-E38A-dhfr3/ pUAB100- wag31(Mtb)-wt-dhfr1,2 | 1 |

| | | |
|---|---|---|
| CB2103 | mc2155 L5::pUAB200- wag31(Mtb)-L34A-dhfr3/ pUAB100-wag31(Mtb)-wt-dhfr1,2 | 1 |
| CB2104 | mc2155 L5::pUAB200- wag31(Mtb)-R41A-dhfr3/ pUAB100-wag31(Mtb)-wt-dhfr1,2 | 1 |
| CB2105 | mc2155 L5::pUAB200- wag31(Mtb)-NSD46-8AAA-dhfr3/ pUAB100-wag31(Mtb)-wt-dhfr1,2 | 1 |
| CB2106 | mc2155 L5::pUAB200- wag31(Mtb)-ΔNSD46-48-dhfr3/ pUAB100-wag31(Mtb)-wt-dhfr1,2 | 1 |
| CB2107 | mc2155 L5::pUAB200- wag31(Mtb)-ER153,155AA-dhfr3/ pUAB100-wag31(Mtb)-wt-dhfr1,2 | 1 |
| CB2108 | mc2155 L5::pUAB200- wag31(Mtb)-QR156,157AA-dhfr3/ pUAB100-wag31(Mtb)-wt-dhfr1,2 | 1 |
| CB2109 | mc2155 L5::pUAB200- wag31(Mtb)-QSRS166-9AAAA-dhfr3/ pUAB100-wag31(Mtb)-wt-dhfr1,2 | 1 |
| CB2110 | mc2155 L5::pUAB200- wag31(Mtb)-RQQ174,5,7AAA-dhfr3/ pUAB100-wag31(Mtb)-wt-dhfr1,2 | 1 |
| CB2111 | mc2155 L5::pUAB200- wag31(Mtb)-DER185,7,8AAA-dhfr3/ pUAB100-wag31(Mtb)-wt-dhfr1,2 | 1 |
| CB2112 | mc2155 L5::pUAB200- wag31(Mtb)-NQQR199-202AAAA-dhfr3/ pUAB100- wag31(Mtb)-wt-dhfr1,2 | 1 |
| CB2113 | mc2155 L5::pUAB200- wag31(Mtb)-EQR210,11,13AAA-dhfr3/ pUAB100- wag31(Mtb)-wt-dhfr1,2 | 1 |
| CB2114 | mc2155 L5::pUAB200- wag31(Mtb)-RRK220,2,4AAA-dhfr3/ pUAB100-wag31(Mtb)-wt-dhfr1,2 | 1 |
| CB2115 | mc2155 L5::pUAB200- wag31(Mtb)-QEE230,2,3AAA-dhfr3/ pUAB100-wag31(Mtb)-wt-dhfr1,2 | 1 |
| CB2116 | mc2155 L5::pUAB200- wag31(Mtb)-D244A-dhfr3/ pUAB100-wag31(Mtb)-wt-dhfr1,2 | 1 |
| CB2117 | mc2155 L5::pUAB200- wag31(Mtb)-N246A-dhfr3/ pUAB100-wag31(Mtb)-wt-dhfr1,2 | 1 |
| CB1821 | mc2155 Δwag31::hygR L5::pCT94-Pwag31-wag31(Msmeg)/ pNitRecET | |
| CB1927 | mc2155 Δwag31::hygR L5::pKK158-Pwag31-wag31(Mtb)-wt/ pNitRecET- clone1 | 3 |
| CB1928 | mc2155 Δwag31::hygR L5::pKK158- Pwag31-wag31(Mtb)-wt/ pNitRecET- clone2 | 3 |
| CB1929 | mc2155 Δwag31::hygR L5::pKK158- Pwag31-wag31(Mtb)-wt/ pNitRecET- clone3 | 3 |
| CB1966 | mc2155 Δwag31::hygR L5::pKK158 Pwag31-wag31(Mtb)-N10A/ pNitRecET- clone1 | S2, S3, S4, S5 |
| CB1967 | mc2155 Δwag31::hygR L5:: pKK158 Pwag31-wag31(Mtb)-N10A/ pNitRecET- clone2 | S2, S3, S4, S5 |
| CB1968 | mc2155 Δwag31::hygR L5:: pKK158 Pwag31-wag31(Mtb)-N10A/ pNitRecET- clone3 | S2, S3, S4, S5 |

| | | |
|---|---|---|
| CB1969 | mc$^2$155 Δwag31::hygR L5:: pKK158 Pwag31-wag31(Mtb)-KR20,21AA/ pNitRecET- clone1 | S2, S3, S4, S5 |
| CB1970 | mc$^2$155 Δwag31::hygR L5:: pKK158 Pwag31-wag31(Mtb)-KR20,21AA/ pNitRecET- clone2 | S2, S3, S4, S5 |
| CB1971 | mc$^2$155 Δwag31::hygR L5:: pKK158 Pwag31-wag31(Mtb)-KR20,21AA/ pNitRecET- clone3 | S2, S3, S4, S5 |
| CB1985 | mc$^2$155 Δwag31::hygR L5:: pKK158 Pwag31-wag31(Mtb)-ENE36-8AAA/ pNitRecET- clone1 | S2, S3, S4 ,S5 |
| CB1986 | mc$^2$155 Δwag31::hygR L5:: pKK158 Pwag31-wag31(Mtb)-ENE36-8AAA/ pNitRecET- clone2 | S2, S3, S4, S5 |
| CB1987 | mc$^2$155 Δwag31::hygR L5:: pKK158 Pwag31-wag31(Mtb)-ENE36-8AAA/ pNitRecET- clone3 | S2, S3, S4 ,S5 |
| CB1988 | mc$^2$155 Δwag31::hygR L5:: pKK158 Pwag31-wag31(Mtb)-K257A/ pNitRecET- clone1 | S2, S3, S4, S6 |
| CB1989 | mc$^2$155 Δwag31::hygR L5:: pKK158 Pwag31-wag31(Mtb)-K257A/ pNitRecET- clone2 | S2, S3, S4, S6 |
| CB1990 | mc$^2$155 Δwag31::hygR L5:: pKK158 Pwag31-wag31(Mtb)-K257A/ pNitRecET- clone3 | S2, S3, S4, S6 |
| CB2025 | mc$^2$155 Δwag31::hygR L5:: pKK158 Pwag31-wag31(Mtb)-S14A/ pNitRecET- clone1 | S2, S3, S4, S5 |
| CB2026 | mc$^2$155 Δwag31::hygR L5:: pKK158 Pwag31-wag31(Mtb)-S14A/ pNitRecET- clone2 | S2, S3, S4, S5 |
| CB2027 | mc$^2$155 Δwag31::hygR L5:: pKK158 Pwag31-wag31(Mtb)-S14A/ pNitRecET- clone3 | S2, S3, S4, S5 |
| CB2028 | mc$^2$155 Δwag31::hygR L5:: pKK158 Pwag31-wag31(Mtb)-K20A/ pNitRecET- clone1 | 3, S2, S3, S4, S5 |
| CB2029 | mc$^2$155 Δwag31::hygR L5:: pKK158 Pwag31-wag31(Mtb)-K20A/ pNitRecET- clone2 | 3, S2, S3, S4, S5 |
| CB2030 | mc$^2$155 Δwag31::hygR L5:: pKK158 Pwag31-wag31(Mtb)-K20A/ pNitRecET- clone3 | 3, S2, S3, S4, S5 |
| CB2031 | mc$^2$155 Δwag31::hygR L5:: pKK158 Pwag31-wag31(Mtb)-R21A/ pNitRecET- clone1 | S2, S3, S4, S5 |
| CB2032 | mc$^2$155 Δwag31::hygR L5:: pKK158 Pwag31-wag31(Mtb)-R21A/ pNitRecET- clone2 | S2, S3, S4, S5 |
| CB2033 | mc$^2$155 Δwag31::hygR L5:: pKK158 Pwag31-wag31(Mtb)-F255A/ pNitRecET- clone1 | 3, S2, S3, S4, S6 |
| CB2034 | mc$^2$155 Δwag31::hygR L5:: pKK158 Pwag31-wag31(Mtb)-F255A/ pNitRecET- clone2 | 3, S2, S3, S4, S6 |
| CB2035 | mc$^2$155 Δwag31::hygR L5:: pKK158 Pwag31-wag31(Mtb)-F255A/ pNitRecET- clone3 | 3, S2, S3, S4, S6 |
| CB2038 | mc$^2$155 Δwag31::hygR L5:: pKK158 Pwag31-wag31(Mtb)-D253A/ pNitRecET- clone1 | S2, S3, S4, S6 |
| CB2039 | mc$^2$155 Δwag31::hygR L5:: pKK158 Pwag31-wag31(Mtb)-D253A/ pNitRecET- clone2 | S2, S3, S4, S6 |

| | | |
|---|---|---|
| CB2040 | mc$^2$155 Δwag31::hygR L5:: pKK158 Pwag31-wag31(Mtb)-D253A/ pNitRecET- clone3 | S2, S3, S4, S6 |
| CB2041 | mc$^2$155 Δwag31::hygR L5:: pKK158 Pwag31-wag31(Mtb)-Q254A/ pNitRecET- clone1 | S2, S3, S4, S6 |
| CB2042 | mc$^2$155 Δwag31::hygR L5:: pKK158 Pwag31-wag31(Mtb)-Q254A/ pNitRecET- clone2 | S2, S3, S4, S6 |
| CB2043 | mc$^2$155 Δwag31::hygR L5:: pKK158 Pwag31-wag31(Mtb)-Q254A/ pNitRecET- clone3 | S2, S3, S4, S6 |
| CB2057 | mc$^2$155 Δwag31::hygR L5:: pKK158 Pwag31-wag31(Mtb)-E45R/ pNitRecET- clone1 | S2, S3, S4, S5 |
| CB2058 | mc$^2$155 Δwag31::hygR L5:: pKK158 Pwag31-wag31(Mtb)-E45R/ pNitRecET- clone2 | S2, S3, S4, S5 |
| CB2059 | mc$^2$155 Δwag31::hygR L5:: pKK158 Pwag31-wag31(Mtb)-E45R/ pNitRecET- clone3 | S2, S3, S4, S5 |
| CB2060 | mc$^2$155 Δwag31::hygR L5:: pKK158 Pwag31-wag31(Mtb)-E45D/ pNitRecET- clone1 | S2, S3, S4, S5 |
| CB2061 | mc$^2$155 Δwag31::hygR L5:: pKK158 Pwag31-wag31(Mtb)-E45D/ pNitRecET- clone2 | S2, S3, S4, S5 |
| CB2062 | mc$^2$155 Δwag31::hygR L5:: pKK158 Pwag31-wag31(Mtb)-E45D/ pNitRecET- clone3 | S2, S3, S4, S5 |
| CB2063 | mc$^2$155 Δwag31::hygR L5:: pKK158 Pwag31-wag31(Mtb)-N260A/ pNitRecET- clone1 | S2, S3, S4, S6 |
| CB2064 | mc$^2$155 Δwag31::hygR L5:: pKK158 Pwag31-wag31(Mtb)-N260A/ pNitRecET- clone2 | S2, S3, S4, S6 |
| CB2065 | mc$^2$155 Δwag31::hygR L5:: pKK158 Pwag31-wag31(Mtb)-N260A/ pNitRecET- clone3 | S2, S3, S4, S6 |
| CB2066 | mc$^2$155 Δwag31::hygR L5:: pKK158 Pwag31-wag31(Mtb)-K259A/ pNitRecET- clone1 | S2, S3, S4, S6 |
| CB2067 | mc$^2$155 Δwag31::hygR L5:: pKK158 Pwag31-wag31(Mtb)-K259A/ pNitRecET- clone2 | S2, S3, S4, S6 |
| CB2068 | mc$^2$155 Δwag31::hygR L5:: pKK158 Pwag31-wag31(Mtb)-K259A/ pNitRecET- clone3 | S2, S3, S4, S6 |
| CB2069 | mc$^2$155 Δwag31::hygR L5:: pKK158 Pwag31-wag31(Mtb)-N256A/ pNitRecET- clone1 | S2, S3, S4, S6 |
| CB2070 | mc$^2$155 Δwag31::hygR L5:: pKK158 Pwag31-wag31(Mtb)-N256A/ pNitRecET- clone2 | S2, S3, S4, S6 |
| CB2071 | mc$^2$155 Δwag31::hygR L5:: pKK158 Pwag31-wag31(Mtb)-N256A/ pNitRecET- clone3 | S2, S3, S4, S6 |
| CB2072 | mc$^2$155 Δwag31::hygR L5:: pKK158 Pwag31-wag31(Mtb)-F252A/ pNitRecET- clone1 | S2, S3, S4, S6 |
| CB2073 | mc$^2$155 Δwag31::hygR L5:: pKK158 Pwag31-wag31(Mtb)-F252A/ pNitRecET- clone2 | S2, S3, S4, S6 |
| CB2074 | mc$^2$155 Δwag31::hygR L5:: pKK158 Pwag31-wag31(Mtb)-F252A/ pNitRecET- clone3 | S2, S3, S4, S6 |

| | | |
|---|---|---|
| CB2075 | mc[2]155 Δwag31::hygR L5:: pKK158 Pwag31-wag31(Mtb)-ΔCt 10 aa/ pNitRecET- clone1 | S2, S3, S4 |
| CB2076 | mc[2]155 Δwag31::hygR L5:: pKK158 Pwag31-wag31(Mtb)-ΔCt 10 aa/ pNitRecET- clone2 | S2, S3, S4 |
| CB2077 | mc[2]155 Δwag31::hygR L5:: pKK158 Pwag31-wag31(Mtb)-ΔCt 10 aa/ pNitRecET- clone3 | S2, S3, S4 |
| CB2087 | mc[2]155 Δwag31::hygR L5:: pKK158 Pwag31-wag31(Mtb)-D248A/ pNitRecET- clone1 | S2, S3, S4, S6 |
| CB2088 | mc[2]155 Δwag31::hygR L5:: pKK158 Pwag31-wag31(Mtb)-D248A/ pNitRecET- clone2 | S2, S3, S4, S6 |
| CB2089 | mc[2]155 Δwag31::hygR L5:: pKK158 Pwag31-wag31(Mtb)-D248A/ pNitRecET- clone3 | S2, S3, S4, S6 |
| CB2090 | mc[2]155 Δwag31::hygR L5:: pKK158 Pwag31-wag31(Mtb)-DQE 57-9AAA/ pNitRecET- clone1 | S2, S3, S4, S5 |
| CB2091 | mc[2]155 Δwag31::hygR L5:: pKK158 Pwag31-wag31(Mtb)-DQE 57-9AAA/ pNitRecET- clone2 | S2, S3, S4, S5 |
| CB2092 | mc[2]155 Δwag31::hygR L5:: pKK158 Pwag31-wag31(Mtb)-DQE 57-9AAA/ pNitRecET- clone3 | S2, S3, S4, S5 |
| CB2093 | mc[2]155 Δwag31::hygR L5:: pKK158 Pwag31-wag31(Mtb)-NEEQ 96-9AAAA/ pNitRecET- clone1 | S2, S3, S4, S5 |
| CB2094 | mc[2]155 Δwag31::hygR L5:: pKK158 Pwag31-wag31(Mtb)-NEEQ 96-9AAAA/ pNitRecET- clone2 | S2, S3, S4, S5 |
| CB2095 | mc[2]155 Δwag31::hygR L5:: pKK158 Pwag31-wag31(Mtb)-NEEQ 96-9AAAA/ pNitRecET- clone3 | S2, S3, S4, S5 |
| CB2096 | mc[2]155 Δwag31::hygR L5:: pKK158 Pwag31-wag31(Mtb)-ESDK 124-127AAAA/ pNitRecET- clone1 | S2, S3, S4, S5 |
| CB2097 | mc[2]155 Δwag31::hygR L5:: pKK158 Pwag31-wag31(Mtb)-ESDK 124-127AAAA/ pNitRecET- clone2 | S2, S3, S4, S5 |
| CB2098 | mc[2]155 Δwag31::hygR L5:: pKK158 Pwag31-wag31(Mtb)-ESDK 124-127AAAA/ pNitRecET- clone3 | S2, S3, S4, S5 |
| CB2130 | mc[2]155 Δwag31::hygR L5:: pKK158 Pwag31-wag31(Mtb)- DQFN253-6AAAA/ pNitRecET- clone1 | S2, S3, S4, S6 |
| CB2131 | mc[2]155 Δwag31::hygR L5:: pKK158 Pwag31-wag31(Mtb)- DQFN253-6AAAA/ pNitRecET- clone2 | S2, S3, S4, S6 |
| CB2132 | mc[2]155 Δwag31::hygR L5:: pKK158 Pwag31-wag31(Mtb)- DQFN253-6AAAA/ pNitRecET- clone3 | S2, S3, S4, S6 |
| CB2133 | mc[2]155 Δwag31::hygR L5:: pKK158 Pwag31-wag31(Mtb)-GGF250-2AAA/ pNitRecET- clone1 | S2, S3, S4, S6 |
| CB2134 | mc[2]155 Δwag31::hygR L5:: pKK158 Pwag31-wag31(Mtb)-GGF250-2AAA/ pNitRecET- clone2 | S2, S3, S4, S6 |
| CB2135 | mc[2]155 Δwag31::hygR L5:: pKK158 Pwag31-wag31(Mtb)-GGF250-2AAA/ pNitRecET- clone3 | S2, S3, S4, S6 |
| CB2177 | mc[2]155 Δwag31::hygR L5:: pKK158 Pwag31-wag31(Mtb)-ER153,155AA/ pNitRecET- clone1 | S2, S3, S4, S6 |

| | | |
|---|---|---|
| CB2178 | mc$^2$155 Δwag31::hygR L5:: pKK158 Pwag31-wag31(Mtb)-ER153,155AA/ pNitRecET- clone2 | S2, S3, S4, S6 |
| CB2179 | mc$^2$155 Δwag31::hygR L5:: pKK158 Pwag31-wag31(Mtb)-ER153,155AA/ pNitRecET- clone3 | S2, S3, S4, S6 |
| CB2180 | mc$^2$155 Δwag31::hygR L5:: pKK158 Pwag31-wag31(Mtb)- QSRS166-9AAAA/ pNitRecET- clone1 | S2, S3, S4, S6 |
| CB2181 | mc$^2$155 Δwag31::hygR L5:: pKK158 Pwag31-wag31(Mtb)-QSRS166-9AAAA/ pNitRecET- clone2 | S2, S3, S4, S6 |
| CB2182 | mc$^2$155 Δwag31::hygR L5:: pKK158 Pwag31-wag31(Mtb)-QSRS166-9AAAA)/ pNitRecET- clone3 | S2, S3, S4, S6 |
| CB2183 | mc$^2$155 Δwag31::hygR L5:: pKK158 Pwag31-wag31(Mtb)-NQQR199-202AAAA/ pNitRecET- clone1 | 3, S2, S3, S4, S6 |
| CB2184 | mc$^2$155 Δwag31::hygR L5:: pKK158 Pwag31-wag31(Mtb)-NQQR199-202AAAA/ pNitRecET- clone2 | 3, S2, S3, S4, S6 |
| CB2185 | mc$^2$155 Δwag31::hygR L5:: pKK158 Pwag31-wag31(Mtb)-NQQR199-202AAAA/ pNitRecET- clone3 | 3, S2, S3, S4, S6 |
| CB2186 | mc$^2$155 Δwag31::hygR L5:: pKK158 Pwag31-wag31(Mtb)-QEE230,2,3AAA/ pNitRecET- clone1 | S2, S3, S4, S6 |
| CB2187 | mc$^2$155 Δwag31::hygR L5:: pKK158 Pwag31-wag31(Mtb)-QEE230,2,3AAA/ pNitRecET- clone2 | S2, S3, S4, S6 |
| CB2188 | mc$^2$155 Δwag31::hygR L5:: pKK158 Pwag31-wag31(Mtb)-QEE230,2,3AAA/ pNitRecET- clone3 | S2, S3, S4, S6 |
| CB2195 | mc$^2$155 Δwag31::hygR L5::pKK158-Pwag31- wag31(Mtb)-L34A/ pNitRecET- clone1 | 3, S2, S3, S4, S5 |
| CB2196 | mc$^2$155 Δwag31::hygR L5::pKK158-Pwag31- wag31(Mtb)-L34A/ pNitRecET- clone 2 | 3, S2, S3, S4, S5 |
| CB2197 | mc$^2$155 Δwag31::hygR L5::pKK158-Pwag31- wag31(Mtb)-L34A/ pNitRecET- clone3 | 3, S2, S3, S4, S5 |
| CB2198 | mc$^2$155 Δwag31::hygR L5::pKK158-Pwag31- wag31(Mtb)-E38A/ pNitRecET- clone1 | S2, S3, S4, S5 |
| CB2199 | mc$^2$155 Δwag31::hygR L5::pKK158-Pwag31- wag31(Mtb)-E38A/ pNitRecET- clone2 | S2, S3, S4, S5 |
| CB2200 | mc$^2$155 Δwag31::hygR L5::pKK158-Pwag31- wag31(Mtb)-E38A/ pNitRecET- clone3 | S2, S3, S4, S5 |
| CB2201 | mc2155 Δwag31::hygR L5::pKK158-Pwag31- wag31(Mtb)-NSD46-8AAA/ pNitRecET- clone1 | S2, S3, S4, S5 |
| CB2202 | mc$^2$155 Δwag31::hygR L5::pKK158-Pwag31- wag31(Mtb)-NSD46-8AAA/ pNitRecET- clone2 | S2, S3, S4, S5 |
| CB2203 | mc$^2$155 Δwag31::hygR L5::pKK158-Pwag31- wag31(Mtb)-NSD46-8AAA/ pNitRecET- clone3 | S2, S3, S4, S5 |
| CB2204 | mc$^2$155 Δwag31::hygR L5::pKK158-Pwag31- wag31(Mtb)-QR156,157AA/ pNitRecET- clone1 | S2, S3, S4, S6 |
| CB2205 | mc$^2$155 Δwag31::hygR L5::pKK158-Pwag31- wag31(Mtb)-QR156,157AA/ pNitRecET- clone2 | S2, S3, S4, S6 |

| | | |
|---|---|---|
| CB2206 | mc$^2$155 Δwag31::hygR L5::pKK158-Pwag31- wag31(Mtb)-QR156,157AA/ pNitRecET- clone3 | S2, S3, S4, S6 |
| CB2207 | mc$^2$155 Δwag31::hygR L5::pKK158-Pwag31- wag31(Mtb)-DER185,7,8AAA/ pNitRecET- clone1 | S2, S3, S4, S6 |
| CB2208 | mc$^2$155 Δwag31::hygR L5::pKK158-Pwag31- wag31(Mtb)-DER185,7,8AAA/ pNitRecET- clone2 | S2, S3, S4, S6 |
| CB2209 | mc$^2$155 Δwag31::hygR L5::pKK158-Pwag31- wag31(Mtb)-DER185,7,8AAA/ pNitRecET- clone3 | S2, S3, S4, S6 |
| CB2210 | mc$^2$155 Δwag31::hygR L5::pKK158-Pwag31- wag31(Mtb)-D244A/ pNitRecET- clone1 | S2, S3, S4, S6 |
| CB2211 | mc$^2$155 Δwag31::hygR L5::pKK158-Pwag31- wag31(Mtb)-D244A/ pNitRecET- clone2 | S2, S3, S4, S6 |
| CB2212 | mc$^2$155 Δwag31::hygR L5::pKK158-Pwag31- wag31(Mtb)-D244A/ pNitRecET- clone3 | S2, S3, S4, S6 |
| CB2213 | mc$^2$155 Δwag31::hygR L5::pKK158-Pwag31- wag31(Mtb)-N246A/ pNitRecET- clone1 | S2, S3, S4, S6 |
| CB2214 | mc$^2$155 Δwag31::hygR L5::pKK158-Pwag31- wag31(Mtb)-N246A/ pNitRecET- clone2 | S2, S3, S4, S6 |
| CB2215 | mc$^2$155 Δwag31::hygR L5::pKK158-Pwag31- wag31(Mtb)-N246A/ pNitRecET- clone3 | S2, S3, S4, S6 |
| CB228 | mc$^2$155 L5:: pkk158-Psmeg- wag31(Mtb)-wt-GFP-clone1 | 4 |
| CB2229 | mc$^2$155 L5:: pkk158-Psmeg- wag31(Mtb)-wt-GFP-clone2 | 4 |
| CB2230 | mc$^2$155 L5:: pkk158-Psmeg- wag31(Mtb)-wt-GFP-clone3 | 4 |
| CB2231 | mc$^2$155 Δwag31::hygR L5::pKK158-Pwag31- wag31(Mtb)-H9A/ pNitRecET- clone1 | S2, S3, S4, S5 |
| CB2232 | mc$^2$155 Δwag31::hygR L5::pKK158-Pwag31- wag31(Mtb)-H9A/ pNitRecET- clone2 | S2, S3, S4, S5 |
| CB2233 | mc$^2$155 Δwag31::hygR L5::pKK158-Pwag31- wag31(Mtb)-H9A/ pNitRecET- clone3 | S2, S3, S4, S5 |
| CB2234 | mc$^2$155 Δwag31::hygR L5::pKK158-Pwag31- wag31(Mtb)-RQQ174,5,7AAA/ pNitRecET- clone1 | S2, S3, S4, S6 |
| CB2235 | mc$^2$155 Δwag31::hygR L5::pKK158-Pwag31- wag31(Mtb)-RQQ174,5,7AAA/ pNitRecET- clone2 | S2, S3, S4, S6 |
| CB2236 | mc$^2$155 Δwag31::hygR L5::pKK158-Pwag31- wag31(Mtb)-RQQ174,5,7AAA/ pNitRecET- clone3 | S2, S3, S4, S6 |
| CB2238 | mc$^2$155 Δwag31::hygR L5::pCT197-Pwag31- wag31(Mtb)-T4A / pNitRecET-clone1 | S2, S3, S4, S5 |
| CB2269 | mc$^2$155 Δwag31::hygR L5::pKK158-Pwag31- wag31(Mtb)-P5A/ pNitRecET- clone1 | S2, S3, S4, S5 |
| CB2270 | mc$^2$155 Δwag31::hygR L5::pKK158-Pwag31- wag31(Mtb)-P5A/ pNitRecET- clone2 | S2, S3, S4, S5 |
| CB2271 | mc$^2$155 Δwag31::hygR L5::pKK158-Pwag31- wag31(Mtb)-D7A/ pNitRecET- clone1 | 3, S2, S3, S4, S5 |
| CB2272 | mc$^2$155 Δwag31::hygR L5::pKK158-Pwag31- wag31(Mtb)-D7A/ pNitRecET- clone2 | 3, S2, S3, S4, S5 |

| | | |
|---|---|---|
| CB2273 | mc$^{2}$155 Δwag31::hygR L5::pKK158-Pwag31- wag31(Mtb)-D7A/ pNitRecET- clone3 | 3, S2, S3, S4, S5 |
| CB2296 | mc$^{2}$155 L5:: pkk158-Psmeg- wag31(Mtb)-NQQR199-202AAAA-GFP-clone1 | 4 |
| CB2297 | mc$^{2}$155 L5:: pkk158-Psmeg- wag31(Mtb)-NQQR199-202AAAA-GFP-clone2 | 4 |
| CB2298 | mc$^{2}$155 L5:: pkk158-Psmeg- wag31(Mtb)-NQQR199-202AAAA-GFP-clone3 | 4 |
| CB2305 | mc$^{2}$155 L5:: pkk158-Psmeg- wag31(Mtb)-L34A-GFP-clone1 | 4 |
| CB2306 | mc$^{2}$155 L5:: pkk158-Psmeg- wag31(Mtb)-L34A-GFP-clone2 | 4 |
| CB2307 | mc$^{2}$155 L5:: pkk158-Psmeg- wag31(Mtb)-L34A-GFP-clone3 | 4 |
| CB2357 | mc$^{2}$155 Δwag31::hygR L5:: pKK158-Psmeg- wag31(Mtb)-wt giles::pGtetO-GlfT2-mRFP/ pNitRecET- clone1 | 5 |
| CB2358 | mc$^{2}$155 Δwag31::hygR L5:: pKK158-Psmeg- wag31(Mtb)-wt giles::pGtetO-GlfT2-mRFP/ pNitRecET- clone2 | 5 |
| CB2359 | mc$^{2}$155 Δwag31::hygR L5:: pKK158-Psmeg- wag31(Mtb)-wt giles::pGtetO-GlfT2-mRFP/ pNitRecET- clone3 | 5 |
| CB2360 | mc$^{2}$155 Δwag31::hygR L5:: pKK158-Pwag31- wag31(Mtb)-K20A giles::pGtetO-GlfT2-mRFP/ pNitRecET- clone1 | 5 |
| CB2361 | mc$^{2}$155 Δwag31::hygR L5:: pKK158-Pwag31- wag31(Mtb)-K20A giles::pGtetO-GlfT2-mRFP/ pNitRecET- clone2 | 5 |
| CB2362 | mc$^{2}$155 Δwag31::hygR L5:: pKK158-Pwag31- wag31(Mtb)-K20A giles::pGtetO-GlfT2-mRFP/ pNitRecET- clone3 | 5 |
| CB2363 | mc$^{2}$155 Δwag31::hygR L5:: pKK158-Pwag31- wag31(Mtb)-F255A giles::pGtetO-GlfT2-mRFP/ pNitRecET- clone1 | 5 |
| CB2364 | mc$^{2}$155 Δwag31::hygR L5:: pKK158-Pwag31- wag31(Mtb)-F255A giles::pGtetO-GlfT2-mRFP/ pNitRecET- clone2 | 5 |
| CB2365 | mc$^{2}$155 Δwag31::hygR L5:: pKK158-Pwag31- wag31(Mtb)-F255A giles::pGtetO-GlfT2-mRFP/ pNitRecET- clone3 | 5 |
| CB2366 | mc$^{2}$155 Δwag31::hygR L5:: pKK158-Pwag31- wag31(Mtb)-NQQR199-202AAAA giles::pGtetO-GlfT2-mRFP/ pNitRecET- clone1 | 5 |
| CB2367 | mc$^{2}$155 Δwag31::hygR L5: pKK158-Pwag31- wag31(Mtb)-NQQR199-202AAAA giles::pGtetO-GlfT2-mRFP/ pNitRecET- clone2 | 5 |
| CB2368 | mc$^{2}$155 Δwag31::hygR L5::pKK158-Pwag31- wag31(Mtb)-NQQR199-202AAAA giles::pGtetO-GlfT2-mRFP/ pNitRecET- clone3 | 5 |
| CB2369 | mc$^{2}$155 Δwag31::hygR L5:: pKK158-Pwag31- wag31(Mtb)-L34A giles::pGtetO-GlfT2-mRFP/ pNitRecET- clone1 | 5 |
| CB2370 | mc$^{2}$155 Δwag31::hygR L5: pKK158-Pwag31- wag31(Mtb)-L34A giles::pGtetO-GlfT2-mRFP/ pNitRecET- clone2 | 5 |
| CB2371 | mc$^{2}$155 Δwag31::hygR L5:: pKK158-Pwag31- wag31(Mtb)-L34A giles::pGtetO-GlfT2-mRFP/ pNitRecET- clone3 | 5 |
| CB2372 | mc$^{2}$155 Δwag31::hygR L5:: pKK158-Pwag31- wag31(Mtb)-D7A giles::pGtetO-GlfT2-mRFP/ pNitRecET- clone1 | 5 |
| CB2373 | mc$^{2}$155 Δwag31::hygR L5:: pKK158-Pwag31- wag31(Mtb)-D7A giles::pGtetO-GlfT2-mRFP/ pNitRecET- clone2 | 5 |

| | | |
|---|---|---|
| CB2374 | mc$^2$155 Δwag31::hygR L5:: pKK158-Pwag31- wag31(Mtb)-D7A giles::pGtetO-GlfT2-mRFP/ pNitRecET- clone3 | 5 |
| CB2325 | mc$^2$155 / ptetOR L5:: CT94 – wag31-GFP1-10 giles::pGtetO - wag31-GFP11-clone1 | 2, S7 |
| CB2384 | mc$^2$155 / ptetOR L5:: CT94 - wag31-GFP1-10 giles::pGtetO - wag31-GFP11-clone2 | 2, S7 |
| CB2385 | mc$^2$155 / ptetOR L5:: CT94 - wag31-GFP1-10 giles::pGtetO - wag31-GFP11-clone3 | 2, S7 |
| CB2386 | mc$^2$155 / ptetOR L5:: CT94 - wag31-GFP1-10 giles::pGtetO - wag31-GFP11-clone4 | 2, S7 |
| CB2333 | mc$^2$155 L5:: CT94 - wag31-GFP1-10 giles::pGtetO-mcherry2b-GFP11/ pTetOR-CRISPR Repeat-clone1 | S7 |
| CB2387 | mc$^2$155 L5:: CT94 - wag31-GFP1-10 giles::pGtetO-mcherry2b-GFP11/ pTetOR-CRISPR Repeat-clone2 | S7 |
| CB2388 | mc$^2$155 L5:: CT94 - wag31-GFP1-10 giles::pGtetO-mcherry2b-GFP11/ pTetOR-CRISPR Repeat-clone3 | S7 |
| CB2335 | mc$^2$155 L5:: CT94 -GFP1-10- wag31 giles::pGtetO-mcherry2b-GFP11/ pTetOR-CRISPR Repeat-clone1 | S7 |
| CB2389 | mc$^2$155 L5:: CT94 -GFP1-10- wag31 giles::pGtetO-mcherry2b-GFP11/ pTetOR-CRISPR Repeat-clone2 | S7 |
| CB2390 | mc$^2$155 L5:: CT94 -GFP1-10- wag31 giles::pGtetO-mcherry2b-GFP11/ pTetOR-CRISPR Repeat-clone3 | S7 |
| CB2337 | mc$^2$155 L5:: CT94 -GFP1-10- wag31 giles::pGtetO - wag31-GFP11/ pTetOR-CRISPR Repeat-clone1 | 2, S7 |
| CB2391 | mc$^2$155 L5:: CT94 -GFP1-10- wag31 giles::pGtetO - wag31-GFP11/ pTetOR-CRISPR Repeat-clone2 | 2, S7 |
| CB2392 | mc$^2$155 L5:: CT94 -GFP1-10-Wag31 giles::pGtetO - wag31-GFP11/ pTetOR-CRISPR Repeat-clone3 | 2, S7 |
| CB2336 | mc$^2$155 L5:: CT94 -GFP1-10- wag31 giles::pGtetO-GFP11-mcherry2b/ pTetOR-CRISPR Repeat | |
| CB2338 | mc$^2$155 L5:: CT94 -GFP1-10- wag31 giles::pGtetO-GFP11- wag31/ pTetOR-CRISPR Repeat | |
| CB2339 | mc$^2$155 L5:: CT94 - wag31-GFP1-10 giles::pGtetO-GFP11- wag31/ pTetOR-CRISPR Repeat | |
| CB2342 | mc$^2$155 L5:: pkk158-Psmeg- wag31(Mtb)-D7A-GFP-clone1 | 4 |
| CB2343 | mc$^2$155 L5:: pkk158-Psmeg- wag31(Mtb)-D7A-GFP-clone2 | 4 |
| CB2344 | mc$^2$155 L5:: pkk158-Psmeg- wag31(Mtb)-D7A-GFP-clone3 | 4 |
| CB2345 | mc$^2$155 L5:: pkk158-Psmeg- wag31(Mtb)-K20A-GFP-clone1 | 4 |
| CB2346 | mc$^2$155 L5:: pkk158-Psmeg- wag31(Mtb)-K20A-GFP-clone2 | 4 |
| CB2347 | mc$^2$155 L5:: pkk158-Psmeg- wag31(Mtb)-K20A-GFP-clone3 | 4 |
| CB2348 | mc$^2$155 L5:: pkk158-Psmeg- wag31(Mtb)-F255A-GFP-clone1 | 4 |
| CB2349 | mc$^2$155 L5:: pkk158-Psmeg- wag31(Mtb)-F255A-GFP-clone2 | 4 |
| CB2350 | mc$^2$155 L5:: pkk158-Psmeg- wag31(Mtb)-F255A-GFP-clone3 | 4 |

| | | |
|---|---|---|
| CB2398 | $mc^2$155 L5:: CT94 -wag31-GFP1-10 giles::pGtetO- wag31-D7A-GFP11/ pTetOR-CRISPR Repeat-clone1 | 2 |
| CB2399 | $mc^2$155 L5:: CT94 -Wag31-GFP1-10 giles::pGtetO- wag31-D7A-GFP11/ pTetOR-CRISPR Repeat-clone2 | 2 |
| CB2400 | $mc^2$155 L5:: CT94 -Wag31-GFP1-10 giles::pGtetO- wag31-D7A-GFP11/ pTetOR-CRISPR Repeat-clone3 | 2 |
| CB2401 | $mc^2$155 L5:: CT94 -Wag31-GFP1-10 giles::pGtetO- wag31-K20A-GFP11/ pTetOR-CRISPR Repeat-clone1 | 2 |
| CB2402 | $mc^2$155 L5:: CT94 -Wag31-GFP1-10 giles::pGtetO- wag31-K20A-GFP11/ pTetOR-CRISPR Repeat-clone2 | 2 |
| CB2403 | $mc^2$155 L5:: CT94 -Wag31-GFP1-10 giles::pGtetO- wag31-K20A-GFP11/ pTetOR-CRISPR Repeat-clone3 | 2 |
| CB2404 | $mc^2$155 L5:: CT94 -Wag31-GFP1-10 giles::pGtetO- wag31-L34A-GFP11/ pTetOR-CRISPR Repeat-clone1 | 2 |
| CB2405 | $mc^2$155 L5:: CT94 -Wag31-GFP1-10 giles::pGtetO- wag31-L34A-GFP11/ pTetOR-CRISPR Repeat-clone2 | 2 |
| CB2406 | $mc^2$155 L5:: CT94 -Wag31-GFP1-10 giles::pGtetO-wag31-L34A-GFP11/ pTetOR-CRISPR Repeat-clone3 | 2 |
| CB2407 | $mc^2$155 L5:: CT94 -Wag31-GFP1-10 giles::pGtetO- wag31-NQQR199-202AAAA-GFP11/ pTetOR-CRISPR Repeat-clone1 | 2 |
| CB2408 | $mc^2$155 L5:: CT94 -Wag31-GFP1-10 giles::pGtetO- wag31-NQQR199-202AAAA-GFP11/ pTetOR-CRISPR Repeat-clone2 | 2 |
| CB2409 | $mc^2$155 L5:: CT94 -Wag31-GFP1-10 giles::pGtetO- wag31-NQQR199-202AAAA-GFP11/ pTetOR-CRISPR Repeat-clone3 | 2 |
| CB2410 | $mc^2$155 L5:: CT94 -Wag31-GFP1-10 giles::pGtetO- wag31-F255A-GFP11/ pTetOR-CRISPR Repeat-clone1 | 2 |
| CB2411 | $mc^2$155 L5:: CT94 -Wag31-GFP1-10 giles::pGtetO-wag31-F255A-GFP11/ pTetOR-CRISPR Repeat-clone2 | 2 |
| CB2412 | $mc^2$155 L5:: CT94 -Wag31-GFP1-10 giles::pGtetO- wag31-F255A-GFP11/ pTetOR-CRISPR Repeat-clone3 | 2 |
| CB2413 | $mc^2$155 L5:: CT94 -GFP1-10-Wag31giles::pGtetO- wag31-D7A-GFP11/ pTetOR-CRISPR Repeat-clone1 | 2 |
| CB2414 | $mc^2$155 L5:: CT94 -GFP1-10-Wag31giles::pGtetO- wag31-D7A-GFP11/ pTetOR-CRISPR Repeat-clone2 | 2 |
| CB2415 | $mc^2$155 L5:: CT94 -GFP1-10-Wag31giles::pGtetO- wag31-D7A-GFP11/ pTetOR-CRISPR Repeat-clone3 | 2 |
| CB2416 | $mc^2$155 L5:: CT94 -GFP1-10-Wag31giles::pGtetO- wag31-K20A-GFP11/ pTetOR-CRISPR Repeat-clone1 | 2 |
| CB2417 | $mc^2$155 L5:: CT94 -GFP1-10-Wag31giles::pGtetO- wag31-K20A-GFP11/ pTetOR-CRISPR Repeat-clone2 | 2 |
| CB2418 | $mc^2$155 L5:: CT94 -GFP1-10-Wag31giles::pGtetO- wag31-K20A-GFP11/ pTetOR-CRISPR Repeat-clone3 | 2 |
| CB2419 | $mc^2$155 L5:: CT94 -GFP1-10-Wag31giles::pGtetO- wag31-L34A-GFP11/ pTetOR-CRISPR Repeat-clone1 | 2 |

| | | |
|---|---|---|
| CB2420 | mc$^2$155 L5:: CT94 -GFP1-10-Wag31giles::pGtetO- wag31-L34A-GFP11/ pTetOR-CRISPR Repeat-clone2 | 2 |
| CB2421 | mc$^2$155 L5:: CT94 -GFP1-10-Wag31giles::pGtetO- wag31-L34A-GFP11/ pTetOR-CRISPR Repeat-clone3 | 2 |
| CB2422 | mc$^2$155 L5:: CT94 -GFP1-10-Wag31giles::pGtetO- wag31-NQQR199-202-GFP11/ pTetOR-CRISPR Repeat-clone1 | 2 |
| CB2423 | mc$^2$155 L5:: CT94 -GFP1-10-Wag31giles::pGtetO- wag31-NQQR199-202-GFP11/ pTetOR-CRISPR Repeat-clone2 | 2 |
| CB2424 | mc$^2$155 L5:: CT94 -GFP1-10-Wag31giles::pGtetO- wag31-NQQR199-202-GFP11/ pTetOR-CRISPR Repeat-clone3 | 2 |
| CB2425 | mc$^2$155 L5:: CT94 -GFP1-10-Wag31giles::pGtetO- wag31-F255A-GFP11/ pTetOR-CRISPR Repeat-clone1 | 2 |
| CB2426 | mc$^2$155 L5:: CT94 -GFP1-10-Wag31giles::pGtetO- wag31-F255A-GFP11/ pTetOR-CRISPR Repeat-clone2 | 2 |
| CB2427 | mc$^2$155 L5:: CT94 -GFP1-10-Wag31giles::pGtetO- wag31-F255A-GFP11/ pTetOR-CRISPR Repeat-clone3 | 2 |
| CB2524 | mc2155 Δwag31::hygR L5::pKK158-Psmeg- wag31(Mtb)-wt/ pMEK-P21-murG-Venus-clone1 | 5 |
| CB2527 | mc2155 Δwag31::hygR L5::pKK158-Psmeg- wag31(Mtb)-wt/ pMEK-P21-murG-Venus-clone2 | 5 |
| CB2528 | mc2155 Δwag31::hygR L5::pKK158-Psmeg- wag31(Mtb)-wt/ pMEK-P21-murG-Venus-clone3 | 5 |
| CB2529 | mc2155 Δwag31::hygR L5::pKK158-Pwag31- wag31(Mtb)-K20A/ pMEK-P21-murG-Venus-clone1 | 5 |
| CB2530 | mc2155 Δwag31::hygR L5::pKK158-Pwag31- wag31(Mtb)-K20A/ pMEK-P21-murG-Venus- clone2 | 5 |
| CB2531 | mc2155 Δwag31::hygR L5::pKK158-Pwag31- wag31(Mtb)-K20A/ pMEK-P21-murG-Venus-clone3 | 5 |
| CB2532 | mc2155 Δwag31::hygR L5::pKK158-Pwag31- wag31(Mtb)-F255A/ pMEK-P21-murG-Venus- clone1 | 5 |
| CB2533 | mc2155 Δwag31::hygR L5::pKK158-Pwag31- wag31(Mtb)-F255A/ pMEK-P21-murG-Venus- clone2 | 5 |
| CB2534 | mc2155 Δwag31::hygR L5::pKK158-Pwag31- wag31(Mtb)-F255A/ pMEK-P21-murG-Venus- clone #3 | 5 |
| CB2535 | mc2155 Δwag31::hygR L5::pKK158-Pwag31- wag31(Mtb)-NQQR199-202-AAAA/ pMEK-P21-murG-Venus- clone1 | 5 |
| CB2536 | mc2155 Δwag31::hygR L5:pKK158-Pwag31- wag31(Mtb)-NQQR199-202-4AAAA/ pMEK-P21-murG-Venus-clone2 | 5 |
| CB2537 | mc2155 Δwag31::hygR L5::pKK158-Pwag31- wag31(Mtb)-NQQR199-202-AAAA/ pMEK-P21-murG-Venus-clone3 | 5 |
| CB2538 | mc2155 Δwag31::hygR L5::pKK158-Pwag31- wag31(Mtb)-L34A/ pMEK-P21-murG-Venus-clone1 | 5 |
| CB2539 | mc2155 Δwag31::hygR L5:pKK158-Pwag31- wag31(Mtb)-L34A/ pMEK-P21-murG-Venus-clone2 | 5 |

| | | |
|---|---|---|
| CB2540 | mc2155 ∆wag31::hygR L5::pKK158-Pwag31- wag31(Mtb)-L34A/ pMEK-P21-murG-Venus-clone3 | 5 |
| CB2541 | mc2155 ∆wag31::hygR L5::pKK158-Pwag31- wag31(Mtb)-D7A/ pMEK-P21-murG-Venus- clone1 | 5 |
| CB2542 | mc2155 ∆wag31::hygR L5:pKK158-Pwag31- wag31(Mtb)-D7A/ pMEK-P21-murG-Venus-clone2 | 5 |
| CB2543 | mc2155 ∆wag31::hygR L5::pKK158-Pwag31- wag31(Mtb)-D7A/ pMEK-P21-murG-Venus- clone3 | 5 |

**Table S2B. Plasmid list.**

| Strain | Genotype | Used in strain | |
|---|---|---|---|
| CB207 | DH5a/ pNit-RecET-SacB-gent | CB1821 | Sassetti Lab |
| CB1260 | DH5a/ pMEK-Ptb21-murG-Venus | CB2524, CB2527-43 | Xavier Meniche |
| CB1323 | DH5a/ pUAB200-pL5-gcn4-kanR | CB1613 | Adrie Steyn |
| CB1324 | DH5a/ pUAB300-groEl -ORF6-hygR | CB1612 | Adrie Steyn |
| CB1337 | DH5a/ pUAB400-pl5-groEl -ORF5-KanR | CB1612 | Adrie Steyn |
| CB1338 | DH5a/ pUAB100-HSp60-gcn4 dimerization domain-HygR | CB1613 | Adrie Steyn |
| CB1672 | Top10/ pUAB100- wag31(Mtb)-wt-dhfr1,2 | CB1704, CB1757, CB1769-73, CB1800-2, CB1843- 1854, CB1891- 1899, CB1900- 1905, CB2101-2117 | |
| CB1683 | XL1 Blue/ pKK158- Pwag31- wag31(Msmeg) | CB1821 | |
| CB1690 | Top10/ pUAB200-wag31(Mtb)-wt-dhfr3 | CB1704 | |
| CB1721 | XL1 Blue/ pUAB200- wag31(Mtb)-∆Ct 10 aa | CB1757 | |
| CB1759 | XL1 Blue/ pUAB200- wag31(Mtb)-PLLT2-5AAAA-dhfr3 | CB1769 | |
| CB1760 | XL1 Blue/ pUAB200- wag31(Mtb)-DVH7-9AAA-dhfr3 | CB1770 | |
| CB1761 | XL1 Blue/ pUAB200- wag31(Mtb)-N10A, V11A, F13A-dhfr3 | CB1771 | |
| CB1762 | XL1 Blue/ pUAB200- wag31(Mtb)-SKPP14-17AAAA-dhfr3 | CB1772 | |
| CB1763 | XL1 Blue/ pUAB200- wag31(Mtb)-IGKRI18-21AAAA)-dhfr3 | CB1773 | |

| | | |
|---|---|---|
| CB1789 | Top10/ pUAB200- wag31(Mtb)- RGKN257-60AAAA-dhfr3 | CB1800 |
| CB1790 | Top 10/ pUAB200- wag31(Mtb)- DQFN253-6AAAA -dhfr3 | CB1801 |
| CB1791 | Top 10/ pUAB200- wag31(Mtb)- GGF250-2AAA-dhfr3 | CB1802 |
| CB1827 | Top10/ pUAB200- wag31(Mtb)-N10A-dhfr3 | CB1843 |
| CB1806 | Top10/ pUAB200- wag31(Mtb)-F13A-dhfr3 | CB1844 |
| CB1807 | Top10/ pUAB200- wag31(Mtb)-S14A-dhfr3 | CB1845 |
| CB1808 | Top10/ pUAB200- wag31(Mtb)-K15A-dhfr3 | CB1846 |
| CB1828 | Top10/ pUAB200- wag31(Mtb)-K20A-dhfr3 | CB1847 |
| CB1829 | Top10/ pUAB200- wag31(Mtb)-R21A-dhfr3 | CB1848 |
| CB1830 | Top10/ pUAB200- wag31(Mtb)-KR20, 21AA-dhfr3 | CB1849 |
| CB1831 | Top10/ pUAB200- wag31(Mtb)-EDED 25-29 4A-dhfr3 | CB1850 |
| CB1832 | Top10/ pUAB200- wag31(Mtb)-FD 31,33 AA-dhfr3 | CB1851 |
| CB1833 | Top10/ pUAB200- wag31(Mtb)-ENE 36-38 3A-dhfr3 | CB1852 |
| CB1834 | Top10/ pUAB200- wag31(Mtb)-TRE 40-41-44 3A-dhfr3 | CB1853 |
| CB1859 | Top10/ pUAB200- wag31(Mtb)-N260A-dhfr3 | CB1893 |
| CB1860 | Top10/ pUAB200- wag31(Mtb)-K259A-dhfr3 | CB1894 |
| CB1861 | Top10/ pUAB200- wag31(Mtb)-R257A-dhfr3 | CB1895 |
| CB1862 | Top10/ pUAB200- wag31(Mtb)-N256A-dhfr3 | CB1896 |
| CB1863 | Top10/ pUAB200- wag31(Mtb)-F255A-dhfr3 | CB1897 |
| CB1864 | Top10/ pUAB200- wag31(Mtb)-Q254A-dhfr3 | CB1898 |
| CB1865 | Top10/ pUAB200- wag31(Mtb)-D253A-dhfr3 | CB1899 |
| CB1866 | Top10/ pUAB200- wag31(Mtb)-F252A-dhfr3 | CB1900 |
| CB1867 | Top10/ pUAB200- wag31(Mtb)-D248A-dhfr3 | CB1901 |
| CB1868 | Top10/ pUAB200- wag31(Mtb)-DQE 57-9AAA-dhfr3 | CB1902 |
| CB1869 | Top10/ pUAB200- wag31(Mtb)-NEEQ 96-9AAAA-dhfr3 | CB1903 |
| CB1870 | Top10/ pUAB200- wag31(Mtb)-ESDK 124-127-4AAAA-dhfr3 | CB1904 |
| CB1871 | Top10/ pUAB200- wag31(Mtb)-ERH 141,143,144AAA-dhfr3 | CB1905 |
| CB1872 | Top10/ pUAB200- wag31(Mtb)-E45R-dhfr3 | CB1891 |
| CB1873 | Top10/ pUAB200- wag31(Mtb)-E45D-dhfr3 | CB1892 |
| CB1991 | Top10/ pUAB200- wag31(Mtb)-E27A-dhfr3 | CB2101 |
| CB1992 | Top10/ pUAB200- wag31(Mtb)-E38A-dhfr3 | CB2102 |

| | | |
|---|---|---|
| CB1993 | Top10/ pUAB200- wag31(Mtb)-L34A-dhfr3 | CB2103 |
| CB1994 | Top10/ pUAB200- wag31(Mtb)-R41A-dhfr3 | CB2104 |
| CB1995 | Top10/ pUAB200- wag31(Mtb)-NSD46-8AAA-dhfr3 | CB2105 |
| CB1996 | Top10/ pUAB200- wag31(Mtb)-ΔNSD46-48)-dhfr3 | CB2106 |
| CB1997 | Top10/ pUAB200- wag31(Mtb)-R153,155AA-dhfr3 | CB2107 |
| CB1998 | Top10/ pUAB200- wag31(Mtb)-R156,157AA-dhfr3 | CB2108 |
| CB1999 | Top10/ pUAB200- wag31(Mtb)-QSRS166-9AAAA-dhfr3 | CB2109 |
| CB2000 | Top10/ pUAB200- wag31(Mtb)-RQQ174,5,7AAA-dhfr3 | CB2110 |
| CB2001 | Top10/ pUAB200- wag31(Mtb)-DER185,7,8AAA-dhfr3 | CB2111 |
| CB2002 | Top10/ pUAB200- wag31(Mtb)-NQQR199-202AAAA-dhfr3 | CB2112 |
| CB2003 | Top10/ pUAB200- wag31(Mtb)-EQR210-11-13AAA-dhfr3 | CB2113 |
| CB2004 | Top10/ pUAB200- wag31(Mtb)-RRK220,2,4AAA-dhfr3 | CB2114 |
| CB2005 | Top10/ pUAB200- wag31(Mtb)-QEE230,2,3AAA-dhfr3 | CB2115 |
| CB2006 | Top10/ pUAB200- wag31(Mtb)-D244A-dhfr3 | CB2116 |
| CB2007 | Top10/ pUAB200- wag31(Mtb)-N246A-dhfr3 | CB2117 |
| CB1909 | Top10/ pKK158 - Pwag31-wag31(Mtb)-wt | CB1927-9 |
| CB1938 | Top10/ pkk158- Pwag31- wag31(Mtb)-N10A | CB1966-8 |
| CB1939 | Top10/ pkk158- Pwag31- wag31(Mtb)-S14A | CB2025-7 |
| CB1941 | Top10/ pkk158-Pwag31- wag31(Mtb)-K20A | CB2028-30 |
| CB1942 | Top10/ pkk158-Pwag31- wag31(Mtb)--R21A | CB2031, 2 |
| CB1943 | Top10/ pkk158-Pwag31- wag31(Mtb)-KR20,21AA | CB1969-71 |
| CB1944 | Top10/ pkk158-Pwag31- wag31(Mtb)-ENE36-38AAA | CB1985-7 |
| CB1945 | Top10/ pkk158-Pwag31- wag31(Mtb)-E45R | CB2057-9 |
| CB1946 | Top10/ pkk158-Pwag31- wag31(Mtb)-E45D | CB2060-2 |
| CB1947 | Top10/ pkk158-Pwag31- wag31(Mtb)-N260A | CB2063-5 |
| CB1948 | Top10/ pkk158-Pwag31- wag31(Mtb)-K259A | CB2066-8 |
| CB1949 | Top10/ pkk158-Pwag31- wag31(Mtb)-R257A | CB1988-90 |
| CB1950 | Top10/ pkk158-Pwag31- wag31(Mtb)-N256A | CB2069-71 |
| CB1951 | Top10/ pkk158-Pwag31- wag31(Mtb)-F255A | CB2033-5 |
| CB1952 | Top10/ pkk158-Pwag31- wag31(Mtb)-Q254A | CB2041-3 |

| | | |
|---|---|---|
| CB1953 | Top10/ pkk158-Pwag31- wag31(Mtb)-D253A | CB2038-40 |
| CB1954 | Top10/ pkk158-Pwag31- wag31(Mtb)-F252A | CB2072- 4 |
| CB1955 | Top10/ pkk158-Pwag31- wag31(Mtb)-D248A | CB2087- 9 |
| CB1980 | Top10/ pkk158-Pwag31- wag31(Mtb)-DQFN253-6AAAA | CB2130-2 |
| CB1981 | Top10/ pkk158-Pwag31- wag31(Mtb)-GGF250-2AAA | CB2133-5 |
| CB1984 | Top10/ pkk158-Pwag31- wag31(Mtb)-ΔCt-10aa | CB2075-7 |
| CB2047 | Top10/ pkk158-Pwag31- wag31(Mtb)-DQE57-9AAA | CB2090-2 |
| CB2048 | Top10/ pkk158-Pwag31- wag31(Mtb)-NEEQ96-9AAAA | CB2093-5 |
| CB2049 | Top10/ pkk158-Pwag31-wag31 (ESDK 124-1274A) | CB2096-8 |
| CB2154 | Top10/ pkk158-Pwag31- wag31(Mtb)-E38 | CB2198-2200 |
| CB2155 | Top10/ pkk158-Pwag31- wag31(Mtb)-L34A | CB2195-7 |
| CB2157 | Top10/ pkk158-Pwag31- wag31(Mtb)-NSD46-8AAA | CB2201-3 |
| CB2159 | Top10/ pkk158-Pwag31- wag31(Mtb)-ER153,155AA | CB2177-9 |
| CB2160 | Top10/ pkk158-Pwag31- wag31(Mtb)-QR156,157AA | CB2204-6 |
| CB2161 | Top10/ pkk158-Pwag31- wag31(Mtb)-QSRS166-9AAAA | CB2180-2 |
| CB2162 | Top10/ pkk158-Pwag31- wag31(Mtb)-RQQ174,5,7AAA | CB2234-6 |
| CB2163 | Top10/ pkk158-Pwag31- wag31(Mtb)-DER185,7,8AAA | CB2207-9 |
| CB2164 | Top10/ pkk158-Pwag31- wag31(Mtb)-NQQR199-202AAAA | CB2183-5 |
| CB2167 | Top10/ pkk158-Pwag31- wag31(Mtb)-QEE230,2,3AAA | CB2186-8 |
| CB2168 | Top10/ pkk158-Pwag31- wag31(Mtb)-D244A | CB2210-12 |
| CB2169 | Top10/ pkk158-Pwag31- wag31(Mtb)-N246A | CB2213-5 |
| CB2223 | Top10/ pkk158-Pwag31- wag31(Mtb)-T4A | CB2238 |
| CB2224 | Top10/ pkk158-Pwag31- wag31(Mtb)-P5A | CB2269-70 |
| CB2225 | Top10/ pkk158-Pwag31- wag31(Mtb)-D7A | CB2271-3 |
| CB2226 | Top10/ pkk158-Pwag31- wag31(Mtb)-H9A | CB2231-3 |
| CB2227 | Top10/ pkk158-Pwag31- wag31(Mtb)-wt-GFP | CB2228-30 |
| CB2279 | Top10/ pkk158-Pwag31- wag31(Mtb)NQQR199-202AAAA-GFP | CB2296-8 |
| CB2284 | Top10/ pkk158-Pwag31- wag31(Mtb)-L34A-GFP | CB2305-7 |
| CB2319 | Top10/ pGtetO-GlfT2-mRFP-ZeoR | CB2357-74 |

| | | |
|---|---|---|
| CB2320 | Top 10/ pGtetO - wag31-GFP11 | CB2325, CB2337, CB2384-6, CB2391-2 |
| CB2326 | Top10/ pkk158-Pwag31- wag31(Mtb)-D7A-GFP | CB2342-4 |
| CB2327 | Top10/ pGtetO-mcherry2b-GFP11 | CB2333, CB2335, CB2387-90 |
| CB2328 | Top10/ pGtetO-GFP11-mcherry | CB2336 |
| CB2329 | Top10/ pGtetO-GFP11-wag31 | CB2338-9 |
| CB2330 | Top10/ CT94-GFP1-10-wag31 | CB2337, CB2391-2 |
| CB2340 | Top10/ pkk158-Pwag31- wag31(Mtb)-K20A-GFP | CB2345-7 |
| CB2341 | Top10/ pkk158-Pwag31- wag31(Mtb)-F255A-GFP | CB2348-50 |
| CB2393 | Top10/ pGtetO- wag31(Mtb)-D7A-GFP11 | CB2398-400 CB2413-5 |
| CB2394 | Top10/ pGtetO- wag31(Mtb)-K20A-GFP11 | CB2401-3, CB2416-8 |
| CB2395 | Top10/ pGtetO- wag31(Mtb)-L34A-GFP11 | CB2404-6, CB2419-21 |
| CB2396 | Top10/ pGtetO- wag31(Mtb)-NQQR199-202AAAA-GFP11 | CB2407-9, CB2422-4 |
| CB2397 | Top10/ pGtetO- wag31(Mtb)-F255A-GFP11 | CB2410-12 CB2425-7 |

**Table S2C. Primer list.**

| Strain | Genotype | Primers |
|---|---|---|
| CB1672 | Top10/ pUAB100- wag31(Mtb)-wt-dhfr1,2 | CAATGGCCAAGACAATTGCGGATCCgatgccgcttacacctgccgacgtccacaat<br>CCACCACCTCCAGAGCCACCGCCACCATCGATgttcttgccccggttgaattgatcgaag |
| CB1690 | Top10/ pUAB200 - wag31(Mtb)-wt-dhfr3 | AGGAATCACTTCGCAATGGCCAAGACAATTGcgatgccgcttacacctgc<br>CCACCACCTCCAGAGCCACCGCCACCATCGATgttcttgccccggttgaattgatcgaag |
| CB1721 | XL1 Blue/ pUAB200- wag31(Mtb)-ΔCt-10 aa | AGGAATCACTTCGCAATGGCCAAGACAATTGcgatgccgcttacacctgc<br>ACCACCTCCAGAGCCACCGCCACCATCGATaccggcatccgcattggaatcgaccggcg |
| CB1759 | XL1 Blue/ pUAB200- wag31(Mtb)-PLTP2-5AAAA-dhfr3 | GCCCGTCGTCGCAGCGAGTGGCAGCGAGGACAACTTGAGC<br>gaacgccacattgtggacgtcggcagctgcggcagccatcgCAATTGTCTTGGCCATTG<br>ATGGCCAAGACAATTGcgatggctgccgcagctgccgacgtccacaatgtggcgtt<br>CCACCACCTCCAGAGCCACCGCCACCATCGATgttcttgccccggttgaattgatcgaag |

| | | |
|---|---|---|
| CB1760 | XL1 Blue/ pUAB200- wag31(Mtb) - DVH7-9AAA-dhfr3 | GCCCGTCGTCGCAGCGAGTGGCAGCGAGGACA ACTTGAGC<br>gcttactgaacgccacattagccgcagcggcaggtgtaagcggcatc gCAATTGTCTTG<br>AAGACAATTGcgatgccgcttacacctgccgctgcggctaatgtg gcgttcagtaagcc<br>CCACCACCTCCAGAGCCACCGCCACCATCGATgt tcttgccccggttgaattgatcgaag |
| CB1761 | XL1 Blue/ pUAB200- wag31(Mtb)- N10, V11, F13AAA-dhfr3 | GCCCGTCGTCGCAGCGAGTGGCAGCGAGGACA ACTTGAGC<br>cgtttgccgataggcggcttactcgccgcagctgcgtggacgtcggca ggtgtaagcgg<br>gcttacacctgccgacgtccacgcagctgcggcgagtaagccgcctat cggcaaac<br>CCACCACCTCCAGAGCCACCGCCACCATCGATgt tcttgccccggttgaattgatcgaag |
| CB1762 | XL1 Blue/ pUAB200- wag31(Mtb)- SKPP14-17AAAA-dhfr3 | GCCCGTCGTCGCAGCGAGTGGCAGCGAGGACA ACTTGAGC<br>gttgtacccacgtttgccgattgccgcagccgcgaacgccacattgtgg acgtcggcag<br>ccgacgtccacaatgtggcgttcgcggctgcggcaatcggcaaacgt gggtacaac<br>CCACCACCTCCAGAGCCACCGCCACCATCGATgt tcttgccccggttgaattgatcgaag |
| CB1763 | XL1 Blue/ pUAB200- wag31(Mtb) - IGKR18-21AAAA-dhfr3 | GCCCGTCGTCGCAGCGAGTGGCAGCGAGGACA ACTTGAGC<br>cctcatcttcgttgtacccagctgcagccgcaggcggcttactgaacgc cacattgtg<br>caatgtggcgttcagtaagccgcctgcggctgcagctgggtacaacga agatgaggtc<br>CCACCACCTCCAGAGCCACCGCCACCATCGATgt tcttgccccggttgaattgatcgaag |
| CB1789 | Top10/ pUAB200- wag31(Mtb) - RGKN257-60AAAA-dhfr3 | ggcttcgatcaattcaacgcggcagctgcgATCGATGGTGGC GGTGGCTCTGGAGGTGG<br>TTATACTTGATGCCTTTTTCCTCCT<br>AGGAATCACTTCGCAATGGCCAAGACAATTGcgat gccgcttacacctgc<br>CAGAGCCACCGCCACCATCGATcgcagctgccgcgttg aattgatcgaagccacc |
| CB1790 | Top 10/ pUAB200- wag31(Mtb)- DQFN253-6AAAA -dhfr3 | atgcggatgccggtggcttcgcagctgcagctcggggcaagaacAT CGATGGTGGCG<br>TTATACTTGATGCCTTTTTCCTCCT<br>AGGAATCACTTCGCAATGGCCAAGACAATTGcgat gccgcttacacctgc<br>CGCCACCATCGATgttcttgccccgagctgcagctgcgaagc caccggcatccgcattg |

| | | |
|---|---|---|
| CB1791 | Top 10/ pUAB200- wag31(Mtb)-GGF250-2AAA-dhfr3 | ccggtcgattccaatgcggatgccgcagcggcagatcaattcaaccggggcaaga<br>TTATACTTGATGCCTTTTTCCTCCT<br>AGGAATCACTTCGCAATGGCCAAGACAATTGcgatgccgcttacacctgc<br>Cttgccccggttgaattgatctgccgctgcggcatccgcattggaatcgaccgg |
| CB1827 | Top10/ pUAB200- wag31(Mtb)-N10A-dhfr3 | GCCCGTCGTCGCAGCGAGTGGCAGCGAGGACAACTTGAGC<br>cgtttgccgataggcggcttactgaacgccacagcgtggacgtcggcaggtgtaagcgg<br>gcttacacctgccgacgtccacgctgtggcgttcagtaagccgcctatcggcaaac<br>CCACCACCTCCAGAGCCACCGCCACCATCGATgttcttgccccggttgaattgatcgaag |
| CB1806 | Top10/ pUAB200- wag31(Mtb)-F13A-dhfr3 | GCCCGTCGTCGCAGCGAGTGGCAGCGAGGACAACTTGAGC<br>cgtttgccgataggcggcttactcgccgccacattgtggacgtcggcaggtgtaagcgg<br>cttacacctgccgacgtccacaatgtggcggcgagtaagccgcctatcggcaaacgtg<br>CCACCACCTCCAGAGCCACCGCCACCATCGATgttcttgccccggttgaattgatcgaag |
| CB1807 | Top10/ pUAB200- wag31(Mtb)-S14A-dhfr3 | GCCCGTCGTCGCAGCGAGTGGCAGCGAGGACAACTTGAGC<br>gttgtacccacgtttgccgataggcggcttcgcgaacgccacattgtggacgtcggcag<br>ccgacgtccacaatgtggcgttcgcgaagccgcctatcggcaaacgtgggtacaac<br>CCACCACCTCCAGAGCCACCGCCACCATCGATgttcttgccccggttgaattgatcgaag |
| CB1808 | Top10/ pUAB200- wag31(Mtb)-K15A-dhfr3 | GCCCGTCGTCGCAGCGAGTGGCAGCGAGGACAACTTGAGC<br>gttgtacccacgtttgccgataggcggagcactgaacgccacattgtggacgtcggcag<br>ccgacgtccacaatgtggcgttcagtgctccgcctatcggcaaacgtgggtacaac<br>CCACCACCTCCAGAGCCACCGCCACCATCGATgttcttgccccggttgaattgatcgaag |
| CB1828 | Top10/ pUAB200- wag31(Mtb)-K20A-dhfr3 | GCCCGTCGTCGCAGCGAGTGGCAGCGAGGACAACTTGAGC<br>gacctcatcttcgttgtacccacgtgcgccgataggcggcttactgaacgccacattg<br>caatgtggcgttcagtaagccgcctatcggcgcacgtgggtacaacgaagatgaggtc |

| | | |
|---|---|---|
| | | CCACCACCTCCAGAGCCACCGCCACCATCGATgt tcttgccccggttgaattgatcgaag |
| CB1829 | Top10/ pUAB200- wag31(Mtb)-R21A-dhfr3 | GCCCGTCGTCGCAGCGAGTGGCAGCGAGGACA ACTTGAGC gacctcatcttcgttgtacccagctttgccgataggcggcttactgaacg ccacattg caatgtggcgttcagtaagccgcctatcggcaaagctgggtacaacg aagatgaggtc CCACCACCTCCAGAGCCACCGCCACCATCGATgt tcttgccccggttgaattgatcgaag |
| CB1830 | Top10/ pUAB200- wag31(Mtb)-KR20, 21AA-dhfr3 | GCCCGTCGTCGCAGCGAGTGGCAGCGAGGACA ACTTGAGC cctcatcttcgttgtacccagctgcgccgataggcggcttactgaacgcc acattgtg caatgtggcgttcagtaagccgcctatcggcgcagctgggtacaacga agatgaggtc CCACCACCTCCAGAGCCACCGCCACCATCGATgt tcttgccccggttgaattgatcgaag |
| CB1831 | Top10/ pUAB200- wag31(Mtb)-EDED 25-29AAAA-dhfr3 | GCCCGTCGTCGCAGCGAGTGGCAGCGAGGACA ACTTGAGC cgttttccaccaggtcgaggaaggccgcgactgcagccgcgttgtacc cacgtttgccg cggcaaacgtgggtacaacgcggctgcagtcgcggccttcctcgacc tggtggaaaac CCACCACCTCCAGAGCCACCGCCACCATCGATgt tcttgccccggttgaattgatcgaag |
| CB1832 | Top10/ pUAB200- wag31(Mtb) -FD 31,33AA-dhfr3 | GCCCGTCGTCGCAGCGAGTGGCAGCGAGGACA ACTTGAGC cgggtcagctcgttttccaccagtgcgagtgcggcgtcgacctcatcttc gttgtac gtacaacgaagatgaggtcgacgccgcactcgcactggtggaaaac gagctgaccc CCACCACCTCCAGAGCCACCGCCACCATCGATgt tcttgccccggttgaattgatcgaag |
| CB1833 | Top10/ pUAB200- wag31(Mtb)-ENE 36-38AAA-dhfr3 | GCCCGTCGTCGCAGCGAGTGGCAGCGAGGACA ACTTGAGC agttctcttcgatcaggcgggtcagagctgcagccaccaggtcgagga aggcgtcg cgacgccttcctcgacctggtggctgcagctctgacccgcctgatcgaa gagaactc CCACCACCTCCAGAGCCACCGCCACCATCGATgt tcttgccccggttgaattgatcgaag |
| CB1834 | Top10/ pUAB200- wag31(Mtb)-TRE40-41-44AAA-dhfr3 | GCCCGTCGTCGCAGCGAGTGGCAGCGAGGACA ACTTGAGC ctctgacgcagatcggagttctctgcgatcagagccgccagctcgttttc caccaggtcg |

| | | |
|---|---|---|
| | | gacctggtggaaaacgagctggcggctctgatcgcagagaactccg<br>atctgcgtcagag<br>CCACCACCTCCAGAGCCACCGCCACCATCGATgt<br>tcttgccccggttgaattgatcgaag |
| CB1859 | Top10/ pUAB200- wag31(Mtb) -<br>N260A-dhfr3 | AGGAATCACTTCGCAATGGCCAAGACAATTGcgat<br>gccgcttacacctgc<br>CCACCTCCAGAGCCACCGCCACCATCGATcgcctt<br>gccccggttgaattgatcgaagcc |
| CB1860 | Top10/ pUAB200- wag31(Mtb)-<br>K259A-dhfr3 | AGGAATCACTTCGCAATGGCCAAGACAATTGcgat<br>gccgcttacacctgc<br>CACCTCCAGAGCCACCGCCACCATCGATgttagcg<br>ccccggttgaattgatcgaag |
| CB1861 | Top10/ pUAB200- wag31(Mtb)-<br>R257A-dhfr3 | AGGAATCACTTCGCAATGGCCAAGACAATTGcgat<br>gccgcttacacctgc<br>CTCCAGAGCCACCGCCACCATCGATgttcttgcccgcg<br>ttgaattgatcgaagccacc |
| CB1862 | Top10/ pUAB200- wag31(Mtb)-<br>N256A-dhfr3 | AGGAATCACTTCGCAATGGCCAAGACAATTGcgat<br>gccgcttacacctgc<br>AGAGCCACCGCCACCATCGATgttcttgccccgagcgaa<br>ttgatcgaagccaccggc<br>ggatgccggtggcttcgatcaattcgctcggggcaagaacATCGA<br>TGGTGGCGGTG<br>CCTTTTTCCTCCTGGACCTCAGAGAGGACGCCT<br>GGGTATT |
| CB1863 | Top10/ pUAB200- wag31(Mtb)-<br>F255A-dhfr3 | AGGAATCACTTCGCAATGGCCAAGACAATTGcgat<br>gccgcttacacctgc<br>CCACCGCCACCATCGATgttcttgccccggttagcttgatcg<br>aagccaccggcatccgc<br>atgccggtggcttcgatcaagctaaccggggcaagaacATCGAT<br>GGTGGCGGTGG<br>CCTTTTTCCTCCTGGACCTCAGAGAGGACGCCT<br>GGGTATT |
| CB1864 | Top10/ pUAB200- wag31(Mtb)-<br>Q254A-dhfr3 | AGGAATCACTTCGCAATGGCCAAGACAATTGcgat<br>gccgcttacacctgc<br>CGCCACCATCGATgttcttgccccggttgaacgcatcgaagcc<br>accggcatccgcattg<br>gcggatgccggtggcttcgatgcgttcaaccggggcaagaacATC<br>GATGGTGG<br>CCTTTTTCCTCCTGGACCTCAGAGAGGACGCCT<br>GGGTATT |
| CB1865 | Top10/ pUAB200- wag31(Mtb)-<br>D253A-dhfr3 | AGGAATCACTTCGCAATGGCCAAGACAATTGcgat<br>gccgcttacacctgc<br>gttcttgccccggttgaattgagcgaagccaccggcatccgcattggaa<br>tc<br>ccaatgcggatgccggtggcttcgctcaattcaaccggggcaagaac<br>ATCGATGGTGGC |

| | | |
|---|---|---|
| | | CCTTTTTCCTCCTGGACCTCAGAGAGGACGCCT GGGTATT |
| CB1866 | Top10/ pUAB200- wag31(Mtb)-F252A-dhfr3 | AGGAATCACTTCGCAATGGCCAAGACAATTGcgat gccgcttacacctgc<br>gttcttgccccggttgaattgagcgaagccaccggcatccgcattggaa tc<br>ccaatgcggatgccggtggcttcgctcaattcaaccggggcaagaac ATCGATGGTGGC<br>CCTTTTTCCTCCTGGACCTCAGAGAGGACGCCT GGGTATT |
| CB1867 | Top10/ pUAB200- wag31(Mtb)-D248A-dhfr3 | AGGAATCACTTCGCAATGGCCAAGACAATTGcgat gccgcttacacctgc<br>gttgaattgatcgaagccaccggcagccgcattggaatcgaccggcg ccgccgatc<br>ggcggcgccggtcgattccaatgcggctgccggtggcttcgatcaattc aac<br>CCTTTTTCCTCCTGGACCTCAGAGAGGACGCCT GGGTATT |
| CB1868 | Top10/ pUAB200- wag31(Mtb)-DQE57-9AAA-dhfr3 | cgtcagaggatcaacgagctggctgcagctctcgccgcgggcggcg gtgccggc<br>CCACCACCTCCAGAGCCACCGCCACCATCGATgt tcttgccccggttgaattgatcgaag<br>AGGAATCACTTCGCAATGGCCAAGACAATTGcgat gccgcttacacctgc<br>Gccggcaccgccgcccgcggcgagagctgcagccagctcgttgatc ctctgacgcag |
| CB1869 | Top10/ pUAB200- wag31(Mtb)-NEEQ 96-9AAAA-dhfr3 | ggcggcggtctcggcggggatggctgcagctgcagccctgaaggcg gcgcgagtgctg<br>CCACCACCTCCAGAGCCACCGCCACCATCGATgt tcttgccccggttgaattgatcgaag<br>AGGAATCACTTCGCAATGGCCAAGACAATTGcgat gccgcttacacctgc<br>Cagcactcgcgccgccttcagggctgcagctgcagccatccccgccg agaccgccgcc |
| CB1870 | Top10/ pUAB200- wag31(Mtb)-ESDK124-127AAAA-dhfr3 | gaccggcttacaaacaccgccaaagccgctgcagctgctatgctggc cgatgcccgtgc<br>CCACCACCTCCAGAGCCACCGCCACCATCGATgt tcttgccccggttgaattgatcgaag<br>AGGAATCACTTCGCAATGGCCAAGACAATTGcgat gccgcttacacctgc<br>Gcacgggcatcggccagcatagcagctgcagcggctttggcggtgttt gtaagccggtc |
| CB1871 | Top10/ pUAB200- wag31(Mtb)-ERH 141,143,144AAA-dhfr3 | gccaatgcggagcagatcctcggtgcagccgctgctaccgccgacgc cacggtcgccg<br>CCACCACCTCCAGAGCCACCGCCACCATCGATgt tcttgccccggttgaattgatcgaag |

| | | |
|---|---|---|
| | | AGGAATCACTTCGCAATGGCCAAGACAATTGcgatgccgcttacacctgc<br>Gcacgggcatcggccagcatagcagctgcagcggctttggcggtgtttgtaagccggtc |
| CB1872 | Top10/ pUAB200- wag31(Mtb)-E45R-dhfr3 | CCACCACCTCCAGAGCCACCGCCACCATCGATgttcttgccccggttgaattgatcgaag<br>cgagctgacccgcctgatcgaagacaactccgatctgcgtcagaggatcaacg<br>GCCCGTCGTCGCAGCGAGTGGCAGCGAGGACAACTTGAGC<br>Cgttgatcctctgacgcagatcggagttgtcttcgatcaggcgggtcagctcg |
| CB1873 | Top10/ pUAB200- wag31(Mtb)-E45D-dhfr3 | CCACCACCTCCAGAGCCACCGCCACCATCGATgttcttgccccggttgaattgatcgaag<br>cgagctgacccgcctgatcgaacgtaactccgatctgcgtcagaggatcaacg<br>GCCCGTCGTCGCAGCGAGTGGCAGCGAGGACAACTTGAGC<br>Cgttgatcctctgacgcagatcggagttacgttcgatcaggcgggtcagctcg |
| CB1991 | Top10/ pUAB200- wag31(Mtb)-E27A-dhfr3 | ggcaaacgtgggtacaacgaagatgccgtcgacgccttcctcgacctggtg<br>CCACCACCTCCAGAGCCACCGCCACCATCGATgttcttgccccggttgaattgatcgaag<br>ccaccaggtcgaggaaggcgtcgacggcatcttcgttgtacccacgtttgccg<br>GCCCGTCGTCGCAGCGAGTGGCAGCGAGGACAACTTGAGC |
| CB1992 | Top10/ pUAB200- wag31(Mtb)-E38A-dhfr3 | gccttcctcgacctggtggaaaacgcgctgacccgcctgatcgaagagaactcc<br>CCACCACCTCCAGAGCCACCGCCACCATCGATgttcttgccccggttgaattgatcgaag<br>ggagttctcttcgatcaggcgggtcagcgcgttttccaccaggtcgaggaaggcgt<br>GCCCGTCGTCGCAGCGAGTGGCAGCGAGGACAACTTGAGC |
| CB1993 | Top10/ pUAB200- wag31(Mtb)-L34A-dhfr3 | gaagatgaggtcgacgccttcctcgacgcggtggaaaacgagctgacccgcctg<br>CCACCACCTCCAGAGCCACCGCCACCATCGATgttcttgccccggttgaattgatcgaag<br>caggcgggtcagctcgttttccaccgcgtcgaggaaggcgtcgacctcatcttc<br>GCCCGTCGTCGCAGCGAGTGGCAGCGAGGACAACTTGAGC |
| CB1994 | Top10/ pUAB200- wag31(Mtb)-R41A-dhfr3 | gacctggtggaaaacgagctgaccgcgctgatcgaagagaactccgatctg |

| | | |
|---|---|---|
| | | CCACCACCTCCAGAGCCACCGCCACCATCGATgt tcttgccccggttgaattgatcgaag cagatcggagttctcttcgatcagcgcggtcagctcgttttccaccaggt c GCCCGTCGTCGCAGCGAGTGGCAGCGAGGACA ACTTGAGC |
| CB1995 | Top10/ pUAB200- wag31(Mtb)-NSD46-8AAA-dhfr3 | ctgacccgcctgatcgaagaggccgcggcactgcgtcagaggatca acgagctgg CCACCACCTCCAGAGCCACCGCCACCATCGATgt tcttgccccggttgaattgatcgaag gctcgttgatcctctgacgcagtgccgcggcctcttcgatcaggcgggt cagctc GCCCGTCGTCGCAGCGAGTGGCAGCGAGGACA ACTTGAGC |
| CB1996 | Top10/ pUAB200- wag31(Mtb)-ΔNSD46-48-dhfr3 | ccagctcgttgatcctctgacgcagctcttcgatcaggcgggtcagctc g CCACCACCTCCAGAGCCACCGCCACCATCGATgt tcttgccccggttgaattgatcgaag ctgacccgcctgatcgaagagctgcgtcagaggatcaacgagctgga tca GCCCGTCGTCGCAGCGAGTGGCAGCGAGGACA ACTTGAGC |
| CB1997 | Top10/ pUAB200- wag31(Mtb)-ER153,155AA-dhfr3 | gacacaccgccgacgccacggtcgccgcggccgcgcagcgtgccg atgccatgctgg CCACCACCTCCAGAGCCACCGCCACCATCGATgt tcttgccccggttgaattgatcgaag ccagcatggcatcggcacgctgcgcggccgcggcgaccgtggcgtc ggcggtgtgtc AGGAATCACTTCGCAATGGCCAAGACAATTGcgat gccgcttacacctgc |
| CB1998 | Top10/ pUAB200- wag31(Mtb)-QR156,157AA-dhfr3 | AGGAATCACTTCGCAATGGCCAAGACAATTGcgat gccgcttacacctgc gggcatcggccagcatggcatcggccgccgcgcgggcctcggcgac cgtggcgtcggc gccgacgccacggtcgccgaggcccgcgcggcggccgatgccatg ctggccgatgccc CCACCACCTCCAGAGCCACCGCCACCATCGATgt tcttgccccggttgaattgatcgaag |
| CB1999 | Top10 / pUAB200- wag31(Mtb)-QSRS166-9-4AAAA-dhfr3 | AGGAATCACTTCGCAATGGCCAAGACAATTGcgat gccgcttacacctgc cctgcgcctggcgcaactgggcctcggccgccgccgcggcatcggc cagcatggcatc gatgccatgctggccgatgccgcggcggcggccgaggcccagttgc gccaggcgcagg CCACCACCTCCAGAGCCACCGCCACCATCGATgt tcttgccccggttgaattgatcgaag |

| | | |
|---|---|---|
| CB2000 | Top10/ pUAB200- wag31(Mtb)-RQQ174,5,7AAA-dhfr3 | AGGAATCACTTCGCAATGGCCAAGACAATTGcgatgccgcttacacctgc<br>ctgtaaggcatcggccttctcggccgccgccgccaactgggcctcggatcgggattggg<br>ccaatcccgatccgaggcccagttggcggcggcggccgagaaggccgatgccttacagg<br>CCACCACCTCCAGAGCCACCGCCACCATCGATgttcttgccccggttgaattgatcgaag |
| CB2001 | Top10/ pUAB200- wag31(Mtb)-DER185,7,8AAA-dhfr3 | AGGAATCACTTCGCAATGGCCAAGACAATTGcgatgccgcttacacctgc<br>gttcccatgatctcggagtgcttcgcggccgccgcggcctgtaaggcatcggccttctc<br>gagaaggccgatgccttacaggccgcggcggccgcgaagcactccgagatcatgggaac<br>CCACCACCTCCAGAGCCACCGCCACCATCGATgttcttgccccggttgaattgatcgaag |
| CB2002 | Top10/ pUAB200- wag31(Mtb)-NQQR199-4AAAA-dhfr3 | AGGAATCACTTCGCAATGGCCAAGACAATTGcgatgccgcttacacctgc<br>ctgctcgaggcggccttcaagcaccgcggccgcggccgcgatggttcccatgatctcgg<br>ccgagatcatgggaaccatcgcggccgcggccgcggtgcttgaaggccgcctcgagcag<br>CCACCACCTCCAGAGCCACCGCCACCATCGATgttcttgccccggttgaattgatcgaag |
| CB2003 | Top10/ pUAB200- wag31(Mtb)-EQR210-11-13AAA-dhfr3 | AGGAATCACTTCGCAATGGCCAAGACAATTGcgatgccgcttacacctgc<br>ggtgcggtactcacgttcgaaggtcgccagcgcggcgaggcggccttcaagcaccgcgc<br>gcgcggtgcttgaaggccgcctcgccgcgctggcgaccttcgaacgtgagtaccgcacc<br>CCACCACCTCCAGAGCCACCGCCACCATCGATgttcttgccccggttgaattgatcgaag |
| CB2004 | Top10/ pUAB200- wag31(Mtb)-RRK220,2,4AAA-dhfr3 | AGGAATCACTTCGCAATGGCCAAGACAATTGcgatgccgcttacacctgc<br>ctccagctgcgattccaggtaggtggcgagcgcggtggcgtactcacgttcgaaggtac<br>gtaccttcgaacgtgagtacgccaccgcgctcgccacctacctggaatcgcagctggag<br>GAACTGCCTCCGACTATCCA |
| CB2005 | Top10/ pUAB200- wag31(Mtb)-QEE230,2,3AAA-dhfr3 | AGGAATCACTTCGCAATGGCCAAGACAATTGcgatgccgcttacacctgc<br>cgccgccgatccacgctggccgagcgccgccagggccgattccaggtaggtcttgagcc<br>ggctcaagacctacctggaatcggccctggcggcgctcggccagcgtggatcggcggcg |

| | | |
|---|---|---|
| | | GAACTGCCTCCGACTATCCA |
| CB2006 | Top10/ pUAB200- wag31(Mtb)-D244A-dhfr3 | AGGAATCACTTCGCAATGGCCAAGACAATTGcgatgccgcttacacctgc<br>cgaagccaccggcatccgcattggacgcgaccggcgccgccgatccacgc<br>gcgtggatcggcggcgccggtcgcgtccaatgcggatgccggtggcttcg<br>CCTTTTTCCTCCTGGACCTCAGAGAGGACGCCTGGGTATT |
| CB2007 | Top10/ pUAB200- wag31(Mtb)-N246A-dhfr3 | AGGAATCACTTCGCAATGGCCAAGACAATTGcgatgccgcttacacctgc<br>gaattgatcgaagccaccggcatccgcggcggaatcgaccggcgccgccgatc<br>gatcggcggcgccggtcgattccgccgcggatgccggtggcttcgatcaattc<br>CCTTTTTCCTCCTGGACCTCAGAGAGGACGCCTGGGTATT |
| CB1909 | Top10/ pKK158 - Pwag31-wag31(Mtb)-wt | CTTTTTGCGTTTAATACTGCATGCACTCTAGAcgctgttcatcttctggctgctgctca<br>ggacgtcggcaggtgtaagcggcatATGtgtctgccccttgaagtcttgaaccg<br>ggttcaagacttcaaggggcagacaCATatgccgcttacacctgccgacgtcc<br>GACCTCTAGGGTCCCCAATTAATTAGCTAAAGCTTctagttcttgccccggttgaattg |
| CB1683 | XL1 Blue/ pKK158- Pwag31-wag31(Msmeg) | CTTTTTGCGTTTAATACTGCATGCACTCTAGAcgctgttcatcttctggctgctgctca<br>GACCTCTAGGGTCCCCAATTAATTAGCTAAAGCTTcagttgttgccgcggttgaactg |
| CB1938 | Top10/pkk158-Pwag31--wag31(Mtb)-N10A- | ggttcaagacttcaaggggcagacaCATatgccgcttacacctgccgacgtcc<br>GACCTCTAGGGTCCCCAATTAATTAGCTAAAGCTTctagttcttgccccggttgaattg |
| CB1939 | Top10/pkk158-Pwag31--wag31(Mtb)-S14A- | ggttcaagacttcaaggggcagacaCATatgccgcttacacctgccgacgtcc<br>GACCTCTAGGGTCCCCAATTAATTAGCTAAAGCTTctagttcttgccccggttgaattg |
| CB1941 | Top10/pkk158-Pwag31--wag31(Mtb)-K20A- | CTTTTTGCGTTTAATACTGCATGCACTCTAGAcgctgttcatcttctggctgctgctca<br>ggacgtcggcaggtgtaagcggcatATGtgtctgccccttgaagtcttgaaccg<br>ggttcaagacttcaaggggcagacaCATatgccgcttacacctgccgacgtcc<br>GACCTCTAGGGTCCCCAATTAATTAGCTAAAGCTTctagttcttgccccggttgaattg |

| | | |
|---|---|---|
| CB1942 | Top10/pkk158-Pwag31--wag31(Mtb)-R21A- | CTTTTTGCGTTTAATACTGCATGCACTCTAGAcgctgttcatcttctggctgctgctca<br>ggacgtcggcaggtgtaagcggcatATGtgtctgccccccttgaagtcttgaaccg<br>ggttcaagacttcaaggggggcagacaCATatgccgcttacacctgccgacgtcc<br>GACCTCTAGGGTCCCCAATTAATTAGCTAAAGCTTctagttcttgccccggttgaattg |
| CB1943 | Top10/pkk158-Pwag31--wag31(Mtb)-KR20,21AA- | CTTTTTGCGTTTAATACTGCATGCACTCTAGAcgctgttcatcttctggctgctgctca<br>ggacgtcggcaggtgtaagcggcatATGtgtctgccccccttgaagtcttgaaccg<br>ggttcaagacttcaaggggggcagacaCATatgccgcttacacctgccgacgtcc<br>GACCTCTAGGGTCCCCAATTAATTAGCTAAAGCTTctagttcttgccccggttgaattg |
| CB1944 | Top10/pkk158-Pwag31--wag31(Mtb)-ENE36-38AAA- | CTTTTTGCGTTTAATACTGCATGCACTCTAGAcgctgttcatcttctggctgctgctca<br>ggacgtcggcaggtgtaagcggcatATGtgtctgccccccttgaagtcttgaaccg<br>ggttcaagacttcaaggggggcagacaCATatgccgcttacacctgccgacgtcc<br>GACCTCTAGGGTCCCCAATTAATTAGCTAAAGCTTctagttcttgccccggttgaattg |
| CB1945 | Top10/pkk158-Pwag31--wag31(Mtb)-E45R- | CTTTTTGCGTTTAATACTGCATGCACTCTAGAcgctgttcatcttctggctgctgctca<br>ggacgtcggcaggtgtaagcggcatATGtgtctgccccccttgaagtcttgaaccg<br>ggttcaagacttcaaggggggcagacaCATatgccgcttacacctgccgacgtcc<br>GACCTCTAGGGTCCCCAATTAATTAGCTAAAGCTTctagttcttgccccggttgaattg |
| CB1946 | Top10/pkk158-Pwag31--wag31(Mtb)-E45D- | CTTTTTGCGTTTAATACTGCATGCACTCTAGAcgctgttcatcttctggctgctgctca<br>ggacgtcggcaggtgtaagcggcatATGtgtctgccccccttgaagtcttgaaccg<br>ggttcaagacttcaaggggggcagacaCATatgccgcttacacctgccgacgtcc<br>GACCTCTAGGGTCCCCAATTAATTAGCTAAAGCTTctagttcttgccccggttgaattg |
| CB1947 | Top10/pkk158-Pwag31--wag31(Mtb)-N260A- | CTTTTTGCGTTTAATACTGCATGCACTCTAGAcgctgttcatcttctggctgctgctca<br>ggacgtcggcaggtgtaagcggcatATGtgtctgccccccttgaagtcttgaaccg<br>ggttcaagacttcaaggggggcagacaCATatgccgcttacacctgccgacgtcc |

| | | |
|---|---|---|
| | | GGTCCCCAATTAATTAGCTAAAGCTTctacgccttgcc ccggttgaattgatcgaagcc |
| CB1948 | Top10/pkk158-Pwag31--wag31(Mtb)-K259A- | CTTTTTGCGTTTAATACTGCATGCACTCTAGAcgct gttcatcttctggctgctgctca ggacgtcggcaggtgtaagcggcatATGtgtctgccccttgaagtct tgaaccg ggttcaagacttcaaggggcagacaCATatgccgcttacacctgc cgacgtcc GGTCCCCAATTAATTAGCTAAAGCTTctagttagcgcc ccggttgaattgatcgaag |
| CB1949 | Top10/pkk158-Pwag31-wag31(Mtb)-R257A- | CTTTTTGCGTTTAATACTGCATGCACTCTAGAcgct gttcatcttctggctgctgctca ggacgtcggcaggtgtaagcggcatATGtgtctgccccttgaagtct tgaaccg ggttcaagacttcaaggggcagacaCATatgccgcttacacctgc cgacgtcc CTAGGGTCCCCAATTAATTAGCTAAAGCTTctagttc ttgcccgcgttgaattgatc |
| CB1950 | Top10/pkk158-Pwag31-wag31(Mtb)-N256A- | CTTTTTGCGTTTAATACTGCATGCACTCTAGAcgct gttcatcttctggctgctgctca ggacgtcggcaggtgtaagcggcatATGtgtctgccccttgaagtct tgaaccg ggttcaagacttcaaggggcagacaCATatgccgcttacacctgc cgacgtcc GGGTCCCCAATTAATTAGCTAAAGCTTctagttcttgc cccgagcgaattgatcgaag |
| CB1951 | Top10/pkk158-Pwag31-wag31(Mtb)-F255A- | CTTTTTGCGTTTAATACTGCATGCACTCTAGAcgct gttcatcttctggctgctgctca ggacgtcggcaggtgtaagcggcatATGtgtctgccccttgaagtct tgaaccg ggttcaagacttcaaggggcagacaCATatgccgcttacacctgc cgacgtcc CTAGGGTCCCCAATTAATTAGCTAAAGCTTctagttc ttgccccggttagcttgatcg |
| CB1952 | Top10/pkk158-Pwag31-wag31(Mtb)-Q254A- | CTTTTTGCGTTTAATACTGCATGCACTCTAGAcgct gttcatcttctggctgctgctca ggacgtcggcaggtgtaagcggcatATGtgtctgccccttgaagtct tgaaccg ggttcaagacttcaaggggcagacaCATatgccgcttacacctgc cgacgtcc GGGTCCCCAATTAATTAGCTAAAGCTTctagttcttgc cccggttgaacgcatcgaag |
| CB1953 | Top10/pkk158-Pwag31-wag31(Mtb)-D253A- | CTTTTTGCGTTTAATACTGCATGCACTCTAGAcgct gttcatcttctggctgctgctca ggacgtcggcaggtgtaagcggcatATGtgtctgccccttgaagtct tgaaccg |

| | | |
|---|---|---|
| | | ggttcaagacttcaagggggcagacaCATatgccgcttacacctgc cgacgtcc GGGTCCCCAATTAATTAGCTAAAGCTTctagttcttgc cccggttgaattgagcgaagc |
| CB1954 | Top10/pkk158-Pwag31-wag31(Mtb)-F252A- | CTTTTTGCGTTTAATACTGCATGCACTCTAGAcgct gttcatcttctggctgctgctca ggacgtcggcaggtgtaagcggcatATGtgtctgccccttgaagtct tgaaccg ggttcaagacttcaagggggcagacaCATatgccgcttacacctgc cgacgtcc GGTCCCCAATTAATTAGCTAAAGCTTctagttcttgccc cggttgaattgatctgcgcc |
| CB1955 | Top10/pkk158-Pwag31-wag31(Mtb)-D248A- | CTTTTTGCGTTTAATACTGCATGCACTCTAGAcgct gttcatcttctggctgctgctca ggacgtcggcaggtgtaagcggcatATGtgtctgccccttgaagtct tgaaccg ggttcaagacttcaagggggcagacaCATatgccgcttacacctgc cgacgtcc CTAGGGTCCCCAATTAATTAGCTAAAGCTTctagttc ttgccccggttgaattgatcg |
| CB1979 | Top10/pkk158-Pwag31-wag31(Mtb) -RGKN257-60AAAA | CTTTTTGCGTTTAATACTGCATGCACTCTAGAcgct gttcatcttctggctgctgctca ggacgtcggcaggtgtaagcggcatATGtgtctgccccttgaagtct tgaaccg ggttcaagacttcaagggggcagacaCATatgccgcttacacctgc cgacgtcc GGGTCCCCAATTAATTAGCTAAAGCTTctacgcagct gccgcgttgaattgatcgaag |
| CB1980 | Top10/pkk158-Pwag31-wag31(Mtb)- DQFN253-6AAAA | CTTTTTGCGTTTAATACTGCATGCACTCTAGAcgct gttcatcttctggctgctgctca ggacgtcggcaggtgtaagcggcatATGtgtctgccccttgaagtct tgaaccg ggttcaagacttcaagggggcagacaCATatgccgcttacacctgc cgacgtcc GGTCCCCAATTAATTAGCTAAAGCTTctagttcttgccc cgagctgcagctgcgaagcc |
| CB1981 | Top10/pkk158-Pwag31-wag31(Mtb)-GGF250-2AAA | CTTTTTGCGTTTAATACTGCATGCACTCTAGAcgct gttcatcttctggctgctgctca ggacgtcggcaggtgtaagcggcatATGtgtctgccccttgaagtct tgaaccg ggttcaagacttcaagggggcagacaCATatgccgcttacacctgc cgacgtcc CTAGGGTCCCCAATTAATTAGCTAAAGCTTctagttc ttgccccggttgaattgatctg |
| CB1982 | Top10/pkk158-Pwag31-wag31(Mtb)- PLTP2-5AAAA | CTTTTTGCGTTTAATACTGCATGCACTCTAGAcgct gttcatcttctggctgctgctca |

| | | |
|---|---|---|
| | | GACCTCTAGGGTCCCCAATTAATTAGCTAAAGCT Tctagttcttgccccggttgaattg gttcaagacttcaagggggcagacaCATatggctgccgcagctgcc gacgtccacaatg cattgtggacgtcggcagctgcggcagccatATGtgtctgcccccttg aagtcttgaac |
| CB1983 | Top10/pkk158-Pwag31-wag31(Mtb)- DVH7-9AAA | CTTTTTGCGTTTAATACTGCATGCACTCTAGAcgct gttcatcttctggctgctgctca GACCTCTAGGGTCCCCAATTAATTAGCTAAAGCT Tctagttcttgccccggttgaattg gttcaagacttcaagggggcagacaCATatgccgcttacacctgcc gctgcggctaatg cattagccgcagcggcaggtgtaagcggcatATGtgtctgcccccttt gaagtcttgaac |
| CB1984 | Top10/pkk158-Pwag31-wag31(Mtb)- ΔCt 10aa | CTTTTTGCGTTTAATACTGCATGCACTCTAGAcgct gttcatcttctggctgctgctca ggttcaagacttcaagggggcagacaCATatgccgcttacacctgc cgacgtcc CTAGGGTCCCCAATTAATTAGCTAAAGCTTctaacc ggcatccgcattggaatc |
| CB2047 | Top10/pkk158-Pwag31-wag31(Mtb)-DQE57-9AAA | CTTTTTGCGTTTAATACTGCATGCACTCTAGAcgct gttcatcttctggctgctgctca GACCTCTAGGGTCCCCAATTAATTAGCTAAAGCT Tctagttcttgccccggttgaattg |
| CB2048 | Top10/pkk158-Pwag31-wag31(Mtb)-NEEQ 96-9AAAA | CTTTTTGCGTTTAATACTGCATGCACTCTAGAcgct gttcatcttctggctgctgctca GACCTCTAGGGTCCCCAATTAATTAGCTAAAGCT Tctagttcttgccccggttgaattg |
| CB2049 | Top10/pkk158-Pwag31-wag31(Mtb)-ESDK 124-127AAAA | CTTTTTGCGTTTAATACTGCATGCACTCTAGAcgct gttcatcttctggctgctgctca GACCTCTAGGGTCCCCAATTAATTAGCTAAAGCT Tctagttcttgccccggttgaattg |
| CB2050 | Top10/pkk158-Pwag31-wag31(Mtb) -ERH 141,143,144AAA | CTTTTTGCGTTTAATACTGCATGCACTCTAGAcgct gttcatcttctggctgctgctca GACCTCTAGGGTCCCCAATTAATTAGCTAAAGCT Tctagttcttgccccggttgaattg |
| CB2153 | Top10 / pkk158-Pwag31-wag31-E27A- | CTTTTTGCGTTTAATACTGCATGCACTCTAGAcgct gttcatcttctggctgctgctca GACCTCTAGGGTCCCCAATTAATTAGCTAAAGCT Tctagttcttgccccggttgaattg |
| CB2154 | Top10 / pkk158-Pwag31-wag31(Mtb)-E38A | CTTTTTGCGTTTAATACTGCATGCACTCTAGAcgct gttcatcttctggctgctgctca GACCTCTAGGGTCCCCAATTAATTAGCTAAAGCT Tctagttcttgccccggttgaattg |
| CB2155 | Top10 / pkk158-Pwag31-wag31(Mtb)-L34A | CTTTTTGCGTTTAATACTGCATGCACTCTAGAcgct gttcatcttctggctgctgctca |

| | | |
|---|---|---|
| | | GACCTCTAGGGTCCCCAATTAATTAGCTAAAGCT Tctagttcttgccccggttgaattg |
| CB2157 | Top10 / pkk158-Pwag31-wag31(Mtb)-NSD46-8-AAA | CTTTTTGCGTTTAATACTGCATGCACTCTAGAcgct gttcatcttctggctgctgctca GACCTCTAGGGTCCCCAATTAATTAGCTAAAGCT Tctagttcttgccccggttgaattg |
| CB2159 | Top10 / pkk158-Pwag31-wag31(Mtb)-ER153,155AA | CTTTTTGCGTTTAATACTGCATGCACTCTAGAcgct gttcatcttctggctgctgctca GACCTCTAGGGTCCCCAATTAATTAGCTAAAGCT Tctagttcttgccccggttgaattg |
| CB2160 | Top10 / pkk158-Pwag31-wag31(Mtb)-QR156,157AA | CTTTTTGCGTTTAATACTGCATGCACTCTAGAcgct gttcatcttctggctgctgctca GACCTCTAGGGTCCCCAATTAATTAGCTAAAGCT Tctagttcttgccccggttgaattg |
| CB2161 | Top10 / pkk158-Pwag31-wag31(Mtb)-QSRS166-9AAAA | CTTTTTGCGTTTAATACTGCATGCACTCTAGAcgct gttcatcttctggctgctgctca GACCTCTAGGGTCCCCAATTAATTAGCTAAAGCT Tctagttcttgccccggttgaattg |
| CB2162 | Top10 / pkk158-Pwag31-wag31(Mtb)-RQQ174,5,7AAA | CTTTTTGCGTTTAATACTGCATGCACTCTAGAcgct gttcatcttctggctgctgctca GACCTCTAGGGTCCCCAATTAATTAGCTAAAGCT Tctagttcttgccccggttgaattg |
| CB2163 | Top10 / pkk158-Pwag31-wag31(Mtb)-DER185,7,8AAA | CTTTTTGCGTTTAATACTGCATGCACTCTAGAcgct gttcatcttctggctgctgctca GACCTCTAGGGTCCCCAATTAATTAGCTAAAGCT Tctagttcttgccccggttgaattg |
| CB2164 | Top10 / pkk158-Pwag31-wag31(Mtb)-NQQR199-202AAAA | CTTTTTGCGTTTAATACTGCATGCACTCTAGAcgct gttcatcttctggctgctgctca GACCTCTAGGGTCCCCAATTAATTAGCTAAAGCT Tctagttcttgccccggttgaattg |
| CB2167 | Top10 / pkk158-Pwag31-wag31(Mtb)-QEE230,2,3AAA | CTTTTTGCGTTTAATACTGCATGCACTCTAGAcgct gttcatcttctggctgctgctca GACCTCTAGGGTCCCCAATTAATTAGCTAAAGCT Tctagttcttgccccggttgaattg |
| CB2168 | Top10 / pkk158-Pwag31-wag31(Mtb)-D244A | CTTTTTGCGTTTAATACTGCATGCACTCTAGAcgct gttcatcttctggctgctgctca GACCTCTAGGGTCCCCAATTAATTAGCTAAAGCT Tctagttcttgccccggttgaattg |
| CB2169 | Top10 / pkk158-Pwag31-wag31(Mtb)-N246A | CTTTTTGCGTTTAATACTGCATGCACTCTAGAcgct gttcatcttctggctgctgctca GACCTCTAGGGTCCCCAATTAATTAGCTAAAGCT Tctagttcttgccccggttgaattg |
| CB2222 | Top10 / pkk158-Pwag31-wag31(Mtb)-P2A | CTTTTTGCGTTTAATACTGCATGCACTCTAGAcgct gttcatcttctggctgctgctca |

| | | |
|---|---|---|
| | | CattgtggacgtcggcaggtgtaagggccatATGtgtctgccccctg aagtcttgaac<br>gttcaagacttcaagggggcagacaCATatggcccttacacctgcc gacgtccac<br>GACCTCTAGGGTCCCCAATTAATTAGCTAAAGCT Tctagttcttgccccggttgaattg |
| CB2223 | Top10 / pkk158-Pwag31-wag31(Mtb)-T4A | CTTTTTGCGTTTAATACTGCATGCACTCTAGAcgct gttcatcttctggctgctgctca<br>ccacattgtggacgtcggcaggcgcaagcggcatATGtgtctgcccc cttgaagtcttg<br>caagacttcaagggggcagacaCATatgccgcttgcgcctgccgac gtccacaatgtgg<br>GACCTCTAGGGTCCCCAATTAATTAGCTAAAGCT Tctagttcttgccccggttgaattg |
| CB2224 | Top10 / pkk158-Pwag31-wag31(Mtb)-P5A | CTTTTTGCGTTTAATACTGCATGCACTCTAGAcgct gttcatcttctggctgctgctca<br>cttactgaacgccacattgtggacgtcggccgctgtaagcggcatAT Gtgtctgccccc<br>gggggcagacaCATatgccgcttacagcggccgacgtccacaatg tggcgttcagtaag<br>GACCTCTAGGGTCCCCAATTAATTAGCTAAAGCT Tctagttcttgccccggttgaattg |
| CB2225 | Top10 / pkk158-Pwag31-wag31(Mtb)-D7A | CTTTTTGCGTTTAATACTGCATGCACTCTAGAcgct gttcatcttctggctgctgctca<br>cttactgaacgccacattgtggaccgcggcaggtgtaagcggcatAT Gtgtctgccccc<br>gggggcagacaCATatgccgcttacacctgccgcggtccacaatgt ggcgttcagtaag<br>GACCTCTAGGGTCCCCAATTAATTAGCTAAAGCT Tctagttcttgccccggttgaattg |
| CB2226 | Top10 / pkk158-Pwag31-wag31(Mtb)-H9A | CTTTTTGCGTTTAATACTGCATGCACTCTAGAcgct gttcatcttctggctgctgctca<br>cggcttactgaacgccacattcgcgacgtcggcaggtgtaagcggcat ATGtgtctgcc<br>ggcagacaCATatgccgcttacacctgccgacgtcgcgaatgtggc gttcagtaagccg<br>GACCTCTAGGGTCCCCAATTAATTAGCTAAAGCT Tctagttcttgccccggttgaattg |
| CB2227 | Top10/ pkk158-Pwag31-wag31(Mtb)-wt-GFP | CTTTTTGCGTTTAATACTGCATGCACTCTAGAcgct gttcatcttctggctgctgctca<br>CAGTGAAAAGTTCTTCTCCTTTACTtccggagttcttgcc ccggttgaattgatcg<br>cgatcaattcaaccggggcaagaactccggaAGTAAAGGAGA AGAACTTTTCACTG<br>GACCTCTAGGGTCCCCAATTAATTAGCTAAAGCT TctaTTTGTATAGTTCATCCATGCCA |

| | | |
|---|---|---|
| CB2279 | Top10/ pkk158-Pwag31-wag31(Mtb)-NQQR199-202AAAA-GFP | CTTTTTGCGTTTAATACTGCATGCACTCTAGAcgctgttcatcttctggctgctgctca<br>CAGTGAAAAGTTCTTCTCCTTTACTtccggagttcttgccccggttgaattgatcg<br>cgatcaattcaaccggggcaagaactccggaAGTAAAGGAGAAGAACTTTTCACTG<br>GACCTCTAGGGTCCCCAATTAATTAGCTAAAGCTTctaTTTGTATAGTTCATCCATGCCA |
| CB2284 | Top10/ pkk158-Pwag31-wag31(Mtb)-L34A-GFP | CTTTTTGCGTTTAATACTGCATGCACTCTAGAcgctgttcatcttctggctgctgctca<br>CAGTGAAAAGTTCTTCTCCTTTACTtccggagttcttgccccggttgaattgatcg<br>cgatcaattcaaccggggcaagaactccggaAGTAAAGGAGAAGAACTTTTCACTG<br>GACCTCTAGGGTCCCCAATTAATTAGCTAAAGCTTctaTTTGTATAGTTCATCCATGCCA |
| CB2319 | Top10/ pGtetO-GlfT2-mRFP-ZeoR | GCTTAATTAAGAAGGAGATGAATTCatgagtgacatcccttccggcgcactcgaagccg<br>gagacaccggagaaagtcggacgagagctcATGGCCTCCTCCGAGGACGtcatcaaggag<br>ctccttgatgaCGTCCTCGGAGGAGGCCATgagctctcgtccgactttctccggtgtctcggtgag<br>GTGCAGGACCATGTGGTCCCGAGCGCTTCACTTCTCGAACTGGGGGTGGCTCCAGTCGG |
| CB2320 | Top 10 / pGtetO - Wag31-GFP11 | TTACGCCAAGCTCTAATACGACTCACTATAGGGAAGCTGCAAGGCGATTAAGTTGGGTA<br>CGATCCAATATTGTTAACTACGTCGACATCGATAAACAGCTATGACCATGATTACGCCA<br>ATGCAAGCTTGGCGTAATCATGGTCATAGCTGTTTATCGATGTCGACGTAGTTAACAAT<br>atGAATTCATCTCCTTCTTAATTAAGCATGCGGATCGTGCTCATTTCGGG<br>CTTAATTAAGAAGGAGATGAATTCatgccgctcacaccagcggacgtcca<br>ACGTATTCGTGCAGGACCATGTGGTCCCGAGCGCTGCCgttgttgccgcggttgaactg<br>CCAATTAATTAGCTAACTAGGTGATCCCCGCGGCGTTCACGTATTCGTGCAGGACCATG<br>GGAAAATTTAAAATAAAAAAGGGGACCTCTAGGGTCCCCAATTAATTAGCTAACTAGGT |
| CB2326 | Top10 / pkk158-Pwag31-wag31(Mtb)-D7A-GFP | CTTTTTGCGTTTAATACTGCATGCACTCTAGAcgctgttcatcttctggctgctgctca<br>CAGTGAAAAGTTCTTCTCCTTTACTtccggagttcttgccccggttgaattgatcg<br>cgatcaattcaaccggggcaagaactccggaAGTAAAGGAGAAGAACTTTTCACTG |

| | | |
|---|---|---|
| | | GACCTCTAGGGTCCCCAATTAATTAGCTAAAGCT<br>TctaTTTGTATAGTTCATCCATGCCA |
| CB2327 | Top10/ pGtetO-mcherry2b-GFP11 | TGAGCACGATCCGCATGCTTAATTAAGAAGGAG<br>ATGAATTCATGGATAGCACTGAGAGC<br>TCACGTATTCGTGCAGGACCATGTGGTCCCGAG<br>CGCTTCTGGATCCGCTAGATCCCTGG |
| CB2328 | Top10/ pGtetO-GFP11-mcherry | TGCACGAATACGTGAACGCCGCGGGGATCACC<br>GGGCCCGATAGCACTGAGAGCGGCTCC<br>ATTAAGAAGGAGATGAATTCATGCGGGACCACA<br>TGGTCCTGCACGAATACGTGAACGCC<br>ATTCGCCGCCCGAAATGAGCACGATCCGCATGC<br>TTAATTAAGAAGGAGATGAATTCATG<br>ATTCGTGCAGGACCATGTGGTCCCGAGCGCTCT<br>ATTATCTGGATCCGCTAGATCCCTGG |
| CB2329 | Top10/ pGtetO-GFP11-Wag31 | GCACGAATACGTGAACGCCGCGGGGATCACCG<br>GGCCCccgctcacaccagcggacgtcc<br>ACGTATTCGTGCAGGACCATGTGGTCCCGAGCG<br>CTTCAgttgttgccgcggttgaactg |
| CB2330 | Top10/ CT94-GFP1-10-Wag31 | GATCCGCATGCTTAATTAAGAAGGAGATATACAT<br>atgGGCGGGACCTCGATGTCGAAGGG<br>ttatggacgtccgctggtgtgagcggACCAGCGCTTCCCTTT<br>TCGTTCGGGTCCTTGGA<br>TGCTCTCCAAGGACCCGAACGAAAAGGGAAGCG<br>CTGGTccgctcacaccagcggacgtc<br>GACCTCTAGGGTCCCCAATTAATTAGCTAAAGCT<br>TCTAgttgttgccgcggttgaactg |
| CB2340 | Top10/ pkk158-Pwag31-wag31(Mtb)-K20A-GFP | CTTTTTGCGTTTAATACTGCATGCACTCTAGAcgct<br>gttcatcttctggctgctgctca<br>CAGTGAAAAGTTCTTCTCCTTTACTtccggagttcttgcc<br>ccggttgaattgatcg<br>cgatcaattcaaccggggcaagaactccggaAGTAAAGGAGA<br>AGAACTTTTCACTG<br>GACCTCTAGGGTCCCCAATTAATTAGCTAAAGCT<br>TctaTTTGTATAGTTCATCCATGCCA |
| CB2341 | Top10/ pkk158-Pwag31-wag31(Mtb)-F255A-GFP | CTTTTTGCGTTTAATACTGCATGCACTCTAGAcgct<br>gttcatcttctggctgctgctca<br>CAGTGAAAAGTTCTTCTCCTTTACTtccggagttcttgcc<br>ccggttagcttgatcgaag<br>GACCTCTAGGGTCCCCAATTAATTAGCTAAAGCT<br>TctaTTTGTATAGTTCATCCATGCCA |
| CB2393 | Top10/ pGtetO-wga31--D7A-GFP11 | CATGCTTAATTAAGAAGGAGATGAATTCatgccgctt<br>acacctgccgcggtccac<br>CGTGCAGGACCATGTGGTCCCGAGCGCTGCCgtt<br>cttgccccggttgaattgatc |
| CB2394 | Top10/ pGtetO-wga31--K20A-GFP11 | CTTAATTAAGAAGGAGATGAATTCatgccgcttacacct<br>gccgacgtccacaatg |

| | | |
|---|---|---|
| | | CGTGCAGGACCATGTGGTCCCGAGCGCTGCCgtt cttgccccggttgaattgatc |
| CB2395 | Top10/ pGtetO-wga31--L34A-GFP11 | CTTAATTAAGAAGGAGATGAATTCatgccgcttacacct gccgacgtccacaatg CGTGCAGGACCATGTGGTCCCGAGCGCTGCCgtt cttgccccggttgaattgatc |
| CB2396 | Top10/ pGtetO-wga31--NQQR199-202AAAA-GFP11 | CTTAATTAAGAAGGAGATGAATTCatgccgcttacacct gccgacgtccacaatg CGTGCAGGACCATGTGGTCCCGAGCGCTGCCgtt cttgccccggttgaattgatc |
| CB2397 | Top10/ pGtetO-wga31--F255A-GFP11 | CTTAATTAAGAAGGAGATGAATTCatgccgcttacacct gccgacgtccacaatg CGTGCAGGACCATGTGGTCCCGAGCGCTGCCgtt cttgccccggttagcttgatc |
